# The β-amyloid oligomer Aβ*56 is associated with Alzheimer’s dementia independently of amyloid pathology

**DOI:** 10.1101/2025.04.11.648457

**Authors:** Ian P. Lapcinski, Ashley J. Petersen, Karen H. Ashe, Peng Liu

**Author notes:** Corresponding author: Peng Liu.

## Abstract

The amyloid cascade hypothesis posits that β-amyloid (Aβ) oligomers (Aβo) play a key role in the pathogenesis of Alzheimer’s disease (AD). Aβo are believed to contribute to dementia in AD, and more than a dozen have been identified. While some oligomers are associated with amyloid plaques, others—such as Aβ*56—are not. It remains unclear whether these plaque-independent oligomers contribute to dementia. Aβ*56 is a non-fibrillar, sodium dodecyl sulfate-stable, brain-derived type of Aβo that impairs memory in mice. We recently confirmed and extended the characterization of Aβ*56 in AD mouse models and identified two variants, namely Aβ(40)*56 and Aβ(42)*56 containing canonical Aβ(1-40) and Aβ(1-42), respectively. To examine the role of Aβ*56 in dementia, we measured Aβ*56 in elderly individuals—selected for equivalent levels of amyloid plaques—with no cognitive impairment (NCI), mild cognitive impairment (MCI), and AD dementia. Using immunoprecipitation (IP), western blotting (WB), size exclusion chromatography, and conformation-sensitive antibodies, we confirmed the presence of Aβ(40)*56 and Aβ(42)*56 in human brains. Using IP/WB coupled with densitometry-based semi-quantitative analysis, we found that levels of both Aβ*56 variants were elevated in AD dementia compared to NCI and MCI and correlated inversely with various measures of memory and cognition. Furthermore, mediation analyses showed that these associations were mediated by phosphorylated tau at serine 202 and threonine 205 (ptau 202/205)-containing neurofibrillary tangles. These findings support an association between Aβ*56 and AD dementia independently of amyloid pathology. The results are significant because they suggest that targeting Aβ*56 in patients could provide additional cognitive benefits beyond those achieved by current anti-Aβ therapies that focus on removing amyloid plaques.

## Introduction

A current hypothesis of the etiology of Alzheimer’s disease (AD) suggests that soluble β-amyloid (Aβ) oligomers (Aβo) are an essential contributor to disease pathogenesis [15]. Among the numerous Aβo discovered in the brain [2, 5, 19, 25, 31, 40, 41, 46, 49, 51, 52, 78, 82, 83, 85, 91], Aβ*56 is a ∼56-kilo-Dalton (kDa), non-fibrillar, sodium dodecyl sulfate (SDS)-stable, water-soluble oligomer that has been shown to be associated with aging [53, 73, 102] and cognitive dysfunction [8, 12, 13, 23, 51, 52, 55, 67, 97, 103] in mice [8, 12, 13, 23, 51, 52, 55, 67, 97, 102] and dogs [73]. Aβ*56 binds anti-oligomer A11 antibodies [57, 73, 103], a conformation-sensitive agent [41] that recognizes non-fibrillar structures, such as cylindrically shaped β-barrels [48] and out-of-register antiparallel β-sheets [54], distinct from the in-register parallel β-sheets in brain-derived amyloid fibrils and dense-core neuritic plaques [43, 59, 95, 101]. In Tg2576 mice expressing transgenic human amyloid precursor protein (hAPP) with the Swedish (K670N, M671L) mutation linked to familial AD [14], Aβ*56 appears before amyloid plaques [11, 38, 57], suggesting that the biogenesis of Aβ*56 occurs independently of amyloid fibrils and plaques. Aβ*56 from Tg2576 mice [33] impairs memory when injected intrahippocampally into young, healthy mice [57], confirming previous results in rats [51, 52]. We recently refined methods for detecting Aβ*56 to ensure that the ∼56-kDa entity measured contains canonical Aβ(1-40) or Aβ(1-42) [57], which we named Aβ(40)*56 and Aβ(42)*56, respectively. We found both Aβ(40)*56 and Aβ(42)*56 in various AD mouse models expressing transgenic hAPP mutants [57].

While there is general agreement about the association between Aβ*56 and cognitive dysfunction in mice, there are conflicting results concerning the relevance of Aβ*56 to AD dementia in humans. One study showed that its levels are higher in cognitively impaired individuals with probable AD dementia [103], while another showed that they are lower [53]. Moreover, whether Aβ*56 is associated with memory loss and cognitive decline independently of amyloid plaques is unclear.

In this study, we aimed to reconcile the discrepant results and to understand the role of the plaque-independent oligomer Aβ*56 in AD dementia. To accomplish this, we selected individuals with different cognitive statuses but similar plaque loads. Specifically, we chose individuals with no cognitive impairment (NCI), mild cognitive impairment (MCI), and AD dementia with comparable amyloid plaque loads—determined by immunohistology—in the inferior temporal gyrus (ITG) and measured Aβ(40)*56 and Aβ(42)*56 in that region. Levels of both Aβ(40)*56 and Aβ(42)*56 were significantly higher in AD dementia relative to NCI and MCI. When adjusted for plaque loads, Aβ(40)*56 remained significantly elevated, while Aβ(42)*56 showed a strong trend. In addition, both Aβ*56 variants correlated inversely with performances in tests of several types of memory and cognition. Furthermore, mediation analyses showed that the association between Aβ*56 and cognitive and memory impairment was mediated strongly by phosphorylated tau at serine 202 and threonine 205 (ptau 202/205)-containing neurofibrillary tangles (NFT), but minimally by Aβ plaques, suggesting that the cognitive effects of Aβ*56 depend on pathological tau. Taken altogether, the findings are important because they indicate that patients receiving anti-protofibril [19] and anti-fibril [18, 87] antibodies, which focus on removing plaques, could receive additional cognitive benefit by targeting Aβ*56.

## Materials and methods

### Human brain collection and sample size calculations

Post-mortem frozen ITG (Brodmann area 20) specimens were obtained from the Religious Orders Study (ROS) and the Rush Memory Aging Project (MAP) [7], Rush University, Chicago, Illinois. To estimate the appropriate sample size for the study, we measured levels of Aβ(40)*56 and Aβ(42)*56 using IP/WB (Fig. **1**a and c) in a small cohort containing individuals with AD dementia (*N* = 15) and age-matched individuals with NCI (*N* = 11) with comparable immunohistologically determined Aβ plaque loads in the ITG. We showed that levels of Aβ(40)*56 and Aβ(42)*56 that were normalized to levels of neuronal nuclei (NeuN)—a protein marker for post-mitotic neurons, the principal cell type in which Aβ production occurs via hAPP processing [70, 93]—in AD dementia were 4.6- and 3.6-fold higher (in both cases, *U* = 38; *P* = 0.02; two-tailed, unpaired Mann-Whitney *U* tests), respectively, than in NCI (Online Resource **1**: Figure S1). Using densitometry-based semi-quantitative analyses, we determined that the lowest sample size required for each group to achieve statistical significance under type I error *α* = 0.01 and power (*i.e.*, (1-*β*)) = 0.80 is *N* = 19.

**Fig. 1.**
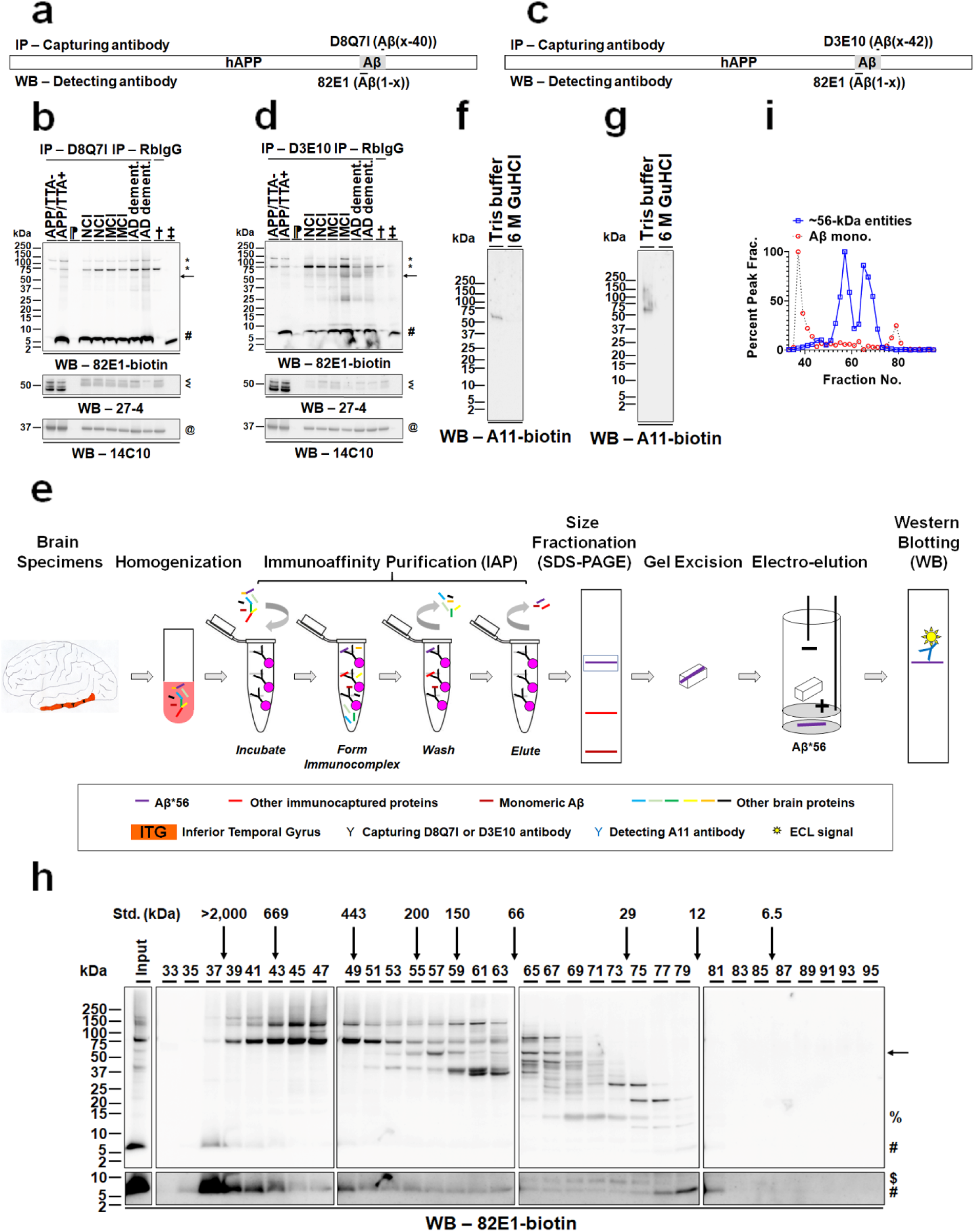
Aβ*56 variants are water-soluble, SDS-stable Aβo present in the inferior temporal gyri of human individuals. (**a, c**) Schematic diagrams illustrating IP/WB settings for detecting Aβ(40)*56 (a) and Aβ(42)*56 (c). hAPP, human amyloid precursor protein; Aβ, β-amyloid. (**b, d**) *Upper*, representative IP/WB showing that D8Q71 (b)- or D3E10 (d)-precipitated proteins from aqueous brain extracts contain ∼56-kDa entities (arrow) and monomeric Aβ (#) that are detected by biotinylated 82E1 (82E1-biotin); *middle*, WB showing NeuN proteins (arrowheads); *bottom*, WB showing GAPDH proteins (@). NCI, no cognitive impairment; MCI, mild cognitive impairment; AD, Alzheimer’s disease dementia. + and -, IP reactions containing a capture antibody and aqueous brain extracts of APP/TTA mice (+) or littermates expressing no hAPP (-) (Online Resource **1**: Table S4). ⁋, IP reactions containing only the capture antibodies. †, IP reactions containing a generic rabbit IgG and aqueous brain extracts (combined from the six individuals shown in b or d, each contributing equivalent protein masses). ‡, synthetic Aβ(1-40) (b) or Aβ(1-42) (d). (**e**) A schematic illustration of isolating Aβ*56 variants from human brains. (**f, g**) Representative IP/WB showing that the detection of D8Q7I (f)- or D3E10 (g)-purified ∼56-kDa entities by anti-oligomer A11 antibodies following treatment with Tris buffer but not 6 M GuHCl. (**h**) Representative WB showing that ∼56-kDa entities (arrow, fractions [Frac.] 51-61 and 63-71), ∼14-kDa entities (%, Frac. 65-79), ∼9-kDa entities ($, Frac. 67-77), and monomeric Aβ (#, Frac. 35-45 and 75-83) are detected in SEC fractions of aqueous AD brain extracts using 82E1-biotin. Lower panels, longer exposure. Std., biomolecule standards. (**i**) Quantification of levels of the ∼56-kDa entities and monomeric Aβ shown in (h). The level of the ∼56-kDa entities in each fraction is normalized to the highest level in fraction 57 and of monomeric Aβ (Aβ mono.), fraction 37. kDa, kilo-Dalton. *, non-specific bands.

To analyze Aβ(40)*56, we selected brain specimens from 61 de-identified individuals, including 22 individuals with NCI, 19 individuals with MCI, and 20 individuals with AD dementia (for detailed demographic and pathological characteristics, see Online Resource **1**: Table S1). To analyze Aβ(42)*56, we selected brain specimens from 62 de-identified individuals, including 23 individuals with NCI, 19 individuals with MCI, and 20 individuals with AD dementia (for detailed demographic and pathological characteristics, see Online Resource **1**: Table S2).

Frozen tissue was stored at −80°C in polypropylene bags prior to shipment to the University of Minnesota, Twin Cities, Minnesota.

All procedures were approved by the Institutional Review Boards of the Rush University and the University of Minnesota.

### Brain protein extraction

To detect and quantify Aβ entities, aqueous brain protein extracts were prepared using a protocol adapted from a previous publication [84]. Specifically, we dissected 0.1-0.8 g of post-mortem frozen tissues from the ITG of de-identified individuals, the hemi-forebrains of 10-11-month-old APP/TTA mice [35], or the hemi-forebrains of age-matched littermates of APP/TTA mice that did not express hAPP, transferred them to 1 mL of ice-cold extraction buffer (25 mM tris(hydroxymethyl)aminomethane (Tris; MilliporeSigma, Burlington, MA; catalog number (Cat. No.): T1378-100G)-hydrochloric acid (HCl; MilliporeSigma, Cat. No.: 320331-500ML), pH 7.4; 140 mM sodium chloride (NaCl; MilliporeSigma, Cat. No.: S9888-5KG); and 3 mM potassium chloride (KCl; MilliporeSigma, Cat. No.: P3911-25G) with the following protease and phosphatase inhibitors: 0.1 mM phenylmethylsulfonyl fluoride (PMSF; MilliporeSigma, Cat. No.: P7626), 0.2 mM 1,10-phenanthroline monohydrate (phen; MilliporeSigma, Cat. No.: P9375), 1% (volume/volume (v/v)) protease inhibitor cocktail (MilliporeSigma, Cat No.: P8340), 1% (v/v) phosphatase inhibitor cocktail 1 (MilliporeSigma, Cat. No.: P2850), and 1% (v/v) phosphatase inhibitor cocktail 2 (MilliporeSigma, Cat. No.: P5726)), and homogenized them using a Dounce homogenizer (PolyScience Corporation, Niles, IL; Cat. No.: 4-741170) at room temperature, full-speed for 10 strokes. We centrifuged the homogenates at 4°C, 16,100 *g* for 1 hr. We then depleted immunoglobulin G (IgG) from the supernatants using Protein G Sepharose 4 Fast Flow resin (GE Healthcare, Piscataway, NJ; Cat. No.: 17-0618-02) and determined protein concentrations of IgG-depleted brain extracts using a bicinchoninic acid (BCA) protein assay kit (Thermo Fisher Scientific, Rockford, IL; Cat. No.: 23225) according to the manufacturer’s instructions. We stored the final aqueous brain protein extracts at −80°C until they were ready to use.

To measure levels of NeuN and the housekeeping protein glyceraldehyde 3-phosphate dehydrogenase (GAPDH), the brain pellets that were recovered from the water-soluble extracts described above were subjected to further processing. For this, we resuspended the pellet from each human specimen in 1 mL of detergent-containing extraction buffer (25 mM Tris-HCl, pH 7.4; 140 mM NaCl; 3 mM KCl; 1% (v/v) polyethylene glycol p-(1,1,3,3-tetramethylbutyl)-phenyl ether (Triton X-100; MilliporeSigma, Cat. No.: T8787); 0.1 mM PMSF; 0.2 mM phen; 1% (v/v) protease inhibitor cocktail; 1% (v/v) phosphatase inhibitor cocktail 1; and 1% (v/v) phosphatase inhibitor cocktail 2) at room temperature and then incubated the resuspended material at 4°C by gently rotating at 15 rpm for 1 hr. Next, we homogenized the resulting material using a Dounce homogenizer at room temperature, full-speed for 25 strokes, and then centrifuged them at 4°C, 16,100 *g* for 90 min. We depleted IgG of the supernatant and determined protein concentrations as described above. We stored the final extracts at −80°C. Similarly, we prepared brain extracts from 4-5-month-old Tg2576 mice modeling AD. We used detergent-containing extracts of Tg2576 [33] mouse brains as an internal control for densitometry-based semiquantitative analysis.

All experiments involving mice were performed in full accordance with the guidelines of the Association for Assessment and Accreditation of Laboratory Animal Care and approved (approval #1202A09927) by the Institutional Animal Care and Use Committee at the University of Minnesota, Twin Cities, Minnesota.

### Size exclusion chromatography (SEC)

To fractionate Aβ-containing entities in the aqueous extracts of human brain tissue by size under non-denaturing conditions, we performed SEC using a Superdex 200 10/300 GL column (GE healthcare, Cat. No.: 17-5175-01) driven by the BioLogic DuoFlow Chromatography system (Bio-Rad, Hercules, CA; research resource identifier (RRID): SCR_019686). To calibrate the column, we determined the elution profiles of a set of eight globular proteins at the molecular weights of 669 (bovine thyroglobulin; MilliporeSigma, Cat. No.: T9145), 443 (apoferritin from horse spleen; MilliporeSigma, Cat. No.: A3660), 200 (β-amylase from sweet potato; MilliporeSigma, Cat. No.: A8781), 150 (alcohol dehydrogenase from yeast; MilliporeSigma, Cat. No.: A8656), 66 (bovine serum albumin; MilliporeSigma, Cat. No.: A8531), 29 (carbonic anhydrase from bovine erythrocytes; MilliporeSigma, Cat. No.: C7025), 12.4 (cytochrome c from horse heart; MilliporeSigma, Cat. No.: C7150), and 6.5 (aprotinin from bovine lung; MilliporeSigma, Cat. No.: A3886) kDa, and Blue Dextran, ∼2,000 kDa (MilliporeSigma, Cat. No.: D4772). We separately injected 250 µL of phosphate-buffered saline (PBS, pH 7.4; MilliporeSigma, Cat. No.: P4417-100TAB) containing 200 µg of each type of biomolecules. We eluted the biomolecule standards at room temperature using PBS with a flow rate of 0.5 mL/min and collected 250-µL fractions. We determined protein concentrations of eluted fractions by the BCA protein assay.

To analyze biological samples, we pooled detergent-free, aqueous brain protein extracts from a total of 10 individuals with AD dementia (Online Resource **1**: Table S1 and S3), each contributing 10% protein mass. We injected 250 µL of the prepared sample containing 2.8 mg of total proteins. We eluted proteins in PBS and determined protein concentrations in the collected fractions as described above. We added 10 µL of inhibitors (2.6 mM PMSF, 5.2 mM phen, 2.6% (v/v) protease inhibitor cocktail, 2.6% (v/v) phosphatase inhibitor cocktail 1, and 2.6% (v/v) phosphatase inhibitor cocktail 2) to each collected fraction immediately after the elution was complete, and stored them at −80°C.

### Antibody crosslinking to Dynabeads Protein G (DynaG) matrix

Covalent crosslinking of antibodies to matrix was performed as previously described [57]. Briefly, we incubated 62 µg of either the D8Q7I antibody (Cell Signaling Technology, Danvers, MA; Cat. No.: 12990S, RRID: AB_2798082) or the D3E10 antibody (Cell Signaling Technology, Cat. No.: 12843BF, RRID: AB_2798041) with 1 mL of DynaG matrix (Thermo Fisher Scientific, Cat. No.: 10004D) in 1 mL of extraction buffer at 4°C for 14-16 hr. To covalently crosslink the antibodies to the matrix, we incubated the immunocomplex in 1 mL of dimethyl pimelimidate (Thermo Fisher Scientific, Cat. No.: PI21667; dissolved in triethanolamine buffer solution (MilliporeSigma, Cat. No.: T0449); final concentration, 70 mM; final pH, 9) solution at room temperature for 15 min. We then quenched the reaction by incubating the antibody-matrix immunocomplex in 1 mL of 10% (v/v) ethanolamine (MilliporeSigma, Cat. No.: 398136; diluted the stock solution 10-fold in distilled deionized water) solution at room temperature for 15 min. Before use, we treated the resulting matrix once with elution buffer I (100 mM glycine (MilliporeSigma, Cat. No.: G8898)-HCl, 1 M urea (MilliporeSigma, Cat. No.: U5378), pH 2.9) by gently resuspending at room temperature for 30 sec. We then treated the resulting matrix twice with elution buffer II (Pierce IgG Elution Buffer (Thermo Fisher Scientific, Cat. No.: 21004), pH 2.9; 1% (v/v) *n*-octyl-1-thio-*β*-*D*-glycopyranoside (OTG); and 2 M urea) by agitating at 50°C, 1,200 rpm for 5 min each time. These elution steps were performed to elute off antibodies that were not crosslinked to the matrix. We then neutralized the pH of the antibody-immobilizing matrix using the extraction buffer and stored it at 4°C in the extraction buffer that contained 0.05% weight/volume (w/v) sodium azide (MilliporeSigma, Cat. No.: S8032) and inhibitors (0.1 mM PMSF, 0.2 mM phen, 0.1% (v/v) protease inhibitor cocktail, 0.1% (v/v) phosphatase inhibitor cocktail 1, and 0.1% (v/v) phosphatase inhibitor cocktail 2).

### Immunoprecipitation (IP)

To detect and measure levels of Aβ(40)*56 and monomeric Aβ(1-40) proteins, we diluted 1.4 mg of proteins from the IgG-depleted aqueous human brain extracts or 0.2 mg of proteins from the IgG-depleted aqueous mouse brain extracts into IP dilution buffer (50 mM Tris-HCl, pH 7.4; 150 mM NaCl) containing inhibitors (0.1 mM PMSF, 0.2 mM phen, 0.1% (v/v) protease inhibitor cocktail, 0.1% (v/v) phosphatase inhibitor cocktail 1, and 0.1% (v/v) phosphatase inhibitor cocktail 2) so that the final volume was 500 µL. We incubated each of the resulting samples with 50 µL of D8Q7I-bound DynaG matrix slurry (*i.e.*, 2-3 µL of settled beads) by rotating them at 4°C, 15 rpm for 14-16 hr. Following the formation of immuno-complexes, we washed the matrix twice in 500 µL of IP wash buffer (50 mM Tris-HCl, pH 7.4; 150 mM NaCl; 1% (v/v) OTG (Abcam, Cambridge, MA; Cat. No.: ab141435); and inhibitors (0.1 mM PMSF, 0.2 mM phen, 0.1% (v/v) protease inhibitor cocktail, 0.1% (v/v) phosphatase inhibitor cocktail 1, and 0.1% (v/v) phosphatase inhibitor cocktail 2)), each time at 4°C, 15 rpm for 5 min. We then eluted the immunocaptured proteins by agitating the matrix in 20 µL of elution buffer II at 50°C, 1,200 rpm for 5 min. One µL of a neutralization solution (1 M Tris, pH not adjusted (pH ∼10.5 upon measurement); 2.1 mM PMSF, 4.2 mM phen, 2.1% (v/v) protease inhibitor cocktail, 2.1% (v/v) phosphatase inhibitor cocktail 1, and 2.1% (v/v) phosphatase inhibitor cocktail 2) was immediately added to the eluted protein solution to neutralize the pH between 7 and 8.

To detect and measure levels of Aβ(42)*56 and monomeric Aβ(1-42) proteins, the same procedures described above were used except that aqueous brain extracts were incubated with D3E10-bound DynaG matrix.

### Western blotting (WB)

WB was performed based on previously published protocols [55, 57]. Detailed use of brain extracts and reagents are shown in Table S3.

Briefly, eluted materials from brain extracts (Fig. **1**b and d, Online Resource **1**: Figure S4, S5, S7, and S8), IP (Fig. **1**b and d, Online Resource **1**: Figure S10-S13), purified Aβ*56 (Fig. **1**f and g, Online Resource **1**: Figure S2), and SEC-fractionated brain extracts (Fig. **1**h, Online Resource **1**: Figure S3) were electrophoretically separated by SDS-polyacrylamide gel electrophoresis (PAGE). To prepare samples for SDS-PAGE, we mixed brain extracts (Fig. **1**b and d, Online Resource **1**: Figure S4, S5, S7 and S8), eluted materials (Fig. **1**b and d, Online Resource **1**: Figure S10-S13), or SEC-fractionated brain extracts (Fig. **1**h, Online Resource **1**: Figure S3) with 4x Laemmli sample buffer (Bio-Rad, Cat. No.: 161-0747) plus 1.42 M β-mercaptoethanol (BME; diluted stock BME (MilliporeSigma, Cat. No.: M6250) 10-fold in 4x Laemmli sample buffer) at 1,000 rpm, 95°C for 5 min. Purified Aβ*56 (Fig. **1**f and g, Online Resource **1**: Figure S2) was mixed with 4x Laemmli sample buffer without BME, agitation, or heating. We performed electrophoresis at room temperature in running buffer (100 mM Tris-HCl, pH 8.3; 100 mM *N*-[1,3-Dihydroxy-2-(hydroxymethyl)propan-2-yl]glycine (Tricine; MilliporeSigma, Cat. No.: T9784), and 0.1% (w/v) SDS (Thermo Fisher Scientific, Cat. No.: BP166)) under a constant voltage of 125 V for 90 min using 1.0-mm-thick, 10-(Thermo Fisher Scientific, Cat. No.: EC6625BOX) or 12-(Thermo Fisher Scientific, Cat. No.: EC66252BOX) well Novex 10-20% Tricine gels and then electrophoretically transferred the fractionated proteins onto 0.2-μm nitrocellulose membranes (Bio-Rad, Cat. No.: 162-0112) in transfer buffer (25 mM *N*,*N*-Bis(2-hydroxyethyl)glycine (Bicine; MilliporeSigma, Cat. No.: B3876), 25 mM 2-[Bis(2-hydroxyethyl)amino]-2-(hydroxymethyl)propane-1,3-diol (Bis-tris methane; MilliporeSigma, Cat. No.: RDD013), 1 mM ethylenediaminetetraacetic acid (EDTA; MilliporeSigma, Cat. No.: E6758-100G), pH 7.2; and 10% (v/v) methanol (Thermo Fisher Scientific, Cat. No.: A412)) under a constant voltage of 25 V at 4°C for 2 hr.

Next, we performed heat-induced antigen retrieval to enhance signal detection and then blocked the membranes at room temperature for 1 hr prior to probing with detection agents. For WB analyses using biotinylated A11 (StressMarq Biosciences, Victoria, BC; Cat. No.: SPC-506D-BI; RRID: AB_10962958) antibodies (Fig. **1**f and g) and their associated secondary detection agent horseradish peroxidase (HRP)-conjugated NeutrAvidin (NA) (Online Resource **1**: Figure S2), we used blocking buffer 1 (10 mM Tris-HCl, pH 7.4; 200 mM NaCl; 0.01% (v/v) polyoxyethylene (20) sorbitan monolaurate (Tween 20; MilliporeSigma, Cat. No.: P9416-100ML); and 5% (w/v) Carnation instant non-fat dry milk powder (Nestlé, Rosslyn, VA; Cat. No.: 12428935)); and for all other WB analyses, we used blocking buffer 2 (10 mM Tris-HCl, pH 7.4; 200 mM NaCl; 0.1% (v/v) Tween 20; and 5% (w/v) bovine serum albumin (MilliporeSigma, Cat. No.: A3803-100G)).

Next, we added primary antibodies (see Online Resource **1**: Table S3 for detailed use of antibodies) to the blocking buffers and incubated membranes at 4°C for 14-16 hr (For Online Resource **1**: Figure S2 and S3, membranes were incubated directly with blocking buffers at 4°C for 14-16 hr). Following primary antibody incubation, we washed membranes with wash buffers four times, each time at room temperature for 5 min. For WB analyses using biotinylated A11 antibodies (Fig. **1**f and g), we used WB wash buffer 1 (10 mM Tris-HCl, pH 7.4; 200 mM NaCl; and 0.01% (v/v) Tween 20); and for all other WB analyses, we used WB wash buffer 2 (10 mM Tris-HCl, pH 7.4; 200 mM NaCl; and 0.1% (v/v) Tween 20).

For WB analyses that were probed with non-biotinylated antibodies (Fig. **1**b and d, Online Resource **1**: Figure S4, S5, S7 and S8, Table S3), following the wash steps, we incubated membranes with HRP-conjugated ImmunoPure goat-anti-rabbit IgG (Thermo Fisher Scientific, Cat. No.: 31463; diluted 1:200,000 in WB wash buffer 2 for rabbit monoclonal anti-NeuN antibody clone 27-4 (MilliporeSigma, Cat. No.: MABN140; RRID: AB_2571567; Fig. **1**b and d, Online Resource **1**: Figure S7 and S8, Table S3) or rabbit monoclonal anti-GAPDH antibody clone 14C10 (Cell Signaling Technology, Cat. No.: 2118S; RRID: AB_561053; Fig. **1**b and d, Online Resource **1**: Figure S4 and S5, Table S3)) at room temperature for 1 hr. Following the incubation with the secondary antibody, we washed membranes using WB wash buffer 2 as described above.

For WB analyses that were probed with biotinylated primary antibodies (Fig. **1**b, d, f, g and h, Online Resource **1**: Figure S10-S13) or no primary antibody (Online Resource **1**: Figure S2 and S3), we directly incubated membranes with HRP-conjugated NA (Thermo Fisher Scientific, Cat. No.: A2664; diluted 1:5,000 in either WB wash buffer 1 (Fig. **1**f and g, Online Resource **1**: Figure S2) or WB wash buffer 2 (Fig. **1**b, d and h, Online Resource **1**: Figure S3 and S10-S13)) at room temperature for 30 min. Next, we washed membranes using WB wash buffers as described above.

For signal detection, we incubated membranes with either SuperSignal West Femto Maximum Sensitivity Substrate (Thermo Fisher Scientific, Cat. No.: 34095) at room temperature for 5 min (Fig. **1**b, d, f, g and h; Online Resource **1**: Figure S2, S3 and S10-S13) or SuperSignal West Pico PLUS Chemiluminescent Substrate (Thermo Fisher Scientific, Cat. No.: 34580) at room temperature for 4 min (Fig. **1**b and d; Online Resource **1**: Figure S4, S5, S7 and S8) and collected chemiluminescence digital images using the ChemiDoc MP Imaging System (Bio-Rad, RRID: SCR_019037).

We performed densitometry-based semi-quantitative analyses of protein band intensities using Image Lab version 6.1 (Bio-Rad, Cat. No.: 17006130). To analyze band intensities of samples in different WB images, we selected WB images in which the unsaturated band intensities of internal controls (the ∼56-kDa entities in APP/TTA mice for Aβ*56 variants, the synthetic Aβ for monomeric Aβ, the 27-4 signals in Tg2576 mice for NeuN, or the 14C10 signals in Tg2576 mice for GAPDH) most closely matching each other. We then calculated the mean intensities of the internal controls, created unique normalization factors accordingly to molecules of interest for each image, adjusted the originally measured band intensities using the normalization factors, and reported the adjusted data.

Experimenters performing (IP/)WB and data analysis were blind to the demographic, pathological, and clinical characteristics of human individuals.

### Purification of Aβ*56

Purification of Aβ*56 was carried out as previously described [57]. Briefly, we used the prepared D8Q7I-bound matrix detailed in *Antibody crosslinking to Dynabeads Protein G (DynaG) matrix* to isolate from AD brain extracts Aβ(40)*56 and other Aβ(x-40) entities; in addition, we used the prepared D3E10-bound matrix to isolate Aβ(42)*56 and other Aβ(x-42) entities. We incubated brain extracts containing 1 mg of total proteins with 167 µL of matrix slurry containing 10.3 µg of antibodies at 4°C for 14-16 hr. We then washed and eluted the matrix as described in *Immunoprecipitation (IP)*. Immediately following the elution step, we adjusted the pH of the eluted materials to pH between 7 and 8 by adding the neutralization solution with its volume being 1/20^th^ volume of the eluted materials. To separate Aβ*56 from other Aβ entities, we fractionated the eluted materials by SDS-PAGE under semi-denaturing conditions using 1.0-mm-thick Novex 10–20% Tricine gels (Thermo Fisher Scientific, Cat. No.: EC6625BOX or EC66252BOX). We used 10% of the fractionated proteins for WB probed with the biotinylated 82E1 antibody (IBL America, Cat. No.: 10326, RRID: AB_10705565; procedures detailed in *Western blotting (WB)*) to guide the identification and excision of Aβ*56-containing gel pieces for the other 90% of eluted materials. Next, we performed electro-elution to release Aβ*56 from gel using Bio-Rad Model 422 Electro-eluter (Bio-Rad, Cat. No.: 1652976) based on the manufacturer’s instructions. We eluted Aβ*56 using electro-elution buffer (50 mM ammonium bicarbonate (NH_4_HCO_3_; MilliporeSigma, Cat. No.: A6141-500G), pH not adjusted (pH ∼8.8 upon measurement); and 0.1% (w/v) SDS) at room temperature. We then removed SDS using High Protein and Peptide Recovery Detergent Removal Resin (Thermo Fisher Scientific, Cat. No.: 88305) following the manufacturer’s instructions and stored the final product at −80°C.

### Denaturant treatment

We treated with denaturants Aβ*56 variants that were purified from aqueous AD brain extracts. Following electro-elution, we completely dried Aβ*56 variants that were isolated from brain extracts containing 1.5 mg of total proteins and stored in 20 µL of 50 mM NH_4_HCO_3_, pH ∼8.8 at room temperature using a speed vacuum concentrator (Eppendorf, Hamburg, Germany; Model 5301). We then reconstituted using 20 µL of Tris buffer (20 mM Tris-HCl, pH 7.4) or 6 M guanidine hydrochloride (GuHCl) solution (20 mM Tris-HCl, pH 7.4; and 6 M GuHCl (PanReac AppliChem ITW Reagents, Darmstadt, Germany; Cat. No.: A1499)). Next, we vigorously agitated the solutions by shaking at room temperature, 1,400 rpm for 2 hr. The resulting materials were subjected to WB as described in *Western blotting (WB)*.

### Statistical analysis

Statistical analyses were performed using GraphPad Prism Version 10.2.0 (GraphPad Software, La Jolla, CA) and R Version 4.2.1 (R Foundation for Statistical Computing, Vienna, Austria). The demographic and neuropathological characteristics were compared between individuals with NCI, MCI, and AD dementia using Kruskal-Wallis tests or two-sided Fisher’s exact tests. The associations between demographic characteristics and levels of Aβ*56 variants (normalized to levels of NeuN proteins) were estimated using Spearman’s rank-order correlations for continuous variables or Mann-Whitney *U* tests for binary variables. Levels of Aβ*56 variants and monomeric Aβ (normalized to levels of NeuN proteins), NeuN, and GAPDH were compared between individuals with NCI, MCI, and AD dementia using Kruskal-Wallis tests followed by post hoc analysis using Dunn’s correction. The correlations between levels of Aβ*56 variants and monomeric Aβ (normalized to levels of NeuN proteins) and clinical and pathological characteristics were estimated using Spearman’s rank-order correlations. Additionally, multiple linear regressions (MLRs) were fit to compare the log-transformed levels between the groups (NCI, MCI, and AD dementia) and assess the association between log-transformed levels and cognitive outcomes with adjustment for the potential confounders, specified below. When comparing levels between the groups, *post hoc* pairwise comparisons were made when the overall *F*-test was statistically significant with simultaneous confidence intervals (CIs) calculated using the *multcomp* R package [32]. Mediation analyses based on the analogous MLRs were performed using the *mediation* R package [89]. *P* values of less than 0.05 were considered statistically significant.

## Results

### Aβ(40)*56 and Aβ(42)*56 are detected in the ITG of elderly human individuals

We chose the ITG because this is the first neocortical region in which NFT appear in Braak Stages III-IV [10], coinciding with isolated memory impairment and sparing of other cognitive domains [88], the earliest sign of AD dementia. In the current cohort, ITG NFT loads were determined using the phosphorylated tau antibody AT8 which stains ptau 202/205 [26] and were significantly higher in AD dementia than MCI and NCI. However, those in MCI and NCI did not differ (Online Resource **1**: Figure S15), suggesting that AD neuropathology had not yet spread to MCI. This similarity between MCI and NCI neuropathology in the ITG is consistent with a previously published finding using the ROSMAP cohort [53].

First, to measure Aβ*56, we used the 82E1 antibody that recognizes the N-terminus of Aβ(1-x) to detect ∼56-kDa entities and monomeric Aβ peptides from aqueous brain extracts of human individuals precipitated by antibodies that target the C-terminus of Aβ(x-40) (Fig. **1**a and b) or Aβ(x-42) (Fig. **1**c and d). These results indicate the presence of ∼56-kDa, SDS-stable, water-soluble protein aggregates that contain canonical Aβ(1-40) and Aβ(1-42).

Next, to verify that the ∼56-kDa entities were oligomers, we isolated them using immunoaffinity purification coupled with electro-elution (Fig. **1**e), and tested their reactivity to A11 antibodies that bind non-fibrillar oligomers [41, 48, 54] as well as their stability upon chaotropic denaturant treatment. The isolated ∼56-kDa entities bound A11 antibodies (Fig. **1**f and g) but not the secondary detecting agent HRP-conjugated NA (Online Resource **1**: Figure S2). However, treating with 6 M GuHCl abolished A11 binding (Fig. **1**f and g). These results indicate that the ∼56-kDa entities are Aβo that adopt structures sensitive to denaturant treatment. We named them Aβ(40)*56 and Aβ(42)*56 to represent oligomers containing Aβ(1-40) and Aβ(1-42), respectively.

To confirm that Aβ*56 is not artificially generated from Aβ monomers exposed to SDS, we used non-denaturing SEC to fractionate aqueous brain extracts from individuals with AD dementia. We revealed WB patterns of eluted fractions probed with the biotinylated 82E1 antibody (Fig. **1**h) and identified non-specific, endogenous biotinylated proteins using the secondary detection agent HRP-conjugated NA only (Online Resource **1**: Figure S3). We found two clusters of ∼56-kDa, 82E1-reactive entities, one in fractions 63-71 corresponding to globular protein sizes of ∼35-84-kDa with a peak at ∼57 kDa, the other in fractions 51-61 corresponding to globular protein sizes of ∼105-318-kDa with a peak at ∼164 kDa (Fig. **1**h and i). The ∼57-kDa peak likely corresponds to singular Aβ*56 oligomers, while the ∼164-kDa peak likely corresponds to small assemblies of Aβ*56 and/or Aβ*56 complexed with other biomolecules. Fractions containing Aβ*56 contained minimal Aβ monomers, which were instead present in fractions 35-45 corresponding to globular protein sizes of ∼619 to >2,000 kDa with a peak at >2,000 kDa and fractions 75-83 that correspond to globular protein sizes of ∼9-22 kDa with a peak at ∼14 kDa (Fig. **1**h and i). The earlier peak may reflect large complexes of monomers and other proteins, while the later peak may reflect Aβo configured to form larger structures (*e.g.*, longer hydration radii) than globular proteins of the same molecule mass. In addition, we found ∼14-kDa entities appearing in fractions 65-79 corresponding to globular protein sizes of ∼14-68 kDa with a peak at ∼43 kDa, and ∼9-kDa entities in fractions 67-77 corresponding to globular protein sizes of ∼18-54 kDa with a peak at ∼28 kDa (Fig. **1**h). The ∼14-kDa and ∼9-kDa entities may represent trimeric and dimeric Aβ species, respectively, that are larger in volume than globular proteins with similar molecular masses. These SEC results are consistent with previous findings [74].

Altogether, these results indicate that human brains contain Aβ(40)*56 and Aβ(42)*56 composed of canonical Aβ(1-40) and Aβ(1-42), respectively. Aβ(40)*56 and Aβ(42)*56 are non-fibrillar, SDS-stable, water-soluble oligomers with molecular masses of ∼56-kDa under both denaturing SDS-PAGE and non-denaturing SEC. They contain denaturant-sensitive A11-reactive epitopes and are not artificially formed from monomeric or low-molecular-weight Aβ species exposed to SDS.

### Levels oxf Aβ*56 are elevated in AD dementia independently of amyloid pathology

The individuals used to measure Aβ(40)*56 and Aβ(42)*56 partially overlapped (*N* = 11 NCI, *N* = 8 MCI, and *N* = 15 AD dementia were common to both cohorts). This is because certain brain specimens were too small in size to measure both Aβ*56 variants; protein extracts prepared from these specimens were used to measure only one Aβ*56 variant.

To control for protein loading, we measured the housekeeping protein GAPDH and found no difference across groups (for Aβ(40)*56, *H* (2) = 0.82, *P* = 0.67, Kruskal-Wallis test, Online Resource **1**: Figure S4 and S6a; for Aβ(42)*56, *H* (2) = 0.22, *P* = 0.90, Kruskal-Wallis test, Online Resource **1**: Figure S5 and S6b), which indicates comparable levels of total proteins between individual samples are used for the experiments. Because post-mitotic neurons are the principal cell type that generate Aβ [70, 93] – the substrate of Aβo, we measured NeuN, a protein marker for post-mitotic neurons, and used its levels as a normalization factor (Online Resource **1**: Figure S7 and S8) for Aβ*56 and monomeric Aβ levels. We showed that NeuN levels did not differ significantly across groups (for Aβ(40)*56, *H* (2) = 2.71, *P* = 0.26; for Aβ(42)*56, *H* (2) = 4.99, *P* = 0.08; Kruskal-Wallis tests; Online Resource **1**: Figure S9a and S9b).

To assess the association of Aβ*56 variants with AD progression independently of plaques, we measured their levels in the ITG of individuals with AD dementia, MCI, and NCI (see Online Resource **1**: Table S1 and S2 for detailed demographic and neuropathological characteristics), using IP/WB to detect Aβ(40)*56 (Online Resource **1**: Figure S10) and Aβ(42)*56 (Online Resource **1**: Figure S11). The three groups (NCI, MCI, and AD dementia) were comparable in age at death, sex distribution, post-mortem interval (PMI), years of education, and apolipoprotein E (*APOE*) genotype distribution (Tables **1** and **2**). Importantly, the three groups did not differ significantly in ITG Aβ plaque load (Fig. **2**a and d, Tables **3** and **4**). Levels of Aβ(40)*56 in AD dementia were 4.0- and 4.7-fold higher than in NCI and MCI, respectively (*H* (2) = 12.83, *P* = 0.002; Kruskal-Wallis test; Fig. **2**b). Levels of Aβ(42)*56 in AD dementia were 2.4- and 3.1-fold higher than in NCI and MCI, respectively (*H* (2) = 8.28, *P* = 0.02; Kruskal-Wallis test; Fig. **2**e). In contrast, we found trends but no significant differences across groups in monomeric Aβ(1-40) (*H* (2) = 5.68, *P* = 0.06; Kruskal-Wallis test; Online Resource **1**: Figure S12 and S14a) or monomeric Aβ(1-42) (*H* (2) = 5.17, *P* = 0.08; Kruskal-Wallis test; Online Resource **1**: Figure S13 and S14b).

**Fig. 2.**
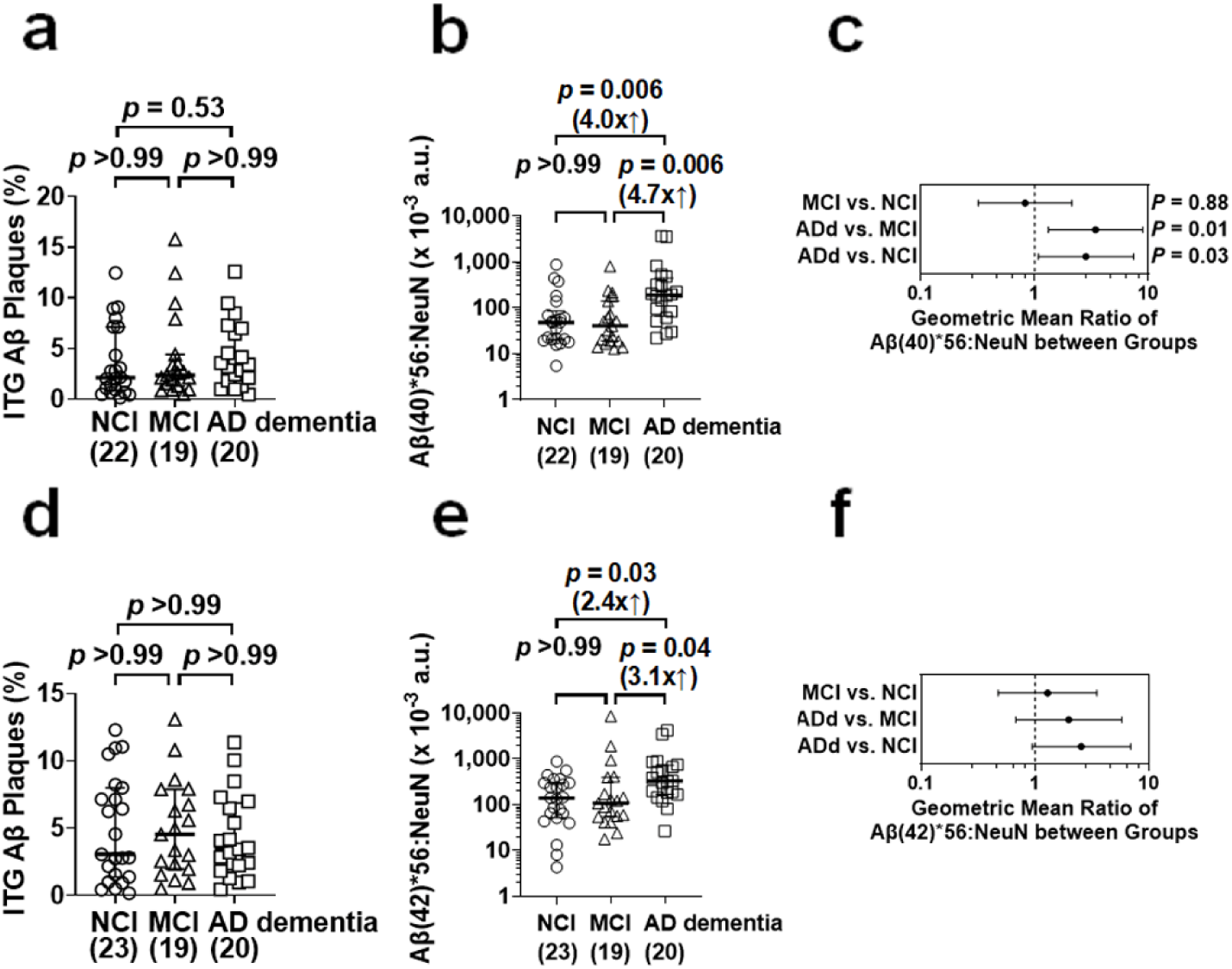
Levels of Aβ*56 variants are elevated in the inferior temporal gyri of individuals with AD dementia. (**a, d**) Following Kruskal-Wallis tests (Tables 3 and 4), *post hoc* analyses show no difference in Aβ plaque loads in the inferior temporal gyrus (ITG) between each pair of the individuals with no cognitive impairment (NCI), those with mild cognitive impairment (MCI), and those with Alzheimer’s disease dementia (AD dementia) that were used to measure Aβ(40)*56 (a) and Aβ(42)*56 (d). (**b, e**) Following a Kruskal-Wallis test (*H* (2) = 12.83, *P* = 0.002), *post hoc* analyses show that levels of Aβ(40)*56 in individuals with AD dementia, normalized to levels of NeuN, are 4.0- and 4.7- fold higher than normalized levels of Aβ(40)*56 in individuals with NCI and MCI, respectively (b). In parallel, following a Kruskal-Wallis test (*H* (2) = 8.28, *P* = 0.02), *post hoc* analyses show that levels of Aβ(42)*56 in individuals with AD dementia, normalized to levels of NeuN, are 2.4- and 3.1-fold higher than normalized levels of Aβ(42)*56 in individuals with NCI and MCI, respectively (e). In both cases, no difference in levels of Aβ*56 is present between individuals with NCI and MCI (b, e). (**c, f**) Following adjustments for age at death, sex, postmortem interval, and log-transformed ITG Aβ plaque loads using multiple linear regression, normalized levels of Aβ(40)*56 remain different between individuals with NCI, MCI, and AD dementia (ADd) (*P* = 0.01) as normalized levels of Aβ(40)*56 in individuals with ADd are higher than those in individuals with NCI as well as MCI (c). In parallel, following the adjustments, there is a trend towards difference in normalized levels of Aβ(42)*56 between the three groups (*P* = 0.08) (f). The numbers of individuals with NCI, MCI, and AD dementia used are shown in parentheses. a.u., arbitrary units.

**Table 1.**
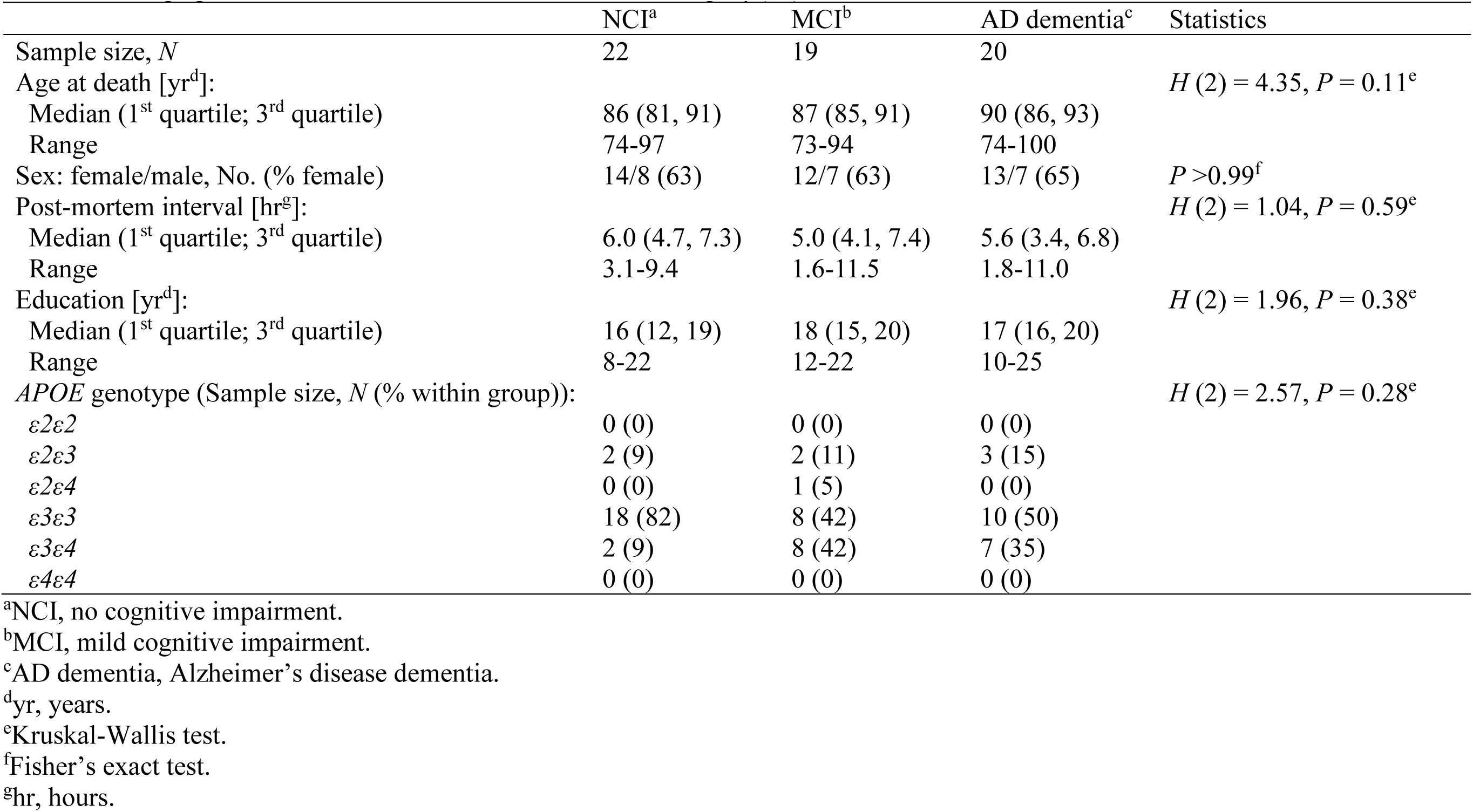
Demographic characteristics of individuals for measuring Aβ(40)*56.

|  | NCI<sup>a</sup> | MCI<sup>b</sup> | AD dementia<sup>c</sup> | Statistics |
| --- | --- | --- | --- | --- |
| Sample size, *N* | 22 | 19 | 20 |  |
| Age at death [yr<sup>d</sup>]: | | | | $H(2) = 4.35, P = 0.11^e$ |
| Median (1<sup>st</sup> quartile; 3<sup>rd</sup> quartile) | 86 (81, 91) | 87 (85, 91) | 90 (86, 93) |  |
| Range | 74-97 | 73-94 | 74-100 |  |
| Sex: female/male, No. (% female) | 14/8 (63) | 12/7 (63) | 13/7 (65) | $P > 0.99^f$ |
| Post-mortem interval [hr<sup>g</sup>]: | | | | $H(2) = 1.04, P = 0.59^e$ |
| Median (1<sup>st</sup> quartile; 3<sup>rd</sup> quartile) | 6.0 (4.7, 7.3) | 5.0 (4.1, 7.4) | 5.6 (3.4, 6.8) |  |
| Range | 3.1-9.4 | 1.6-11.5 | 1.8-11.0 |  |
| Education [yr<sup>d</sup>]: | | | | $H(2) = 1.96, P = 0.38^e$ |
| Median (1<sup>st</sup> quartile; 3<sup>rd</sup> quartile) | 16 (12, 19) | 18 (15, 20) | 17 (16, 20) |  |
| Range | 8-22 | 12-22 | 10-25 |  |
| *APOE* genotype (Sample size, *N* (% within group)): | | | | $H(2) = 2.57, P = 0.28^e$ |
| $\epsilon 2\epsilon 2$ | 0 (0) | 0 (0) | 0 (0) | |
| $\epsilon 2\epsilon 3$ | 2 (9) | 2 (11) | 3 (15) | |
| $\epsilon 2\epsilon 4$ | 0 (0) | 1 (5) | 0 (0) | |
| $\epsilon 3\epsilon 3$ | 18 (82) | 8 (42) | 10 (50) | |
| $\epsilon 3\epsilon 4$ | 2 (9) | 8 (42) | 7 (35) | |
| $\epsilon 4\epsilon 4$ | 0 (0) | 0 (0) | 0 (0) | |

<sup>a</sup>NCI, no cognitive impairment.<sup>b</sup>MCI, mild cognitive impairment.<sup>c</sup>AD dementia, Alzheimer's disease dementia.<sup>d</sup>yr, years.<sup>e</sup>Kruskal-Wallis test.<sup>f</sup>Fisher's exact test.<sup>g</sup>hr, hours.

**Table 2.**
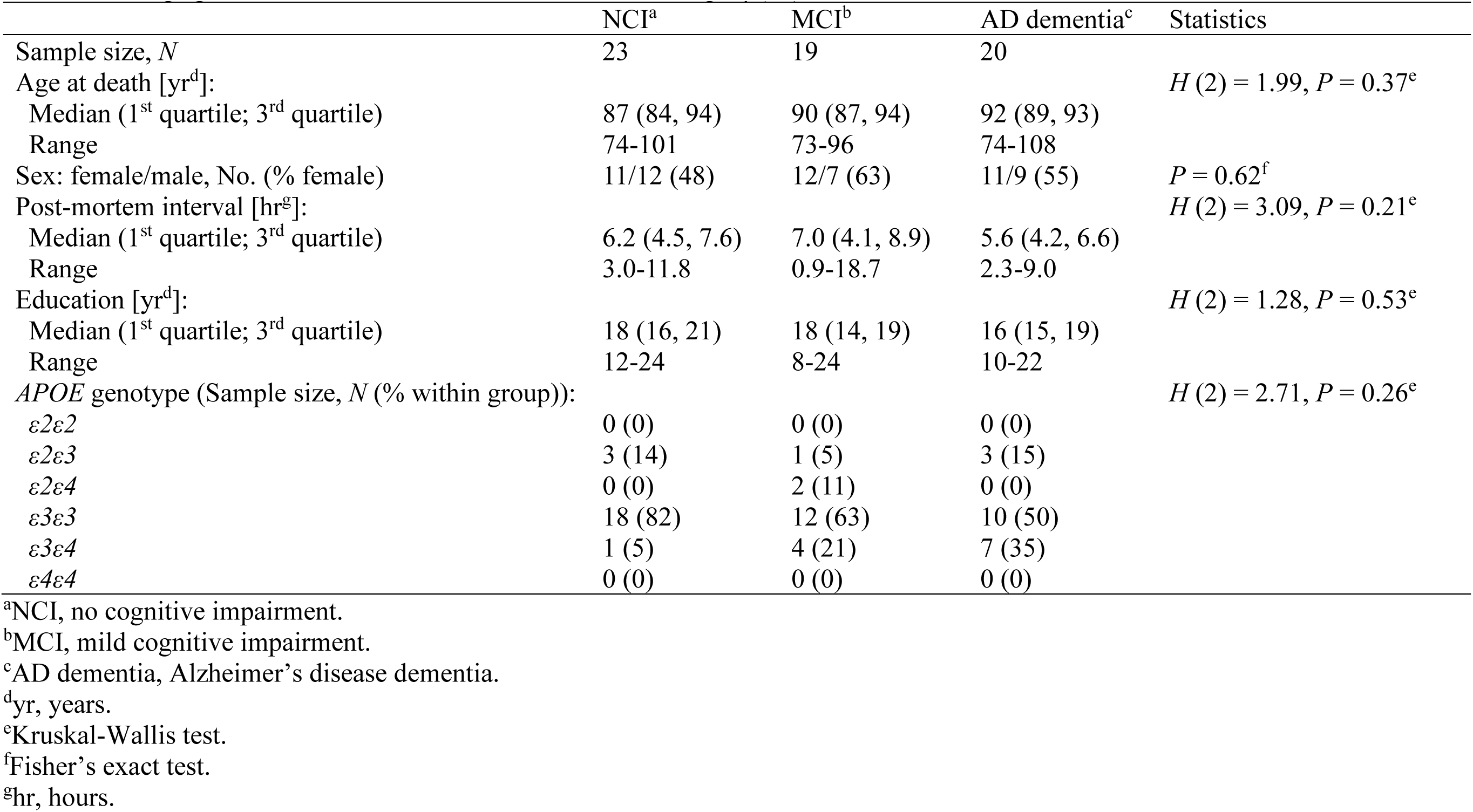
Demographic characteristics of individuals for measuring Aβ(42)*56.

**Table 3.**
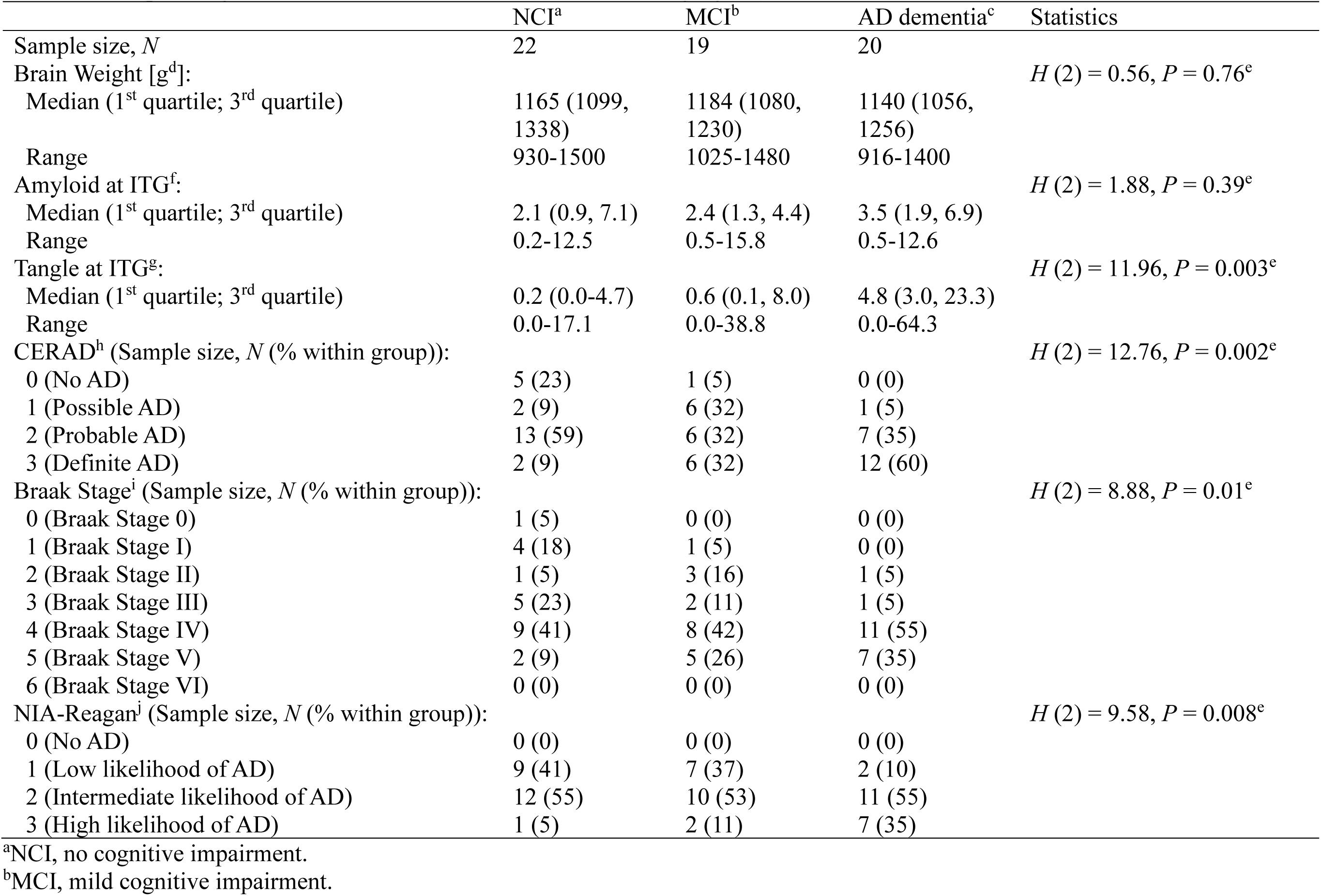
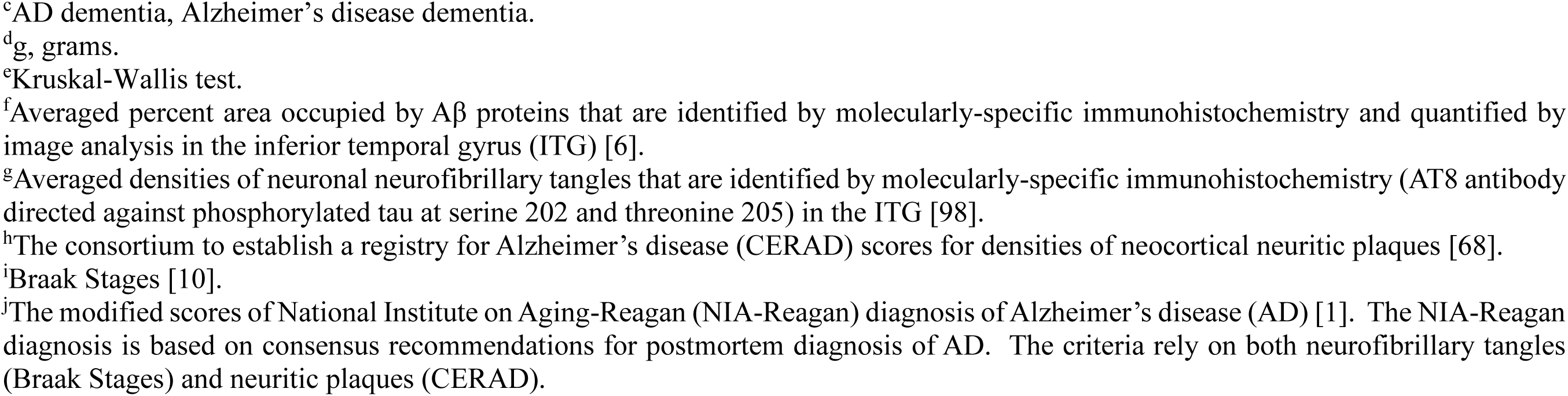
Neuropathological characteristics of individuals for measuring Aβ(40)*56.

|  | NCI<sup>a</sup> | MCI<sup>b</sup> | AD dementia<sup>c</sup> | Statistics |
| --- | --- | --- | --- | --- |
| Sample size, *N* | 22 | 19 | 20 |  |
| Brain Weight [g<sup>d</sup>]: | | | | $H(2) = 0.56, P = 0.76^e$ |
| Median (1<sup>st</sup> quartile; 3<sup>rd</sup> quartile) | 1165 (1099, 1338) | 1184 (1080, 1230) | 1140 (1056, 1256) |  |
| Range | 930-1500 | 1025-1480 | 916-1400 |  |
| Amyloid at ITG<sup>f</sup>: | | | | $H(2) = 1.88, P = 0.39^e$ |
| Median (1<sup>st</sup> quartile; 3<sup>rd</sup> quartile) | 2.1 (0.9, 7.1) | 2.4 (1.3, 4.4) | 3.5 (1.9, 6.9) |  |
| Range | 0.2-12.5 | 0.5-15.8 | 0.5-12.6 |  |
| Tangle at ITG<sup>g</sup>: | | | | $H(2) = 11.96, P = 0.003^e$ |
| Median (1<sup>st</sup> quartile; 3<sup>rd</sup> quartile) | 0.2 (0.0-4.7) | 0.6 (0.1, 8.0) | 4.8 (3.0, 23.3) |  |
| Range | 0.0-17.1 | 0.0-38.8 | 0.0-64.3 |  |
| CERAD<sup>h</sup> (Sample size, *N* (% within group)): | | | | $H(2) = 12.76, P = 0.002^e$ |
| 0 (No AD) | 5 (23) | 1 (5) | 0 (0) |  |
| 1 (Possible AD) | 2 (9) | 6 (32) | 1 (5) |  |
| 2 (Probable AD) | 13 (59) | 6 (32) | 7 (35) |  |
| 3 (Definite AD) | 2 (9) | 6 (32) | 12 (60) |  |
| Braak Stage<sup>i</sup> (Sample size, *N* (% within group)): | | | | $H(2) = 8.88, P = 0.01^e$ |
| 0 (Braak Stage 0) | 1 (5) | 0 (0) | 0 (0) |  |
| 1 (Braak Stage I) | 4 (18) | 1 (5) | 0 (0) |  |
| 2 (Braak Stage II) | 1 (5) | 3 (16) | 1 (5) |  |
| 3 (Braak Stage III) | 5 (23) | 2 (11) | 1 (5) |  |
| 4 (Braak Stage IV) | 9 (41) | 8 (42) | 11 (55) |  |
| 5 (Braak Stage V) | 2 (9) | 5 (26) | 7 (35) |  |
| 6 (Braak Stage VI) | 0 (0) | 0 (0) | 0 (0) |  |
| NIA-Reagan<sup>j</sup> (Sample size, *N* (% within group)): | | | | $H(2) = 9.58, P = 0.008^e$ |
| 0 (No AD) | 0 (0) | 0 (0) | 0 (0) |  |
| 1 (Low likelihood of AD) | 9 (41) | 7 (37) | 2 (10) |  |
| 2 (Intermediate likelihood of AD) | 12 (55) | 10 (53) | 11 (55) |  |
| 3 (High likelihood of AD) | 1 (5) | 2 (11) | 7 (35) |  |

<sup>a</sup>NCI, no cognitive impairment.<sup>b</sup>MCI, mild cognitive impairment.
<sup>c</sup>AD dementia, Alzheimer's disease dementia.
<sup>d</sup>g, grams.
<sup>e</sup>Kruskal-Wallis test.
<sup>f</sup>Averaged percent area occupied by Aβ proteins that are identified by molecularly-specific immunohistochemistry and quantified by image analysis in the inferior temporal gyrus (ITG) [6].
<sup>g</sup>Averaged densities of neuronal neurofibrillary tangles that are identified by molecularly-specific immunohistochemistry (AT8 antibody directed against phosphorylated tau at serine 202 and threonine 205) in the ITG [98].
<sup>h</sup>The consortium to establish a registry for Alzheimer's disease (CERAD) scores for densities of neocortical neuritic plaques [68].
<sup>i</sup>Braak Stages [10].
<sup>j</sup>The modified scores of National Institute on Aging-Reagan (NIA-Reagan) diagnosis of Alzheimer's disease (AD) [1]. The NIA-Reagan diagnosis is based on consensus recommendations for postmortem diagnosis of AD. The criteria rely on both neurofibrillary tangles (Braak Stages) and neuritic plaques (CERAD).

**Table 4.**
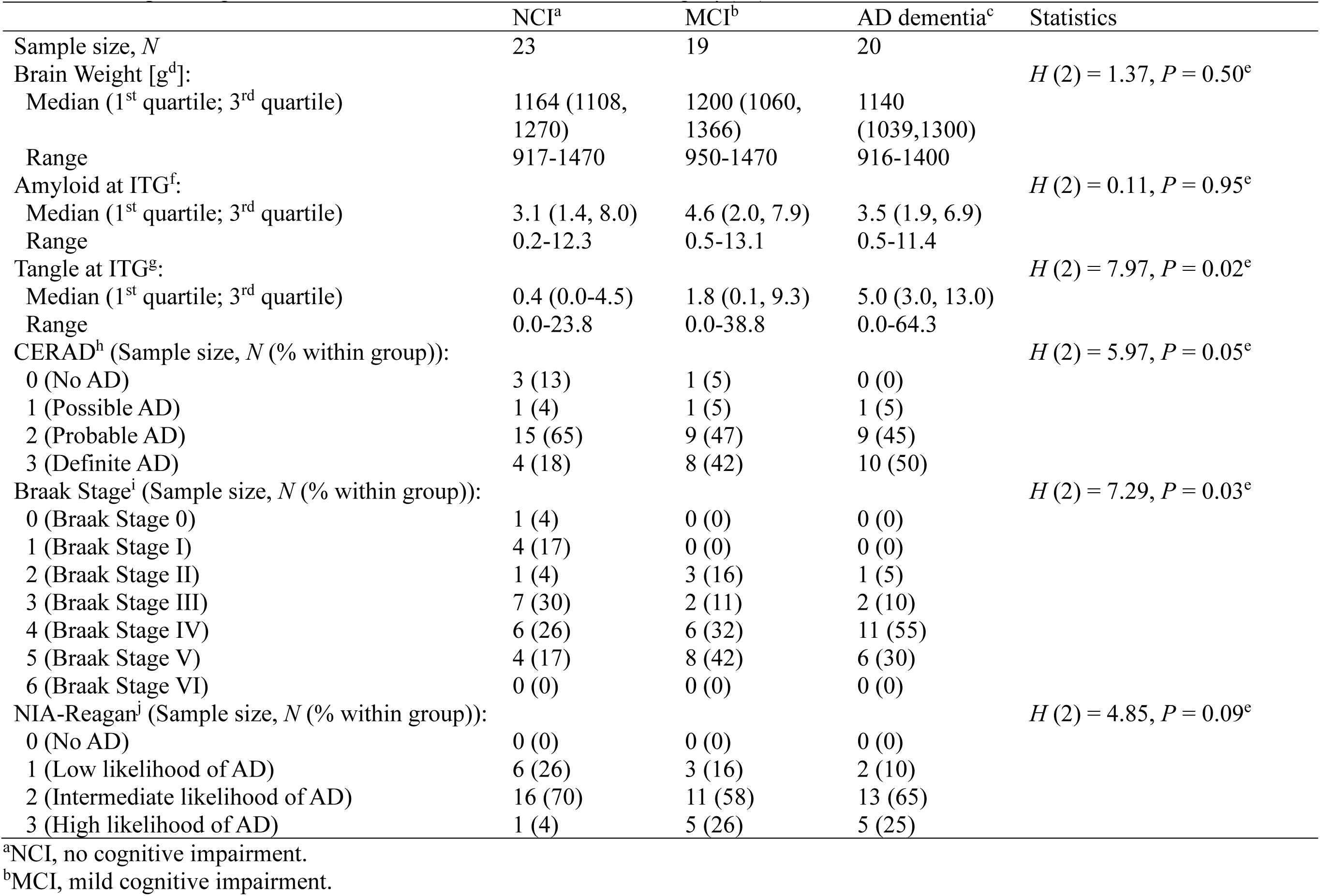
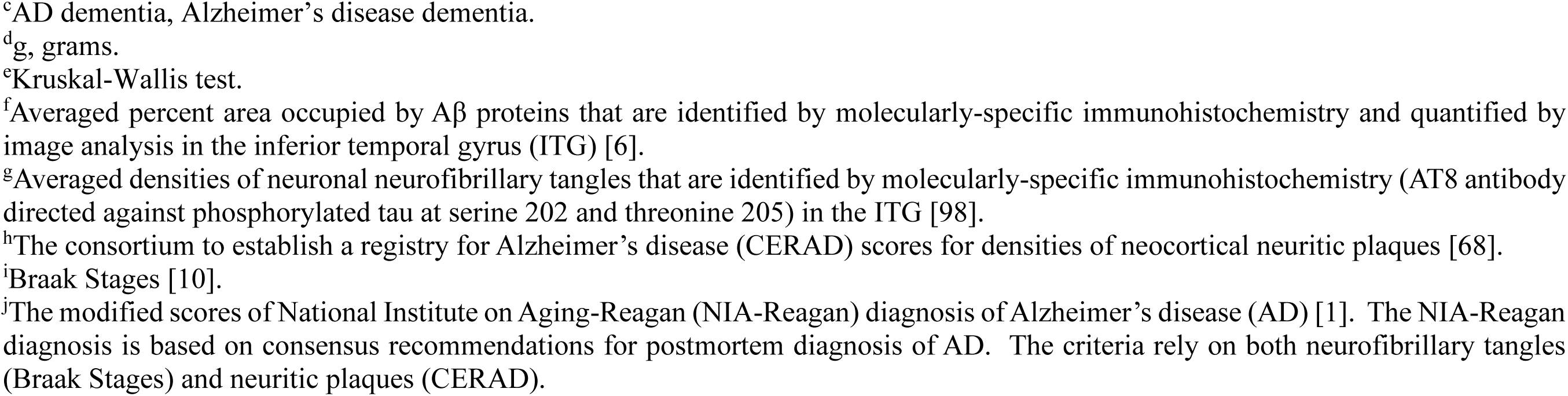
Neuropathological characteristics of individuals for measuring Aβ(42)*56.

|  | NCI<sup>a</sup> | MCI<sup>b</sup> | AD dementia<sup>c</sup> | Statistics |
| --- | --- | --- | --- | --- |
| Sample size, *N* | 23 | 19 | 20 |  |
| Brain Weight [g<sup>d</sup>]: | | | | $H(2) = 1.37, P = 0.50^e$ |
| Median (1<sup>st</sup> quartile; 3<sup>rd</sup> quartile) | 1164 (1108, 1270) | 1200 (1060, 1366) | 1140 (1039, 1300) |  |
| Range | 917-1470 | 950-1470 | 916-1400 |  |
| Amyloid at ITG<sup>f</sup>: | | | | $H(2) = 0.11, P = 0.95^e$ |
| Median (1<sup>st</sup> quartile; 3<sup>rd</sup> quartile) | 3.1 (1.4, 8.0) | 4.6 (2.0, 7.9) | 3.5 (1.9, 6.9) |  |
| Range | 0.2-12.3 | 0.5-13.1 | 0.5-11.4 |  |
| Tangle at ITG<sup>g</sup>: | | | | $H(2) = 7.97, P = 0.02^e$ |
| Median (1<sup>st</sup> quartile; 3<sup>rd</sup> quartile) | 0.4 (0.0-4.5) | 1.8 (0.1, 9.3) | 5.0 (3.0, 13.0) |  |
| Range | 0.0-23.8 | 0.0-38.8 | 0.0-64.3 |  |
| CERAD<sup>h</sup> (Sample size, *N* (% within group)): | | | | $H(2) = 5.97, P = 0.05^e$ |
| 0 (No AD) | 3 (13) | 1 (5) | 0 (0) |  |
| 1 (Possible AD) | 1 (4) | 1 (5) | 1 (5) |  |
| 2 (Probable AD) | 15 (65) | 9 (47) | 9 (45) |  |
| 3 (Definite AD) | 4 (18) | 8 (42) | 10 (50) |  |
| Braak Stage<sup>i</sup> (Sample size, *N* (% within group)): | | | | $H(2) = 7.29, P = 0.03^e$ |
| 0 (Braak Stage 0) | 1 (4) | 0 (0) | 0 (0) |  |
| 1 (Braak Stage I) | 4 (17) | 0 (0) | 0 (0) |  |
| 2 (Braak Stage II) | 1 (4) | 3 (16) | 1 (5) |  |
| 3 (Braak Stage III) | 7 (30) | 2 (11) | 2 (10) |  |
| 4 (Braak Stage IV) | 6 (26) | 6 (32) | 11 (55) |  |
| 5 (Braak Stage V) | 4 (17) | 8 (42) | 6 (30) |  |
| 6 (Braak Stage VI) | 0 (0) | 0 (0) | 0 (0) |  |
| NIA-Reagan<sup>j</sup> (Sample size, *N* (% within group)): | | | | $H(2) = 4.85, P = 0.09^e$ |
| 0 (No AD) | 0 (0) | 0 (0) | 0 (0) |  |
| 1 (Low likelihood of AD) | 6 (26) | 3 (16) | 2 (10) |  |
| 2 (Intermediate likelihood of AD) | 16 (70) | 11 (58) | 13 (65) |  |
| 3 (High likelihood of AD) | 1 (4) | 5 (26) | 5 (25) |  |

<sup>a</sup>NCI, no cognitive impairment.<sup>b</sup>MCI, mild cognitive impairment.
<sup>c</sup>AD dementia, Alzheimer's disease dementia.
<sup>d</sup>g, grams.
<sup>e</sup>Kruskal-Wallis test.
<sup>f</sup>Averaged percent area occupied by Aβ proteins that are identified by molecularly-specific immunohistochemistry and quantified by image analysis in the inferior temporal gyrus (ITG) [6].
<sup>g</sup>Averaged densities of neuronal neurofibrillary tangles that are identified by molecularly-specific immunohistochemistry (AT8 antibody directed against phosphorylated tau at serine 202 and threonine 205) in the ITG [98].
<sup>h</sup>The consortium to establish a registry for Alzheimer's disease (CERAD) scores for densities of neocortical neuritic plaques [68].
<sup>i</sup>Braak Stages [10].
<sup>j</sup>The modified scores of National Institute on Aging-Reagan (NIA-Reagan) diagnosis of Alzheimer's disease (AD) [1]. The NIA-Reagan diagnosis is based on consensus recommendations for postmortem diagnosis of AD. The criteria rely on both neurofibrillary tangles (Braak Stages) and neuritic plaques (CERAD).

Next, we investigated whether Aβ*56 correlated with demographic characteristics. We found no significant effects of sex (Online Resource **1**: Figure S16), PMI (Online Resource **1**: Figure S17), or years of education (Online Resource **1**: Figure S18). Of note, we observed a significant age-associated elevation of Aβ(40)*56 (Spearman *ρ* = 0.35, *P* = 0.006; Online Resource **1**: Figure S19a) and a trend for Aβ(42)*56 (Spearman *ρ* = 0.23, *P* = 0.07; Online Resource **1**: Figure S19b).

To control for the potential impact of demographic characteristics and plaque load on Aβ*56 levels, we adjusted for age at death, sex, PMI, and log-transformed ITG Aβ plaque load using MLR. Adjusted levels of Aβ(40)*56 remained significantly different (*P* = 0.01, MLR *F* test); adjusted levels of Aβ(40)*56 were higher in AD dementia than NCI (geometric mean ratio (95% CI): 2.83 (1.08, 7.41), *P* = 0.03; Fig. **2**c) and MCI (geometric mean ratio (95% CI): 3.44 (1.32, 8.98), *P* = 0.01; Fig. **2**c). Adjusted levels of Aβ(42)*56 trended towards difference between the three groups (*P* = 0.08, MLR *F* test; Fig. **2**f). These data confirm that Aβ(40)*56 is elevated and Aβ(42)*56 trends higher in AD dementia.

### Levels of Aβ*56 are associated with global AD pathology

Aβo have been proposed to induce tau hyperphosphorylation and promote the spread of tau pathology [3, 31, 36, 37, 104]. To determine whether Aβ*56 may contribute to global AD pathology, we examined the association between Aβ*56 and global neuropathology. We identified significant correlations between Aβ(40)*56 levels and modified scores of National Institute on Aging-Reagan (NIA-Reagan) diagnosis of AD [1] (Spearman *ρ* = 0.41, *P* = 0.001; Fig. **3**a), Consortium to Establish a Registry for Alzheimer’s Disease (CERAD) [68] scores (Spearman *ρ* = 0.39, *P* = 0.002; Fig. **3**b), Braak Stage [10] (Spearman *ρ* = 0.34, *P* = 0.007; Fig. **3**c), immunohistologically-determined ITG Aβ plaque loads (Spearman *ρ* = 0.36, *P* = 0.004; Fig. **3**d), and ITG ptau 202/205-containing NFT loads (Spearman *ρ* = 0.38, *P* = 0.002; Fig. **3**e).

**Fig. 3.**
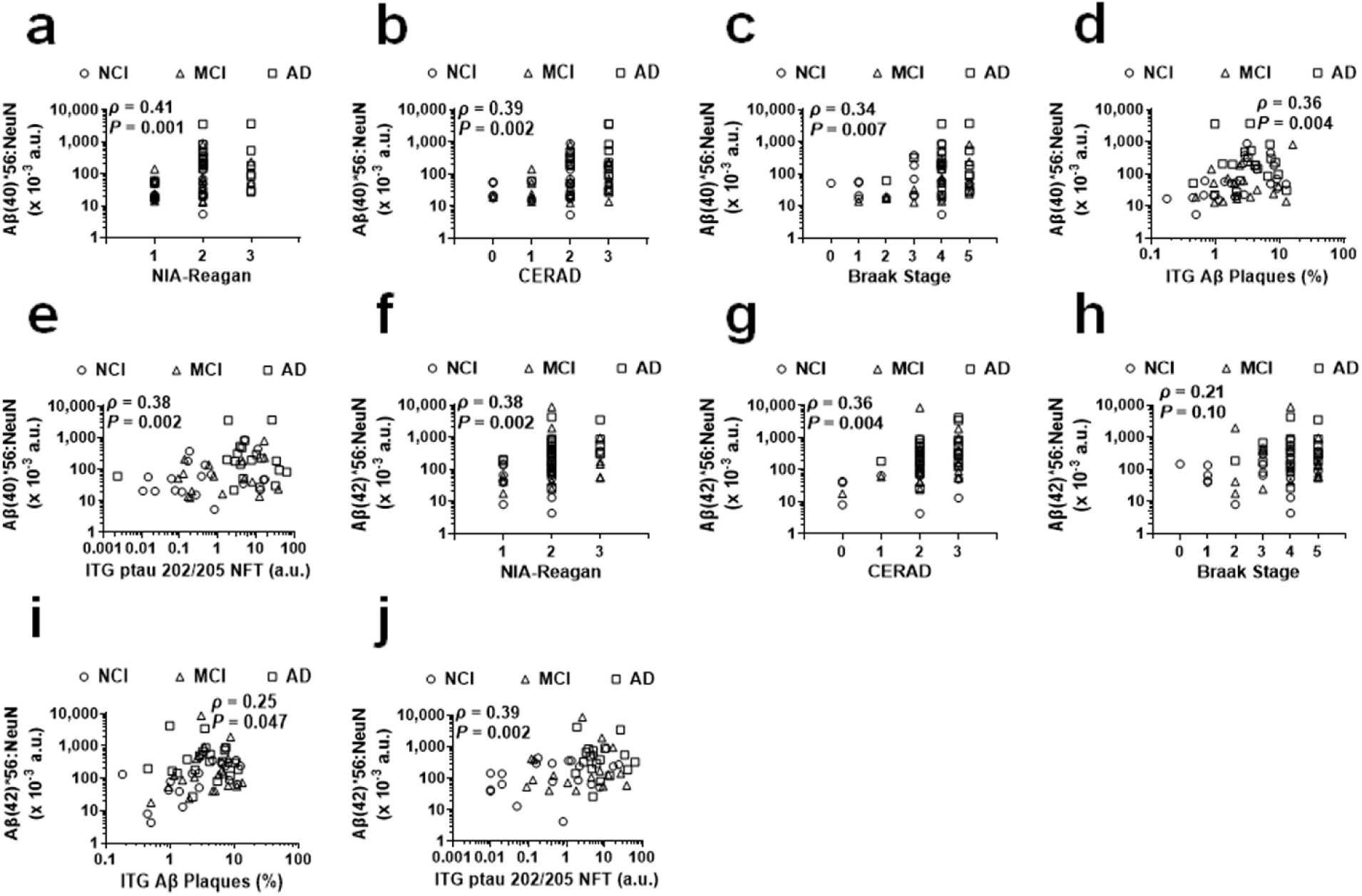
Aβ*56 variants are associated with AD pathology. (**a, f**) Correlations of NeuN-normalized levels of Aβ(40)*56 (a) and Aβ(42)*56 (f) to modified NIA-Reagan diagnosis of AD scores (1, low likelihood of AD; 2, intermediate likelihood of AD; and 3, high likelihood of AD). (**b, g**) Correlations of NeuN-normalized levels of Aβ(40)*56 (b) and Aβ(42)*56 (g) to Consortium to Establish a Registry for Alzheimer’s Disease (CERAD) scores (0, no AD; 1, possible AD; 2, probable AD; and 3, definite AD). (**c, h**) Correlations of NeuN-normalized levels of Aβ(40)*56 (c) and Aβ(42)*56 (h) to Braak Stage values (0, Braak Stage 0; 1, Braak Stage I; 2, Braak Stage II; 3, Braak Stage III; 4, Braak Stage IV; and 5, Braak Stage V). (**d, i**) Correlations of NeuN-normalized levels of Aβ(40)*56 (d) and Aβ(42)*56 (i) to percent areas occupied by Aβ plaques in the inferior temporal gyrus (ITG Aβ Plaques). (**e, j**) Correlations of NeuN-normalized levels of Aβ(40)*56 (e) and Aβ(42)*56 (j) to densities (densities of AT8-positive NFT per mm^2^ of systematically sampled brain regions) of neuronal neurofibrillary tangles that are reactive to antibody AT8 directed against phosphorylated tau at serine 202 and threonine 205 in the ITG (ITG ptau 202/205 NFT). Spearman’s rank-order correlations were used. The x-axes of all figures and the y-axes of Figures d, e, i, and j are displayed on log scale. NCI, no cognitive impairment; MCI, mild cognitive impairment; AD dementia, Alzheimer’s disease dementia. a.u., arbitrary units.

In addition, we found significant correlations between Aβ(42)*56 levels and modified scores of NIA-Reagan diagnosis of AD (Spearman *ρ* = 0.38, *P* = 0.002; Fig. **3**f), CERAD scores (Spearman *ρ* = 0.36, *P* = 0.004; Fig. **3**g), ITG Aβ plaque loads (Spearman *ρ* = 0.25, *P* = 0.047; Fig. **3**i), and ITG ptau 202/205-containing NFT loads (Spearman *ρ* = 0.39, *P* = 0.002; Fig. **3**j). There was a trend between Aβ(42)*56 and Braak Stage (Spearman *ρ* = 0.21, *P* = 0.10; Fig. **3**h).

Taken altogether, these findings suggest a potential link between Aβ*56 and global AD pathology.

### Levels of Aβ*56 are inversely associated with cognitive and memory function independently of Aβ plaque loads

Aβ*56 isolated from the Tg2576 mouse model of AD has been shown to impair memory when injected into the brains of healthy, wild-type rodents [51, 52, 57], indicative of its toxicity. To determine whether Aβ*56 may impair cognitive and memory function in humans, we analyzed the correlation between levels of Aβ*56 variants in the ITG and a set of neuropsychological assessments of memory and cognition. We found inverse correlations between levels of both Aβ*56 variants and the mini-mental state examination (MMSE) [22] scores (for Aβ(40)*56, Spearman *ρ* = −0.49, *P* <0.0001, Fig. **4**a; for Aβ(42)*56, Spearman *ρ* = −0.46, *P* = 0.0001, Fig. **4**h). In addition, we observed significant inverse correlations between both Aβ*56 variants and the z-scores of composite measures of episodic memory [99] (for Aβ(40)*56, Spearman *ρ* = −0.53, *P* <0.0001, Fig. **4**b; for Aβ(42)*56, Spearman *ρ* = −0.37, *P* = 0.003, Fig. **4**i), visuospatial ability/perceptual orientation [99] (for Aβ(40)*56, Spearman *ρ* = −0.32, *P* = 0.01, Fig. **4**c; for Aβ(42)*56, Spearman *ρ* = −0.29, *P* = 0.02, Fig. **4**j), perceptual speed [99] (for Aβ(40)*56, Spearman *ρ* = −0.32, *P* = 0.01, Fig. **4**d; for Aβ(42)*56, Spearman *ρ* = −0.36, *P* = 0.004, Fig. **4**k), semantic memory [99] (for Aβ(40)*56, Spearman *ρ* = −0.45, *P* = 0.0003, Fig. **4**e; for Aβ(42)*56, Spearman *ρ* = −0.42, *P* = 0.0007, Fig. **4**l), and global cognitive function [99] (for Aβ(40)*56, Spearman *ρ* = −0.47, *P* = 0.0001, Fig. **4**g; for Aβ(42)*56, Spearman *ρ* = −0.41, *P* = 0.001, Fig. **4**n). There was a significant inverse correlation between Aβ(42)*56 and working memory [99] (Spearman *ρ* = −0.29, *P* = 0.02; Fig. **4**m), but not between Aβ(40)*56 and working memory (Spearman *ρ* = −0.16, *P* = 0.22; Fig. **4**f).

**Fig. 4.**
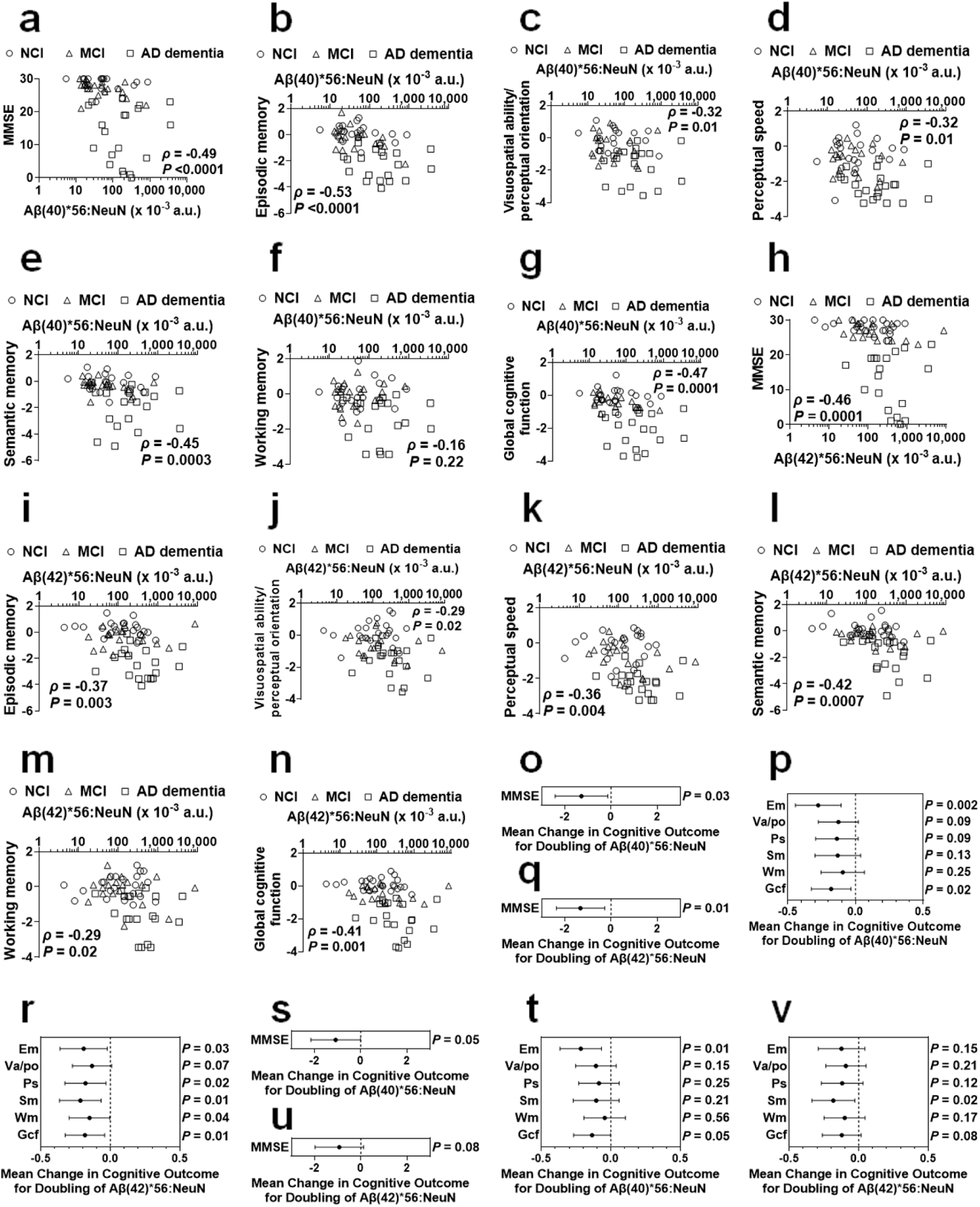
Levels of Aβ*56 variants correlate to degrees of cognitive and memory impairment independently of amyloid but not tau pathology. (**a, h**) Inverse correlations of levels of Aβ(40)*56 (a) and Aβ(42)*56 (h) (normalized to levels of NeuN) to the mini-mental state examination (MMSE) scores. (**b-g, i-n**) Correlations of NeuN-normalized levels of Aβ(40)*56 (b-g) and Aβ(42)*56 (i-n) to z scores of composite measures of episodic memory (b, i), visuospatial ability/perceptual orientation (c, j), perceptual speed (d, k), semantic memory (e, l), working memory (f, m), and global cognitive function (g, n). (**o, q**) Following adjustment for age at death, sex, years of education, and log-transformed Aβ plaque loads in the inferior temporal gyrus (ITG) using multiple linear regression (MLR), NeuN-normalized levels of Aβ(40)*56 (o) and Aβ(42)*56 (q) remain inversely correlated to MMSE scores. (**p, r**) Following adjustment for age at death, sex, years of education, and log-transformed Aβ plaque loads in the ITG using MLR, NeuN-normalized levels of Aβ(40)*56 (p) and Aβ(42)*56 (r) remain correlated to z scores of composite measures of episodic memory and global cognitive function. (**s-v**) Following adjustment for age at death, sex, years of education, and ptau 202/205-containing neurofibrillary tangle load in the ITG using MLR, NeuN-normalized levels of Aβ(40)*56 (s, t) and Aβ(42)*56 (u, v) are not correlated to MMSE scores (s, u) or z scores of composite measures of memory and cognition except for between Aβ(40)*56 and episodmic memory and between Aβ(42)*56 and semantic memory (t, v). For Figures a-n, Spearman’s rank-order correlations were used, and the x-axes are displayed on log scale. a.u., arbitrary units. NCI, no cognitive impairment; MCI, mild cognitive impairment; AD dementia, Alzheimer’s disease dementia. Em, episodic memory; Va/po, visuospatial ability/perceptual orientation; Ps, perceptual speed; Sm, semantic memory; Wm, working memory; Gcf, global cognitive function.

To control for possible effects of demographic characteristics and ITG Aβ plaque loads, we assessed the associations using MLR with adjustment for age at death, sex, years of education, and log-transformed ITG Aβ plaque load. A doubling of Aβ(40)*56 levels were associated with significantly lower mean MMSE score (mean change (95% CI): −1.28 (−2.41, −0.141), *P* = 0.03; Fig. **4**o), lower mean z-scores for episodic memory (−0.275 (−0.444, −0.106), *P* = 0.002; Fig. **4**p) and global cognitive function (−0.180 (−0.326, −0.0332), *P* = 0.02; Fig. **4**p), and trended lower mean z-scores for visuospatial ability/perceptual orientation (−0.126 (−0.274, 0.0212), *P* = 0.09; Fig. **4**p), perceptual speed (−0.137 (−0.294, 0.0205), *P* = 0.09; Fig. **4**p), and semantic memory (−0.131 (−0.300, 0.0385), *P* = 0.13; Fig. **4**p).

Similarly, a doubling of Aβ(42)*56 levels remained associated with significantly lower mean MMSE score (mean change (95% CI): −1.32 (−2.36, −0.27), *P* = 0.01; Fig. **4**q), lower mean z-scores for episodic memory (−0.192 (−0.362, −0.0217), *P* = 0.03; Fig. **4**r), perceptual speed (−0.179 (−0.327, −0.0315), *P* = 0.02; Fig. **4**r), semantic memory (−0.216 (−0.366, −0.0663), *P* = 0.01; Fig. **4**r), working memory (−0.150 (−0.296, −0.00452), *P* = 0.04; Fig. **4**r), and global cognitive function (−0.183 (−0.324, −0.0419), *P* = 0.01; Fig. **4**r), but not for visuospatial ability/perceptual orientation (−0.132 (−0.273, 0.00897), *P* = 0.07; Fig. **4**r). These results indicate that levels in the ITG of both Aβ*56 variants are inversely correlated with cognitive and memory function.

Interestingly, following adjustments for age at death, sex, years of education, and log-transformed ptau 202/205-containing NFT loads, we observed no significant associations between Aβ*56 variants and the seven neuropsychological variables except for between Aβ(40)*56 and episodic memory and between Aβ(42)*56 and semantic memory (Fig. **4**s-v). This suggests that pathological tau may mediate the association between Aβ*56 and cognitive and memory impairment.

### Comparison of the associations between AD-related pathology and cognitive and memory function

To assess the associations of varied AD-related pathological entities with cognition and memory function, we compared correlations between the neuropsychological test results and ITG Aβ plaque loads (Online Resource **1**: Figure S20), ITG ptau 202/205-containing NFT loads (Online Resource **1**: Figure S21), and NeuN-normalized levels of monomeric Aβ(1-40) and Aβ(1-42) in the ITG (Online Resource **1**: Figure S22) besides NeuN-normalized levels of Aβ*56 variants in the ITG (Fig. **4**). Note that since the cohorts used to measure Aβ(40)*56 and Aβ(42)*56 differed, we calculated Aβ plaque and NFT loads separately for each of the two cohorts. First, we compared the Spearman’s rank-order correlation coefficients of the four neuropathological and biochemical measures across all the seven types of neuropsychological scores without adjusting for demographic characteristics. We showed that levels of Aβ*56 variants and ptau 202/205-containing NFT loads exhibit overall stronger inverse correlations with memory and cognition scores than Aβ plaque loads and Aβ monomer levels (Online Resource **1**: Figure S23a and S23f). After adjustment for age at death, sex, and years of education, these observations persisted (Online Resource **1**: Figure S23b-e and S23g-j).

### Tau pathology mediates the association between Aβ*56 and cognitive and memory impairment

As the prevailing amyloid cascade hypothesis proposes that Aβ oligomerization—an early pathological process—instigates amyloid deposition and NFT formation leading eventually to dementia [30], our analyses shown above support that Aβ*56 may affect cognition and memory through pathological tau-mediated pathways independently of amyloid pathology.

To determine to what degree amyloid and tau pathology involves the impact of Aβ*56 on cognitive and memory function, we performed mediation analysis. We showed that ptau 202/205-containing NFT mediated 15-38% of the association between Aβ*56 levels and cognitive and memory scores. In particular, they had strong mediations in MMSE (mediated proportions: 29.8% (*P* = 0.03) and 28.1% (*P* = 0.05) for Aβ(40)*56 and Aβ(42)*56, respectively), episodic memory (mediated proportions: 28.6% (*P* = 0.02) and 38.1% (*P* = 0.03) for Aβ(40)*56 and Aβ(42)*56, respectively), and global cognitive function (mediated proportions: 33.6% (*P* = 0.03) and 32.9% (*P* = 0.02) for Aβ(40)*56 and Aβ(42)*56, respectively) (Fig. **5**, Online Resource **1**: Table S5 and S6). In contrast, Aβ plaques are a weak, non-significant mediator (mediated proportions: <18%) (Fig. **5**, Online Resource **1**: Table S7 and S8). These results further suggest that pathological tau rather than amyloid plaques mediates the association between Aβ*56 and cognitive and memory impairment.

**Fig. 5.**
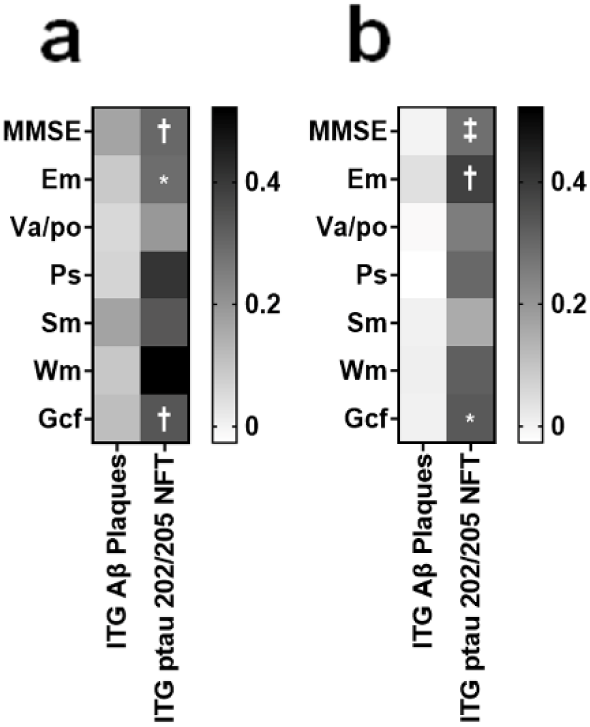
Tau pathology mediates the association between Aβ*56 and cognitive and memory impairment. Heat maps illustrating in the inferior temporal gyrus (ITG) the proportions of the associations between Aβ*56 variants (Aβ(40)*56 (**a**) and Aβ(42)*56 (**b**)) and the seven neuropsychological tests that are mediated by Aβ plaques (ITG Aβ Plaques) and ptau 202/205-containing neurofibrillary tangles (ITG ptau 202/205 NFT). MMSE, mini-mental state examination; Em, episodic memory; Va/po, visuospatial ability/perceptual orientation; Ps, perceptual speed; Sm, semantic memory; Wm, working memory; and Gcf, global cognitive function. The scales represent the percentages of effects mediated by either ITG Aβ Plaques or ITG ptau 202/205 NFT. * *P* = 0.02, † *P* = 0.03, ‡ *P* = 0.05.

## Discussion

In this study, we investigated the relationship between two Aβ*56 variants—Aβ(40)*56 and Aβ(42)*56—and cognitive function in AD. Our analysis focused on the ITG, an area affected early in AD [80]. We found that both variants were significantly elevated in individuals with AD dementia and showed an inverse correlation with various measures of memory and cognition. Notably, after adjusting for amyloid pathology, Aβ(40)*56 remained significantly elevated, while Aβ(42)*56 exhibited a strong trend toward elevation. The relevance of Aβ*56 to neurological function was further underscored by the finding that its correlations with cognitive measures were stronger than those for amyloid plaques ‒ the defining hallmark of AD pathology. These results highlight Aβ*56 as a potential key contributor to memory and cognitive impairment independently of amyloid plaques.

The results are clinically significant because they suggest that patients receiving antibodies that reduce amyloid plaques, such as lecanemab [19] and donanemab [18, 87], could potentially achieve additional cognitive benefit by targeting Aβ*56. Currently, although monoclonal antibodies that specifically bind Aβ*56 are lacking, such antibodies could be useful in testing this hypothesis.

The differential binding of various antibodies to distinct Aβo arises from structural differences. Aβo can be broadly classified into two types: fibrillar Aβo and non-fibrillar Aβo [58]. Fibrillar Aβo, which contain in-register β-sheets ‒ a hallmark of brain-derived Aβ fibrils, are recognized by conformation-sensitive OC antibodies [40, 58]. In contrast, non-fibrillar Aβo, such as Aβ*56, are recognized by A11 antibodies that target unique conformational epitopes absent in fibrillar Aβo, although these epitopes are still not fully defined [39, 41, 48, 54, 58]. Current imaging tools such as amyloid positron emission tomography, plasma biomarkers such as Aβ(x-42), and recently approved anti-Aβ therapies target fibrillar Aβo and/or Aβ fibrils [16, 18, 19, 81, 87]. However, they do not address non-fibrillar Aβo. Encouraging preliminary results from studies using ACU193 antibodies [45] and cyclic D,L-α-peptides [77], which appear to target structures other than fibrils, indicate that therapies targeting non-fibrillar Aβo may be on the horizon [28, 46].

We initially expected to observe higher levels of Aβ*56 in individuals with MCI compared to those with NCI, given that Aβo are thought to play an early role in AD [15]. Surprisingly, we found no significant difference in Aβ*56 levels between the two groups. We believe this may be due to the neurobiological processes driving MCI being more pronounced in the mesial temporal lobe, rather than the neocortex where the ITG—the structure we studied—is located. This is supported by our findings that, in our cohort, levels of NFT (Online Resource **1**: Figure S15) and Aβ plaques (Fig. **2**a and d) in the ITG were similar between NCI and MCI groups. Another possible explanation relates to our selection criteria for NCI samples. To match amyloid pathology across groups, we included NCI individuals with relatively high amyloid pathology. These individuals are more likely to progress to MCI [29, 72], making it plausible that the NCI participants in our cohort were closer to transitioning to MCI and, as a result, had Aβ*56 levels comparable to those in the MCI group.

It is well known that Aβo can promote tau hyperphosphorylation and dysregulate various signaling pathways malfunctioning in AD (for example, see [17, 47, 61, 76, 91, 104]). In addition, accumulating evidence supports a strong association between pathological tau phosphorylation and cognitive deficits in AD dementia [100]. While the exact identities of brain-derived oligomeric Aβ assemblies that elicit tau hyperphosphorylation remain unclear, a recent study showed that soluble brain Aβ protofibrils predict CSF tau pathology better than amyloid plaques [4]. Our findings, for the first time, suggest the effects of Aβ*56—a non-plaque-dependent type of Aβo—is associated with cognitive and memory dysfunction, and ptau 202/205 may mediate such association. The precise biological mechanisms linking Aβ*56 to tau remain unclear, and further research is needed to elucidate this interaction.

There are conflicting published reports on the association between Aβ*56 and AD dementia. One study reported that Aβ*56 in nasal fluid is elevated in patients with AD and that high levels predict more rapid cognitive decline [103]. Our current findings align with this result. However, another study, by Lesné *et al.*, found that Aβ*56 levels in the ITG gradually increased with normal aging but declined in AD dementia [53], which our results do not support. This discrepancy likely stems from differences in experimental settings used in the two studies. Key methodological differences include the use of detergents in Lesné *et al.* to extract brain proteins, specifically 0.01% (v/v) octyl phenoxypolyethoxylethanol (also known as nonidet P-40) and 0.1% (w/v) SDS for the extracellular-enriched (EC) protein fraction; and 0.5% (v/v) Triton X-100, 3% (w/v) SDS, and 1% (w/v) deoxycholate for the membrane-associated protein fraction. Lesné *et al.* also relied on non-Aβ-specific antibodies, specifically monoclonal antibody 6E10 targeting Aβ(3-8) and monoclonal antibody 4G8 targeting Aβ(17-24) and used straight WB to detect proteins. In contrast, our study used detergent-free, aqueous buffers for protein extraction and detected Aβ*56 using IP/WB with antibodies that specifically recognize the C- and N- termini of canonical Aβ peptides. While it is likely that Lesné *et al.* measured *bona fide* Aβ*56 entities, supported by our detection of Aβ*56 in EC protein fractions [56] and human cerebrospinal fluid (CSF) [27], it is possible that their EC protein fractions also included N-terminally extended hAPP fragments, such as those found in CSF by mass spectrometry [27]. Such fragments cannot be distinguished from Aβ*56 using non-Aβ-specific antibodies, contributing to the observed differences between the two studies.

Distinct from our studies on Aβ*56, multiple independent studies have linked various other Aβo to AD dementia. For instance, Esparza *et al.* used an enzyme-linked immunosorbent assay (ELISA) with an Aβ(1-x)-specific antibody for both capture and detection, showing elevated levels of Aβ-related entities in aqueous cortical brain homogenates from AD patients compared to cognitively unimpaired controls with matched levels of amyloid pathology [21]. However, due to the experimental setup, it was unclear whether the ELISA-detected signals were exclusively from Aβo and how fibrillar and non-fibrillar oligomers each contributed to the measurements [20]. Similarly, Bilousova *et al.* employed an ELISA with an Aβ(5-13)-specific antibody and examined Aβo in synaptic terminals of the parietal cortex. They found elevated levels of Aβo—comprising both OC- and A11-reactive entities—in AD patients compared to non-demented controls with comparable amyloid pathology [9]. This suggests that both fibrillar and non-fibrillar Aβo may contribute to disease. Tomic *et al.* used immunoblots with conformation-sensitive antibodies to show that OC-reactive fibrillar oligomers, but not A11-reactive non-fibrillar oligomers, are elevated in multiple brain regions of individuals with AD dementia. These elevations correlate with cognitive decline as well as amyloid and tau pathology [90]. However, the broad binding specificity of the antibodies used [39–41] limits the ability to assess the specific contribution of fibrillar Aβo to signals. Additionally, the low sample size (*N* = 2-6 per diagnostic group) in this study reduces the statistical power of its findings.

Aβ dimers have also been implicated in AD. Immune-based assays have associated Aβ dimers in the brain [64, 84] and blood [94] with AD dementia. While free Aβ dimers appear in blood [94], several findings suggest that Aβ dimers in the brain are closely linked to amyloid fibrils and serve as surrogate markers for Aβ fibrils and fibrillar oligomers rather than independent entities. These include the observations that they are present exclusively in high-molecular-weight SEC fractions of aqueous brain homogenates, are released from neuritic plaque cores by formic acid [84], and rapidly assemble into fibrillar intermediates *in vitro* [71].

Savage *et al.* demonstrated elevated levels of Aβ-derived diffusible ligands (ADDLs) in CSF of AD patients using an ELISA that used the conformation-sensitive 19.3 (ACU193) antibody, which does not bind thioflavin-S-labeled dense-core plaques, to capture and the 82E1 antibody to detect ADDLs [46]. They also found an inverse correlation between ADDLs and MMSE scores [79]. The 19.3 antibody blocks deposition of ADDLs on Aβ plaques in AD mouse brains [75] and dose-dependently reduces Aβ plaques [66], indicating a potential relationship of ADDLs with amyloid plaques.

Previous studies have established a robust link between soluble Aβ and cognitive dysfunction in AD dementia [60, 65, 96]. Consistent with these findings, we found that levels of monomeric Aβ—the primary component of the soluble Aβ pool—correlate with the severity of cognitive dysfunction (Online Resource **1**: Figure S22a-n). However, distinct from Aβ*56, these correlations largely disappear after adjusting demographic features and amyloid plaques (Online Resource **1**: Figure S22o-r). Additionally, we found a strong correlation between Aβ*56 and Aβ monomers (Online Resource **1**: Figure S24), suggesting a dynamic equilibrium between these two entities.

Of note, we found that Aβ*56 levels are higher in individuals with *APOE ɛ3ɛ4* compared to those with *APOE ɛ3ɛ3* (Online Resource **1**: Figure S25). This aligns with previous findings showing that Aβ(x-40) aggregates increase in an *APOE ɛ4* dose-dependent manner in AD patients [24, 34, 62]. Furthermore, in AD brains, Aβo colocalize with apolipoprotein E proteins in synaptic terminals and are more strongly associated with synaptic loss in *APOE ɛ4* carriers compared with *APOE ɛ3* carriers [42]. Further studies are needed to clarify the specific roles of various Aβo in these processes.

This study has four main strengths. First, we measured a specific type of Aβo using techniques that avoid detecting soluble hAPP fragments. Second, we adjusted the associations between Aβ*56 and clinical status for demographic features and amyloid burden. Third, we examined the relationships between Aβ*56 and a comprehensive panel of neuropsychological measures. Fourth, we showed all WB images used in this study (Online Resource **1**: Figure S4, S5, S7, S8, and S10-S13) and provided comprehensive specifications for every lane of each blot (Online Resource **1**: Table S3). These elements of the study design and reporting enhance the reliability, reproducibility, and significance of the results.

However, the study also has two main limitations. First, we focused exclusively on a small specimen (0.1-0.8 g) in a single brain region, the ITG. While this limitation can be addressed in future studies by examining other brain regions and biofluids, it is notable that Aβ*56 measurements from discrete specimens comprising less than 1% of the brain are strongly associated with global clinical and neuropathological features of AD. Second, the study cohort primarily consisted of non-Hispanic white individuals [7] in which the prevalence of amyloid plaques is approximately 2-3 times higher than in Asians, Black individuals, and Hispanics [69]. Since amyloid positivity is a prerequisite for current anti-Aβ therapies [86, 92], white individuals disproportionately benefit from these treatments, despite a similar or higher prevalence of dementia in other racial and ethnic groups [44, 50, 63]. Future studies should investigate the distribution of non-fibrillar Aβo, such as Aβ*56, across diverse racial and ethnic populations. Understanding the pathological role of non-fibrillar Aβo could pave the way for developing disease-modifying therapies that benefit a broader and more representative population.

## Conclusions

In conclusion, our study demonstrated that a specific type of non-fibrillar, plaque-independent Aβo, Aβ*56, is elevated in AD dementia and inversely correlated with multiple domains of memory and cognition. These associations are influenced by NFT, but only minimally by amyloid plaques. Our findings are significant because they suggest that targeting Aβ*56 could provide additional therapeutic benefits beyond those achieved by current anti-Aβ therapies that focus on removing amyloid plaques.

## List of abbreviations

Aβ: β-amyloid
Aβo: β-amyloid oligomers
Aβ(1-40): 40-amino-acid-long β-amyloid peptides with the N-terminus amino acid being aspartate-672 and the C-terminus amino acid being valine-711 (human amyloid precursor protein 770 isoform numbering)
Aβ(1-42): 42-amino-acid-long β-amyloid peptides with the N-terminus amino acid being aspartate-672 and the C-terminus amino acid being alanine-713 (human amyloid precursor protein 770 isoform numbering)
Aβ(x-40): β-amyloid-sequence-containing (poly)peptides with the C-terminus ending at valine-711 (human amyloid precursor protein 770 isoform numbering)
Aβ(x-42): β-amyloid-sequence-containing (poly)peptides with the C-terminus ending at alanine-713 (human amyloid precursor protein 770 isoform numbering)
Aβ*56: a brain-derived type of memory-impairing β-amyloid oligomers that are water-soluble, stable when exposed to sodium dodecyl sulfate, apparently ∼56-kDa in (semi-)denaturing sodium dodecyl sulfate-polyacrylamide gel electrophoresis and in non-denaturing size exclusion chromatography, reactive to anti-oligomer antibody A11, and independent of amyloid plaques
Aβ(40)*56: a variant of Aβ*56 that contains canonical Aβ(1-40)
Aβ(42)*56: a variant of Aβ*56 that contains canonical Aβ(1-42)
AD: Alzheimer’s disease
ADDLs: Aβ-derived diffusible ligands
*APOE*: apolipoprotein E gene
APP/TTA: a bigenic transgenic mouse strain B6.Cg-Tg(tetO-APPSwInd)102Dbo/Mmjax in which the human amyloid precursor protein 695 isoform with the Swedish (K670N, M671L) and Indiana (V717F) mutations linked to familial Alzheimer’s disease is overexpressed under the driven of an engineered tetracycline-responsive promoter (RRID:MMRRC_034846-JAX)
BCA: bicinchoninic acid
Bicine: *N*,*N*-Bis(2-hydroxyethyl)glycine
Bis-tris methane: 2-[Bis(2-hydroxyethyl)amino]-2-(hydroxymethyl)propane-1,3-diol BME β-mercaptoethanol
Cat. No.: catalog number
CERAD: Consortium to Establish a Registry for Alzheimer’s Disease
CI(s): confidence interval(s)
CSF: cerebrospinal fluid
DynaG: Dynabeads Protein G
EC: extracellular-enriched
EDTA: ethylenediaminetetraacetic acid
ELISA: enzyme-linked immunosorbent assay
GAPDH: glyceraldehyde 3-phosphate dehydrogenase
GuHCl: guanidine hydrochloride
hAPP: human amyloid precursor protein
HCl: hydrochloric acid
HRP: horseradish peroxidase
IgG: immunoglobulin G
IP: immunoprecipitation
ITG: inferior temporal gyrus
KCl: potassium chloride
kDa: kilo-Dalton
MAP: Memory Aging Project
MCI: mild cognitive impairment
MLR(s): multiple linear regression(s)
MMSE: mini-mental state examination
NA: NeutrAvidin
NaCl: sodium chloride
NCI: no cognitive impairment
NeuN: neuronal nuclei
NFT: neurofibrillary tangles
NH_4_HCO_3_: ammonium bicarbonate
NIA-Reagan: National Institute on Aging-Reagan
NIH: National Institutes of Health
OTG: *n*-Octyl-1-thio-*β*-*D*-glucopyranoside
PAGE: polyacrylamide gel electrophoresis
PBS: phosphate-buffered saline
phen: 1,10-phenanthroline monohydrate
PMI: post-mortem interval
PMSF: phenylmethylsulfonyl fluoride
ptau 202/205: phosphorylated tau at serine 202 and threonine 205
ROS: Religious Orders Study
RRID: Research resource identifier
SDS: sodium dodecyl sulfate
SEC: size exclusion chromatography
Tg2576: transgenic mouse strains B6;SJL-Tg(APPSWE)2576Kha/Tac, 129S6.Cg-Tg(APPSWE)2576Kha/Tac, and 129S6;FVBF1-Tg(APPSWE)2576Kha that overexpress human amyloid precursor protein 695 isoform with the Swedish (K670N, M671L) mutation linked to familial Alzheimer’s disease
Tricine: *N*-[1,3-Dihydroxy-2-(hydroxymethyl)propan-2-yl]glycine
Tris: tris(hydroxymethyl)aminomethane
Triton X-100: polyethylene glycol p-(1,1,3,3-tetramethylbutyl)-phenyl ether
Tween 20: polyoxyethylene (20) sorbitan monolaurate
v/v: volume/volume
WB: western blotting
w/v: weight/volume

## Declarations

### Ethics approval and consent to participate

Informed consent for clinical and neuropathologic evaluation and autopsy was obtained from all individual participants. All procedures of using de-identified human brain tissue, obtained at autopsy, were reviewed and approved by the Institutional Review Boards of the Rush University and the University of Minnesota.

### Consent for publication

Not applicable.

### Availability of data and material

All data generated or analyzed during this study are included in this published article and its supplementary information files.

## Acknowledgements

The authors thank Drs. David Bennett and Gregory Klein at the Rush University, Chicago, Illinois for providing human brain specimens.

