## Supporting information for "The β-amyloid oligomer Aβ*56 is associated with Alzheimer’s dementia independently of amyloid pathology"

*Acta Neuropathologica (Original Articles)*

Corresponding author: Peng Liu

This document contains 25 supplementary figures, 8 supplementary tables, and supplementary references.

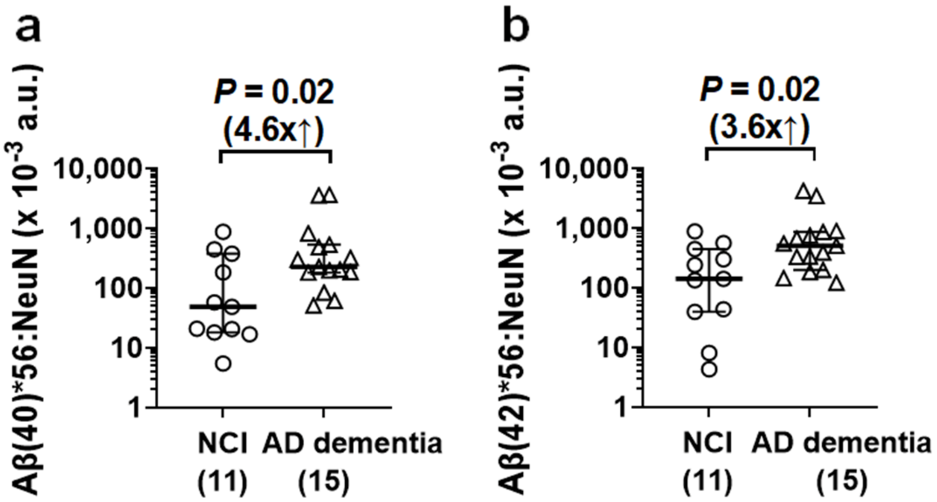
a
Aβ(40)*56:NeuN (x 10-3 a.u.)
P = 0.02
(4.6x↑)
10,000
1,000
100
10
1
NCI
(11)
AD dementia
(15)
b
Aβ(42)*56:NeuN (x 10-3 a.u.)
P = 0.02
(3.6x↑)
10,000
1,000
100
10
1
NCI
(11)
AD dementia
(15)

**Figure S1. Levels of Aβ*56 variants are elevated in the inferior temporal gyri of individuals with Alzheimer's disease (AD) dementia compared to individuals with no cognitive impairment (NCI) – A small cohort study.** Levels of Aβ(40)*56 and Aβ(42)*56, after normalized to levels of neuronal nuclei (NeuN), a protein marker for post-mitotic neurons, in individuals with AD dementia are 4.6- and 3.6-fold higher (in both cases, $U = 38$; $P = 0.02$; two-tailed Mann-Whitney *U* tests) than individuals with NCI, respectively. The scatter plots represent data of individuals plus group median ± interquartile range. The y-axes in figures are displayed on log scale. The numbers of individuals used for the analysis are shown in parentheses. a.u., arbitrary units.

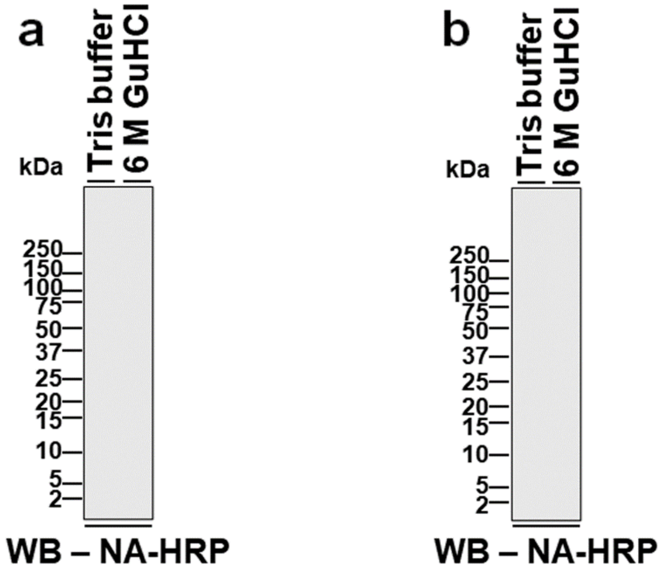
a
Tris buffer
6 M GuHCl
kDa
250
150
100
75
50
37
25
20
15
10
5
2
WB – NA-HRP
b
Tris buffer
6 M GuHCl
kDa
250
150
100
75
50
37
25
20
15
10
5
2
WB – NA-HRP

**Figure S2. Immunopurified ~56-kDa entities are not detected without anti-oligomer A11 antibodies.**

Representative western blotting (WB) digital graphs show that no D8Q7I (**a**)- or D3E10 (**b**)-purified, electro-elution-isolated ~56-kDa entities from AD brains are detected using horseradish peroxidase-conjugated NeutrAvidin (NA-HRP) following the treatment of the ~56-kDa entities by Tris buffer or 6 M guanidine hydrochloride (GuHCl). The exposure times are 3 sec (a) and 1 sec (b), which are the same as the exposure times shown in Figures 1f and g, respectively. kDa, kilo-Daltons.

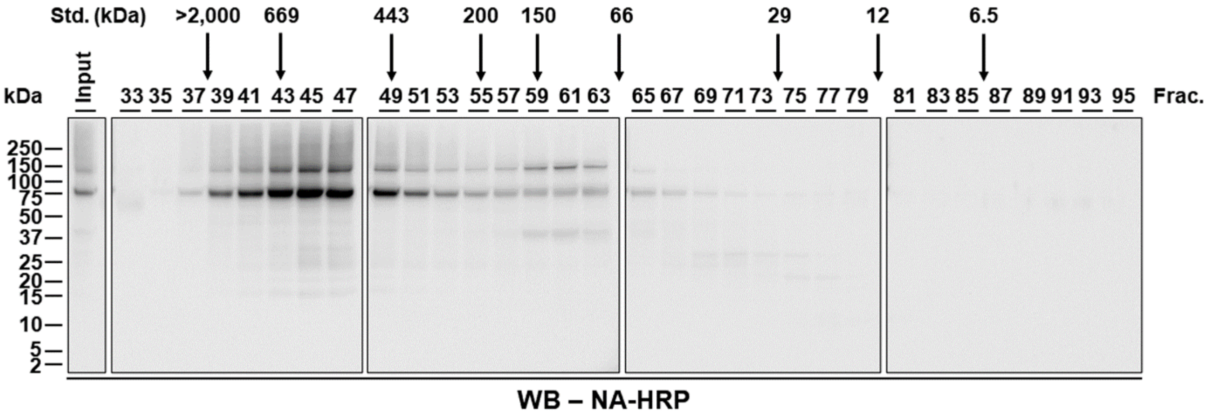
Std. (kDa)
>2,000
669
443
200
150
66
29
12
6.5
kDa
Input
33 35 37 39 41 43 45 47
49 51 53 55 57 59 61 63
65 67 69 71 73 75 77 79
81 83 85 87 89 91 93 95
Frac.
250
150
100
75
50
37
25
20
15
10
5
2
WB – NA-HRP

**Figure S3. The ~56-kDa entities in AD brains are not detected in fractions of size exclusion chromatography (SEC) without anti-β-amyloid (Aβ) antibody.**

Representative western blotting (WB) digital graphs showing the detection of entities in odd-numbered SEC fractions (Frac. 33-95) of aqueous brain extracts from individuals with AD dementia using horseradish peroxidase-conjugated NeutrAvidin (NA-HRP). No ~56-kDa entities, ~14-kDa entities, ~9-kDa entities, or monomeric Aβ are detected. The four WB graphs (*i.e.*, Frac. 33-47, 49-63, 65-79, and 81-95) shown were obtained using the same exposure time (5 sec) as the graphs shown in the upper panels of Figure 1h. The WB patterns and band intensities of the input (2.5% (w/v) of the extracts used for SEC) of the four WB graphs are comparable; a WB graph of the input is shown in the very left panel. The elution profile of a series of biomolecule standards (Std.) is shown above the WB graphs. The detection agent NA-HRP was stripped and then probed with biotinylated 82E1 to obtain graphs shown in Figure 1h. kDa, kilo-Daltons.

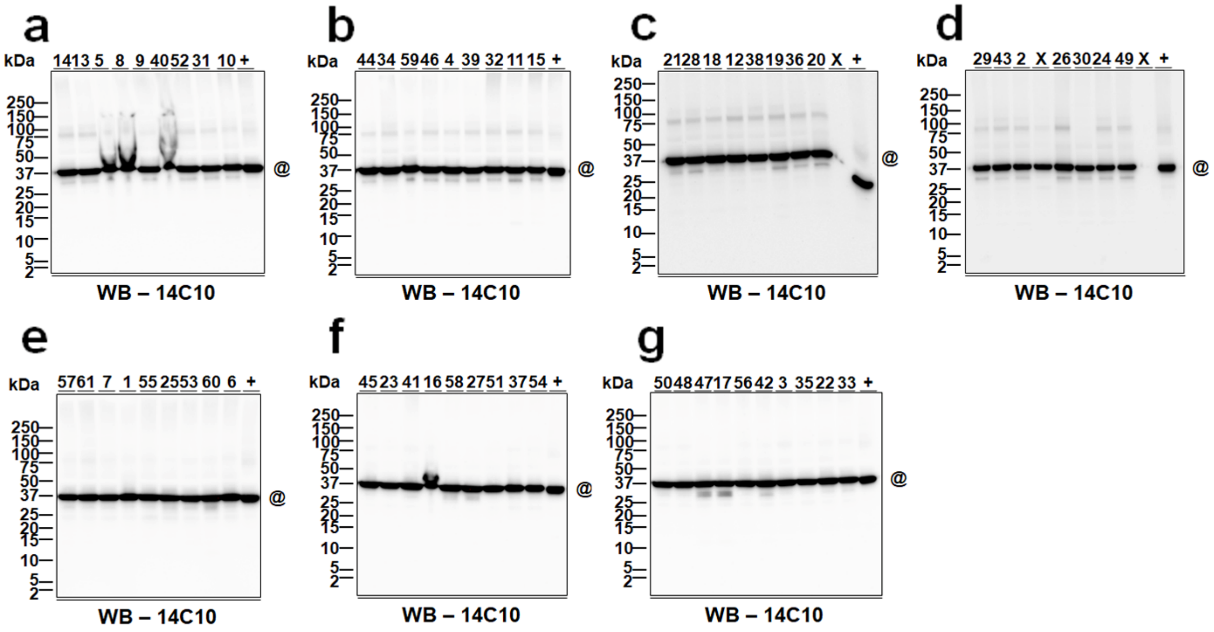
a
b
c
d
e
f
g
kDa
14 13 5 8 9 40 52 31 10 +
44 34 59 46 4 39 32 11 15 +
21 28 18 12 38 19 36 20 X +
29 43 2 X 26 30 24 49 X +
57 61 7 1 55 25 53 60 6 +
45 23 41 16 58 27 51 37 54 +
50 48 47 17 56 42 3 35 22 33 +
250
150
100
75
50
37
25
20
15
10
5
2
@
WB – 14C10

**Figure S4. The detection of housekeeping protein glyceraldehyde 3-phosphate dehydrogenase (GAPDH) in the individuals for measuring Aβ(40)*56. a-g**) Western blotting (WB) digital images used for the detection and quantification of GAPDH (@). IDs of individuals (also see Online Resource **1**: Table S1) are shown above the images. +, aqueous brain extracts of 4-5-month-old Tg2576 mice (serving as an internal control for WB, Online Resource **1**: Table S4). WB – 14C10, anti-GAPDH antibody clone 14C10 serving as the detection antibody for WB. These images were collected by stripping the detecting antibody 27-4 of the corresponding blots in Figure S7 and reprobing with 14C10. Of note, Figures a-d are the same images as Figures a-d in Figure S5. This is because individuals used to measure Aβ(40)*56 and Aβ(42)*56 partially overlapped with 11 individuals with no cognitive impairment, 8 individuals with mild cognitive impairment, and 15 individuals with AD dementia used in both cohorts. kDa, kilo-Daltons.

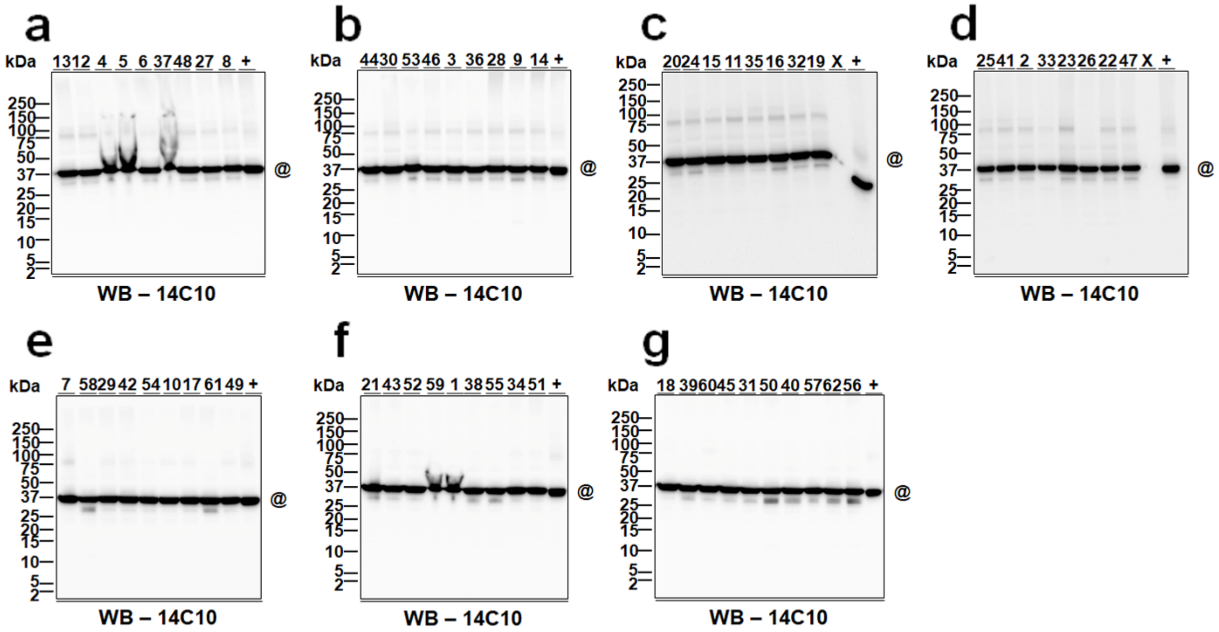
a
b
c
d
e
f
g
kDa
250
150
100
75
50
37
25
20
15
10
5
2
@
WB – 14C10

**Figure S5. The detection of housekeeping protein glyceraldehyde 3-phosphate dehydrogenase (GAPDH) in the individuals for measuring Aβ(42)*56. a-g**) Western blotting (WB) digital images used for the detection and quantification of GAPDH (@). IDs of individuals (also see Online Resource **1**: Table S2) are shown above the images. +, aqueous brain extracts of 4-5-month-old Tg2576 mice (serving as an internal control for WB, Online Resource **1**: Table S4). WB – 14C10, anti-GAPDH antibody clone 14C10 serving as the detection antibody for WB. These images were collected by stripping the detecting antibody 27-4 of the corresponding blots in Figure S8 and reprobing with 14C10. Of note, Figures a-d are the same images as Figures a-d in Figure S4. This is because individuals used to measure Aβ(40)*56 and Aβ(42)*56 partially overlapped with 11 individuals with no cognitive impairment, 8 individuals with mild cognitive impairment, and 15 individuals with AD dementia used in both cohorts. kDa, kilo-Daltons.

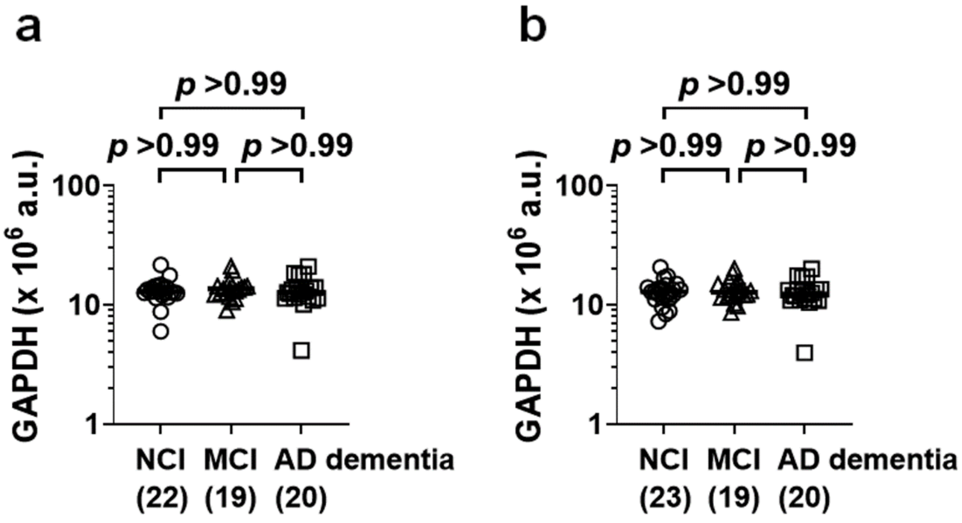
a
p >0.99
p >0.99
p >0.99
GAPDH (x 10^6 a.u.)
100
10
1
NCI (22)
MCI (19)
AD dementia (20)
b
p >0.99
p >0.99
p >0.99
GAPDH (x 10^6 a.u.)
100
10
1
NCI (23)
MCI (19)
AD dementia (20)

**Figure S6. Levels of housekeeping protein glyceraldehyde 3-phosphate dehydrogenase (GAPDH) are comparable in the inferior temporal gyri of individuals with no cognitive impairment (NCI), mild cognitive impairment (MCI), and Alzheimer's disease (AD) dementia.** (**a, b**) For the Aβ(40)*56 study, following a Kruskal-Wallis test ($H$ (2) = 0.82, $P$ = 0.67), *post hoc* analyses show no difference in levels of GAPDH between individuals with NCI, MCI, and AD dementia (a). In parallel, for the Aβ(42)*56 study, following a Kruskal-Wallis test ($H$ (2) = 0.22, $P$ = 0.90), *post hoc* analyses show no difference in levels of GAPDH between individuals with NCI, MCI, and AD dementia (b). The scatter plots represent data of individuals plus group median ± interquartile range. The y-axes in figures are displayed on log scale. The numbers of individuals with NCI, MCI, and AD dementia used are shown in parentheses. a.u., arbitrary units.

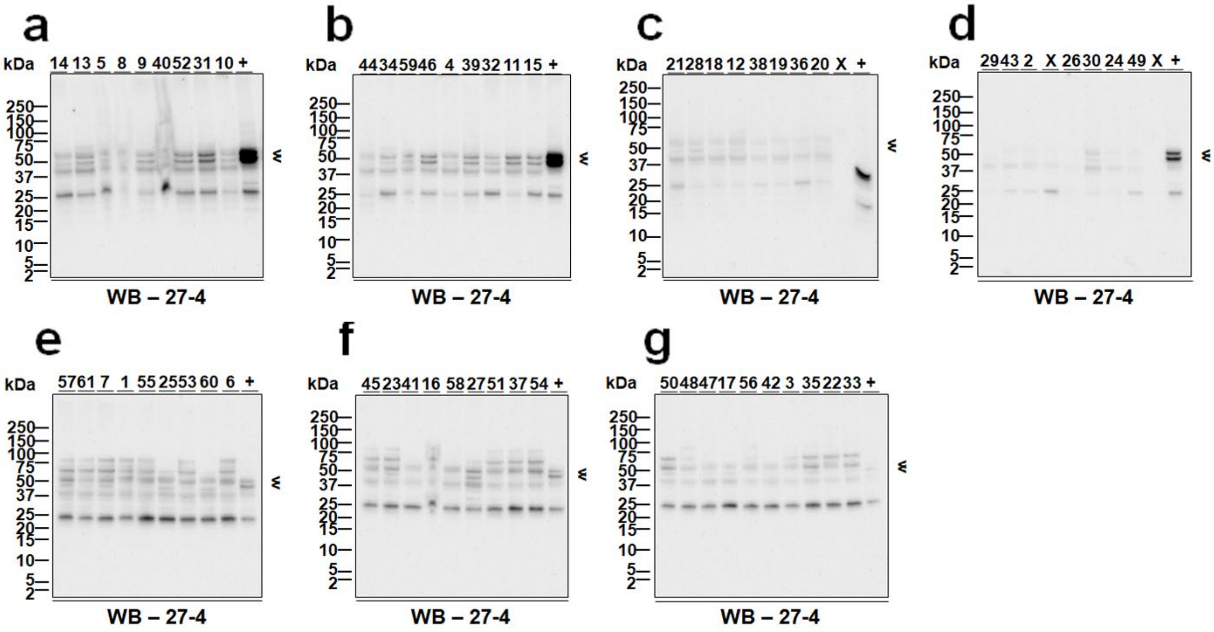
a
kDa
14 13 5 8 9 40 52 31 10 +
250
150
100
75
50
37
25
20
15
10
5
2
WB – 27-4
b
kDa
44 34 59 46 4 39 32 11 15 +
WB – 27-4
c
kDa
21 28 18 12 38 19 36 20 X +
WB – 27-4
d
kDa
29 43 2 X 26 30 24 49 X +
WB – 27-4
e
kDa
57 61 7 1 55 25 53 60 6 +
WB – 27-4
f
kDa
45 23 41 16 58 27 51 37 54 +
WB – 27-4
g
kDa
50 48 47 17 56 42 3 35 22 33 +
WB – 27-4

**Figure S7. The detection of neuronal marker NeuN in the individuals for measuring Aβ(40)*56. a-g**) Western blotting (WB) digital images used for the detection and quantification of NeuN (arrowheads). IDs of individuals (also see Online Resource **1**: Table S1) are shown above the images. +, aqueous brain extracts of 4-5-month-old Tg2576 mice (serving as an internal control for WB, Online Resource **1**: Table S4). WB – 27-4, anti-NeuN antibody clone 27-4 serving as the detection antibody for WB. Of note, Figures a-d are the same images as Figures a-d in Figure S8. This is because individuals used to measure Aβ(40)*56 and Aβ(42)*56 partially overlapped with 11 individuals with no cognitive impairment, 8 individuals with mild cognitive impairment, and 15 individuals with AD dementia used in both cohorts. kDa, kilo-Daltons.

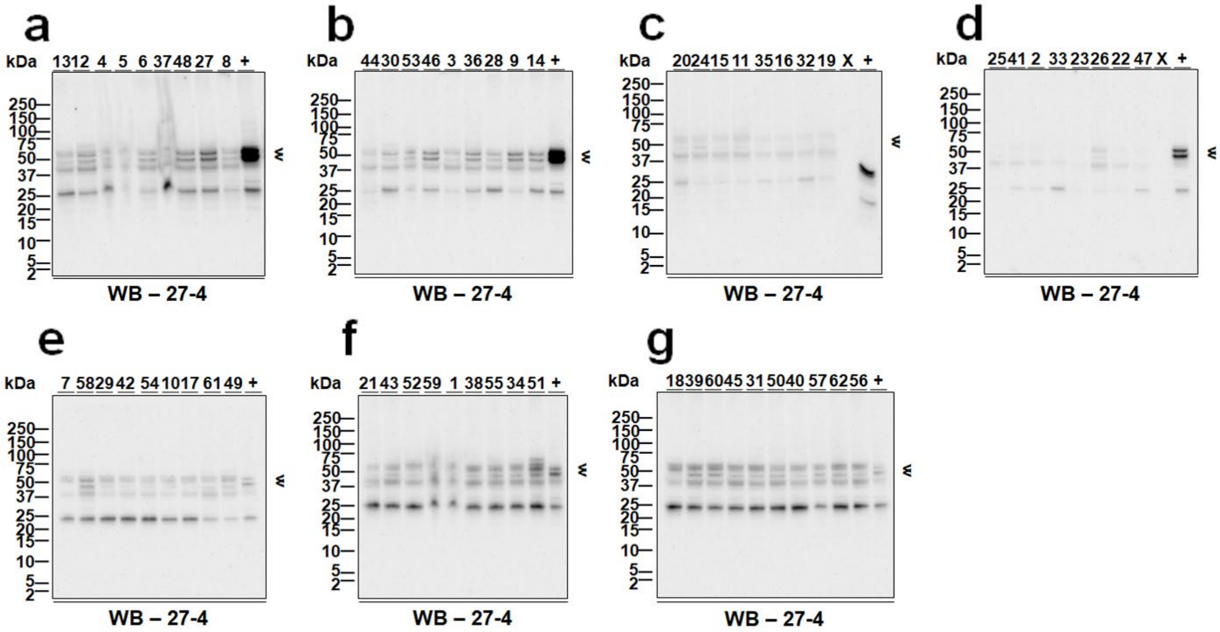
a
b
c
d
e
f
g
kDa
250
150
100
75
50
37
25
20
15
10
5
2
13 12 4 5 6 37 48 27 8 +
44 30 53 46 3 36 28 9 14 +
20 24 15 11 35 16 32 19 X +
25 41 2 33 23 26 22 47 X +
7 58 29 42 54 10 17 61 49 +
21 43 52 59 1 38 55 34 51 +
18 39 60 45 31 50 40 57 62 56 +
WB – 27-4

**Figure S8. The detection of neuronal marker NeuN in the individuals for measuring Aβ(42)*56. a-g**) Western blotting (WB) digital images used for the detection and quantification of NeuN (arrowheads). IDs of individuals (also see Online Resource **1**: Table S2) are shown above the images. +, aqueous brain extracts of 4-5-month-old Tg2576 mice (serving as an internal control for WB, Online Resource **1**: Table S4). WB – 27-4, anti-NeuN antibody clone 27-4 serving as the detection antibody for WB. Of note, Figures a-d are the same images as Figures a-d in Figure S7. This is because individuals used to measure Aβ(40)*56 and Aβ(42)*56 partially overlapped with 11 individuals with no cognitive impairment, 8 individuals with mild cognitive impairment, and 15 individuals with AD dementia used in both cohorts. kDa, kilo-Daltons.

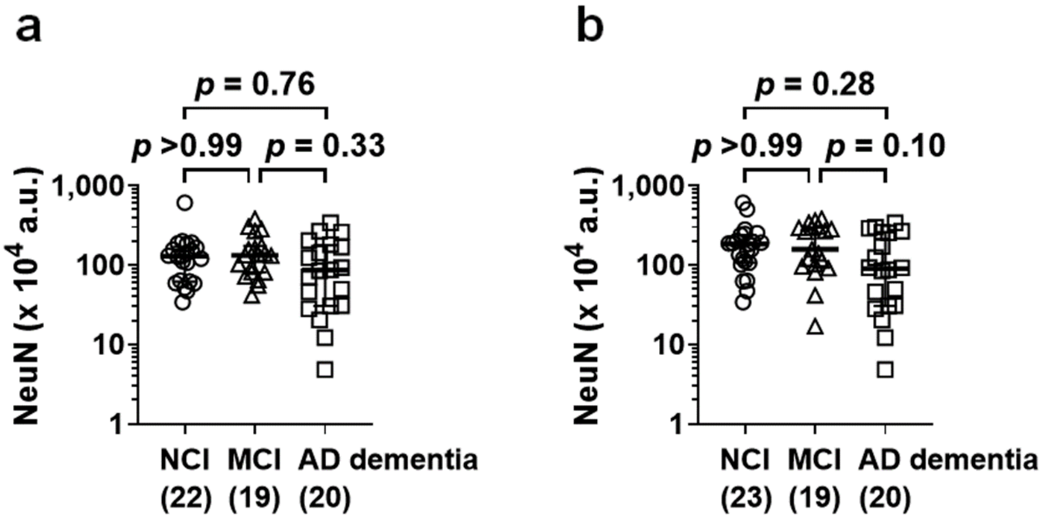
a
p = 0.76
p >0.99
p = 0.33
NeuN (x 10^4 a.u.)
1,000
100
10
1
NCI (22)
MCI (19)
AD dementia (20)
b
p = 0.28
p >0.99
p = 0.10
NeuN (x 10^4 a.u.)
1,000
100
10
1
NCI (23)
MCI (19)
AD dementia (20)

**Figure S9. Levels of neuronal marker NeuN are comparable in the inferior temporal gyri of individuals with no cognitive impairment (NCI), mild cognitive impairment (MCI), and Alzheimer's disease (AD) dementia.** (**a, b**) For the Aβ(40)*56 study, following a Kruskal-Wallis test (*H* (2) = 2.71, *P* = 0.26), *post hoc* analyses show no difference in levels of NeuN between individuals with NCI, MCI, and AD dementia (a). In parallel, for the Aβ(42)*56 study, following a Kruskal-Wallis test (*H* (2) = 4.99, *P* = 0.08), *post hoc* analyses show no difference in levels of NeuN between individuals with NCI, MCI, and AD dementia (b). The scatter plots represent data of individuals plus group median ± interquartile range. The y-axes in figures are displayed on log scale. The numbers of individuals with NCI, MCI, and AD dementia used are shown in parentheses. a.u., arbitrary units.

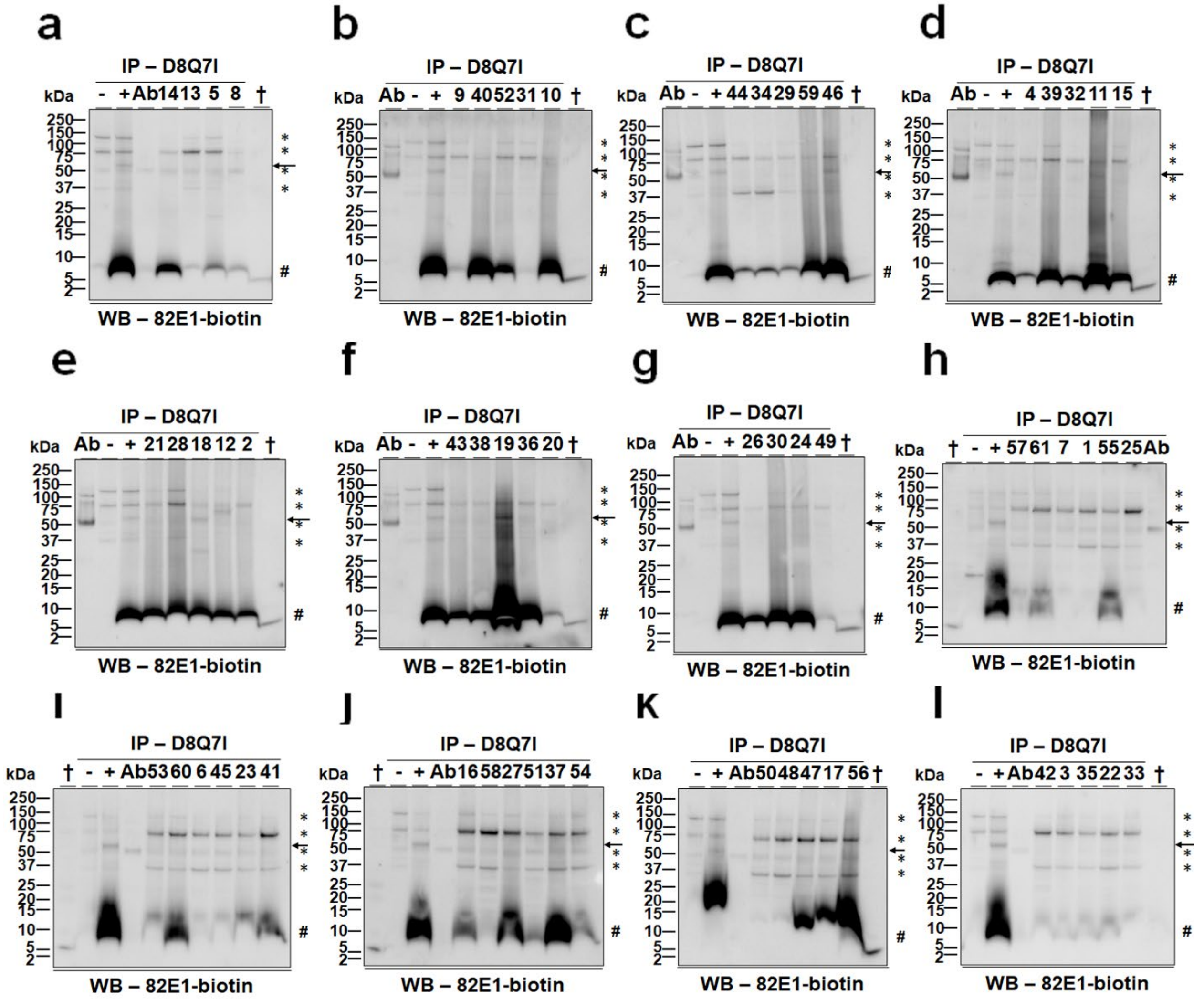

a
IP – D8Q7I
WB – 82E1-biotin
b
c
d
e
f
g
h
I
J
K
l

**Figure S10. The detection of Aβ(40)*56 in the studied individuals. a-l**) Immunoprecipitation (IP)/western blotting (WB) digital images used for the detection and quantification of Aβ(40)*56 (arrow). IDs of individuals (also see Online Resource **1**: Table S1) are shown above the images. +, IP reactions with the D8Q7I-immobilized DynaG matrix and aqueous brain extracts of 10-11-month-old APP/TTA mice (serving as a positive control for IP/WB, Online Resource **1**: Table S4). -, IP reactions with the D8Q7I-immobilized DynaG matrix and aqueous brain extracts of 10-11-month-old littermates of APP/TTA mice that did not express human amyloid precursor proteins (serving as a negative control for IP/WB, Online Resource **1**: Table S4). Ab, IP reactions with the D8Q7I-immobilized DynaG matrix but no brain extracts. †, synthetic Aβ(1-40) (2 ng), serving as a positive control for WB. IP – D8Q7I, IP capture antibody D8Q7I that is directed against the C-terminus of Aβ(x-40). WB – 82E1-biotin, biotinylated 82E1 that is directed against the N-terminus of Aβ(1-x) serving as the detection antibody for WB. #, Aβ(1-40) monomers; *, non-specifically detected entities. kDa, kilo-Daltons.

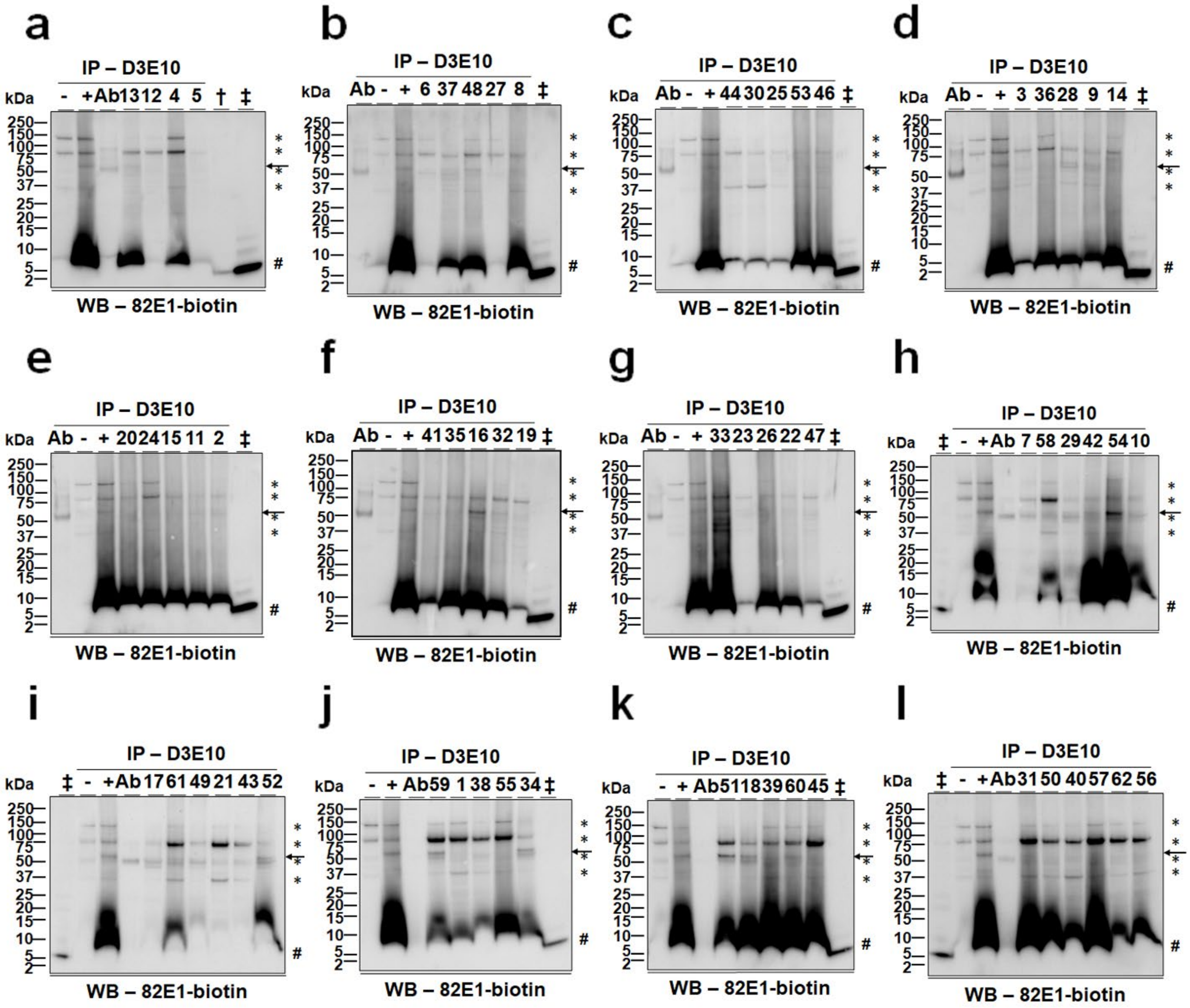
a
b
c
d
e
f
g
h
i
j
k
l
IP – D3E10
kDa
WB – 82E1-biotin
250
150
100
75
50
37
25
20
15
10
5
2
#

**Figure S11. The detection of Aβ(42)*56 in the studied individuals. a-l**) Immunoprecipitation (IP)/western blotting (WB) digital images used for the detection and quantification of Aβ(42)*56 (arrow). IDs of individuals (also see Online Resource **1**: Table S2) are shown above the images. +, IP reactions with the D3E10-immobilized DynaG matrix and aqueous brain extracts of 10-11-month-old APP/TTA mice (serving as a positive control for IP/WB, Online Resource **1**: Table S4). -, IP reactions with the D3E10-immobilized DynaG matrix and aqueous brain extracts of 10-11-month-old littermates of APP/TTA mice that did not express human amyloid precursor proteins (serving as a negative control for IP/WB, Online Resource **1**: Table S4). Ab, IP reactions with the D3E10-immobilized DynaG matrix but no brain extracts. †, synthetic Aβ(1-40) (2 ng), ‡, synthetic Aβ(1-42) (0.5 ng), serving as a positive control for WB. IP – D3E10, IP capture antibody D3E10 that is directed against the C-terminus of Aβ(x-42). WB – 82E1-biotin, biotinylated 82E1 that is directed against the N-terminus of Aβ(1-x) serving as the detection antibody for WB. #, Aβ(1-42) monomers; *, non-specifically detected entities. kDa, kilo-Daltons.

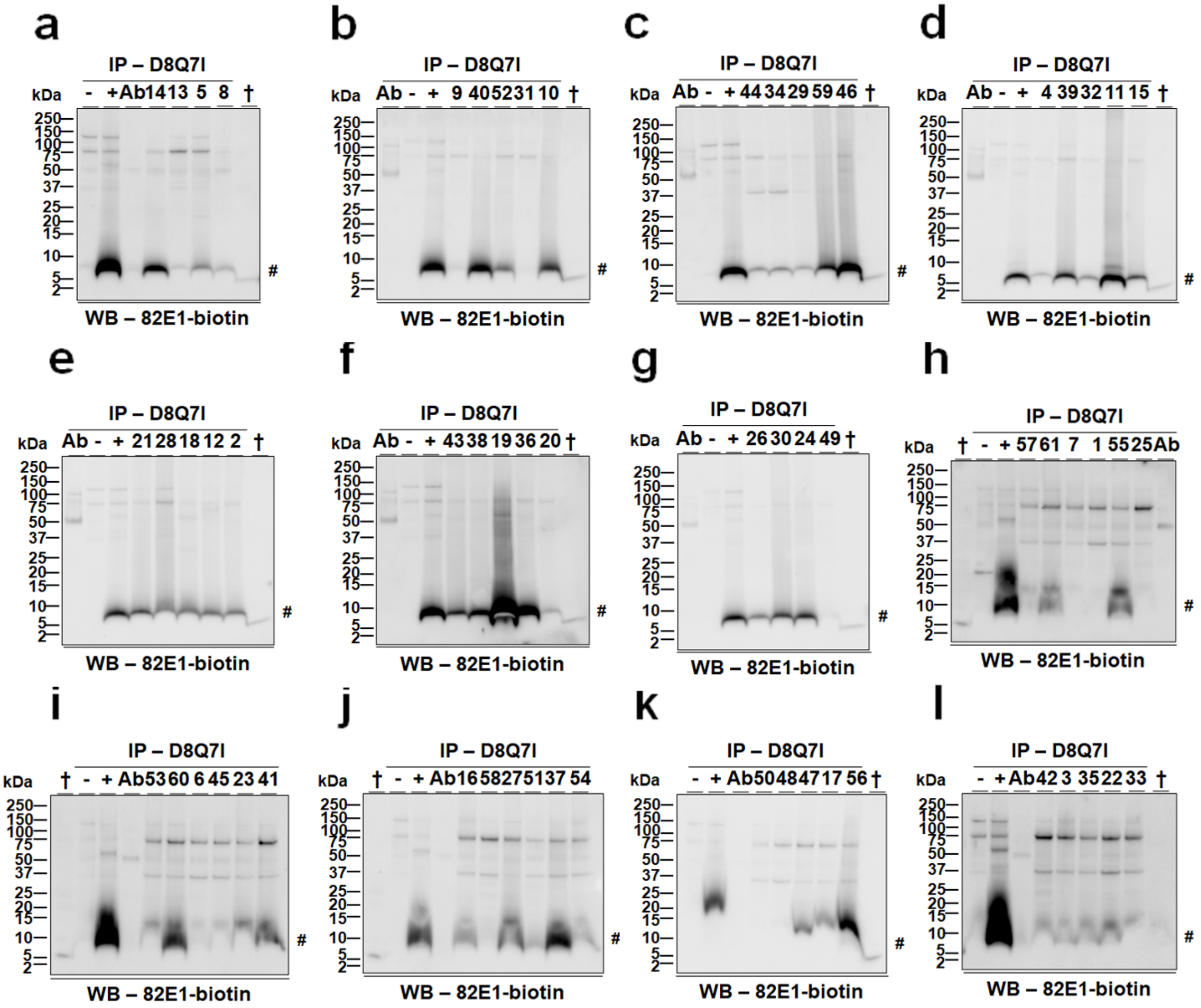
a
IP – D8Q7I
- + Ab 14 13 5 8 †
kDa
250 150 100 75 50 37 25 20 15 10 5 2
#
WB – 82E1-biotin
b
IP – D8Q7I
Ab - + 9 40 52 31 10 †
WB – 82E1-biotin
c
IP – D8Q7I
Ab - + 44 34 29 59 46 †
WB – 82E1-biotin
d
IP – D8Q7I
Ab - + 4 39 32 11 15 †
WB – 82E1-biotin
e
IP – D8Q7I
Ab - + 21 28 18 12 2 †
WB – 82E1-biotin
f
IP – D8Q7I
Ab - + 43 38 19 36 20 †
WB – 82E1-biotin
g
IP – D8Q7I
Ab - + 26 30 24 49 †
WB – 82E1-biotin
h
IP – D8Q7I
† - + 57 61 7 1 55 25 Ab
WB – 82E1-biotin
i
IP – D8Q7I
† - + Ab 53 60 6 45 23 41
WB – 82E1-biotin
j
IP – D8Q7I
† - + Ab 16 58 27 51 37 54
WB – 82E1-biotin
k
IP – D8Q7I
- + Ab 50 48 47 17 56 †
WB – 82E1-biotin
l
IP – D8Q7I
- + Ab 42 3 35 22 33 †
WB – 82E1-biotin

**Figure S12. The detection of Aβ(1-40) monomers in the studied individuals. a-l**) Immunoprecipitation (IP)/western blotting (WB) digital images used for the detection and quantification of Aβ(1-40) monomers (#). IDs of individuals (also see Online Resource **1**: Table S1) are shown above the images. +, IP reactions with the D8Q7I-immobilized DynaG matrix and aqueous brain extracts of 10-11-month-old APP/TTA mice (serving as a positive control for IP/WB, Online Resource **1**: Table S4). -, IP reactions with the D8Q7I-immobilized DynaG matrix and aqueous brain extracts of 10-11-month-old littermates of APP/TTA mice that did not express human amyloid precursor proteins (serving as a negative control for IP/WB, Online Resource **1**: Table S4). Ab, IP reactions with the D8Q7I-immobilized DynaG matrix but no brain extracts. †, synthetic Aβ(1-40) (2 ng), serving as a positive control for WB. IP – D8Q7I, IP capture antibody D8Q7I that is directed against the C-terminus of Aβ(x-40). WB – 82E1-biotin, biotinylated 82E1 that is directed against the N-terminus of Aβ(1-x) serving as the detection antibody for WB. kDa, kilo-Daltons. The WB images shown here are from the same WBs of Figure S10 but at lighter exposure.

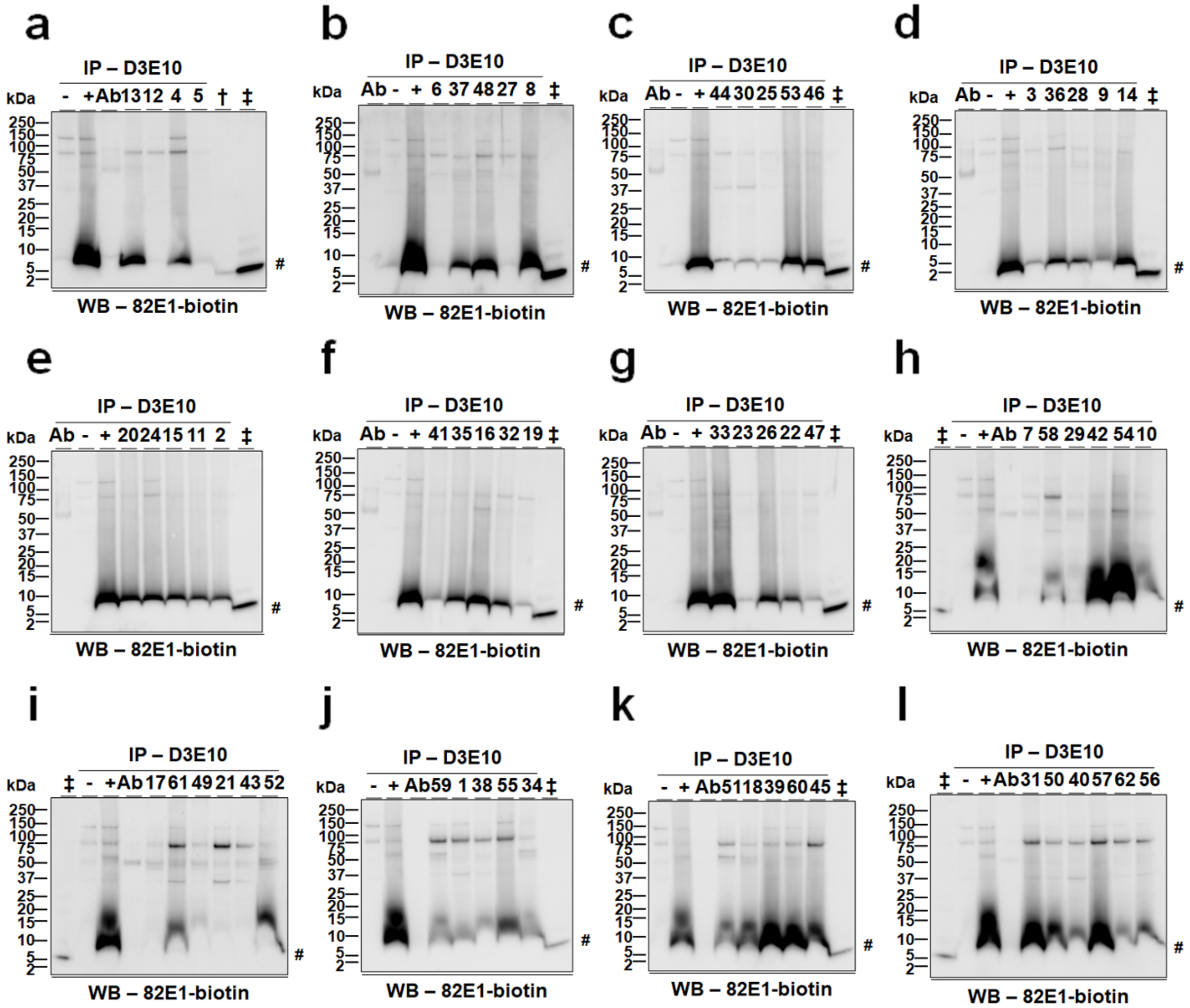
a
b
c
d
e
f
g
h
i
j
k
l
IP – D3E10
kDa
WB – 82E1-biotin
#

**Figure S13. The detection of Aβ(1-42) monomers in the studied individuals. a-l**) Immunoprecipitation (IP)/western blotting (WB) digital images used for the detection and quantification of Aβ(1-42) monomers (#). IDs of individuals (also see Online Resource **1**: Table S2) are shown above the images. +, IP reactions with the D3E10-immobilized DynaG matrix and aqueous brain extracts of 10-11-month-old APP/TTA mice (serving as a positive control for IP/WB, Online Resource **1**: Table S4). -, IP reactions with the D3E10-immobilized DynaG matrix and aqueous brain extracts of 10-11-month-old littermates of APP/TTA mice that did not express human amyloid precursor proteins (serving as a negative control for IP/WB, Online Resource **1**: Table S4). Ab, IP reactions with the D3E10-immobilized DynaG matrix but no brain extracts. †, synthetic Aβ(1-40) (2 ng), ‡, synthetic Aβ(1-42) (0.5 ng), serving as a positive control for WB. IP – D3E10, IP capture antibody D3E10 that is directed against the C-terminus of Aβ(x-42). WB – 82E1-biotin, biotinylated 82E1 that is directed against the N-terminus of Aβ(1-x) serving as the detection antibody for WB. kDa, kilo-Daltons. The WB images shown here are from the same WBs of Figure S11 but at lighter exposure.

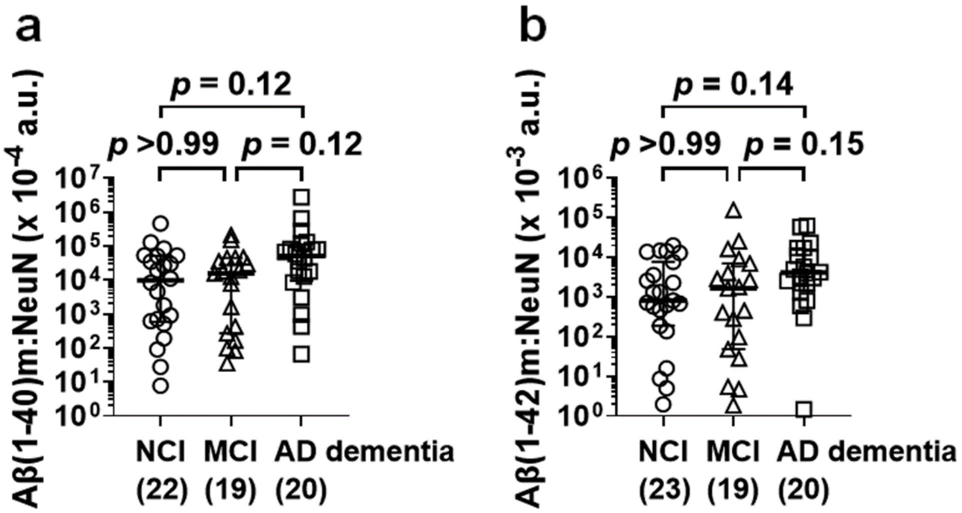
a
b
p = 0.12
p >0.99
p = 0.12
Aβ(1-40)m:NeuN (x 10^-4 a.u.)
NCI (22)
MCI (19)
AD dementia (20)
p = 0.14
p >0.99
p = 0.15
Aβ(1-42)m:NeuN (x 10^-3 a.u.)
NCI (23)
MCI (19)
AD dementia (20)

**Figure S14. Levels of monomeric Aβ are comparable in the inferior temporal gyri of individuals with no cognitive impairment (NCI), mild cognitive impairment (MCI), and Alzheimer's disease (AD) dementia.** (**a, b**) For Aβ(1-40) monomers, following a Kruskal-Wallis test ($H$ (2) = 5.68, $P$ = 0.06), *post hoc* analyses show no difference in NeuN-normalized levels of Aβ(1-40) monomers between individuals with NCI, MCI, and AD dementia (a). In parallel, for Aβ(1-42) monomers, following a Kruskal-Wallis test ($H$ (2) = 5.17, $P$ = 0.08), *post hoc* analyses show no difference in NeuN-normalized levels of Aβ(1-42) monomers between individuals with NCI, MCI, and AD dementia (b). The scatter plots represent data of individuals plus group median ± interquartile range. The y-axes in figures are displayed on log scale. The numbers of individuals with NCI, MCI, and AD dementia used are shown in parentheses. a.u., arbitrary units.

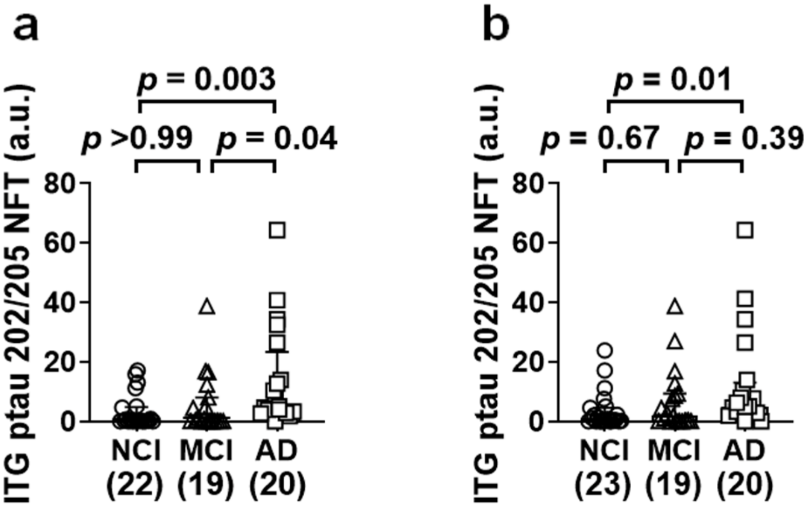
a
p = 0.003
p >0.99
p = 0.04
ITG ptau 202/205 NFT (a.u.)
80
60
40
20
0
NCI (22)
MCI (19)
AD (20)
b
p = 0.01
p = 0.67
p = 0.39
ITG ptau 202/205 NFT (a.u.)
80
60
40
20
0
NCI (23)
MCI (19)
AD (20)

**Figure S15. Comparison of immunohistologically determined neurofibrillary tangle loads using the AT8 antibody recognizing phosphorylated tau at serine 202 and threonine 205 in the inferior temporal gyri of individuals with no cognitive impairment (NCI), mild cognitive impairment (MCI), and Alzheimer's disease (AD) dementia.** Following Kruskal-Wallis tests ("Tangle at ITG" of Tables 3 and 4), *post hoc* analyses show no difference in phosphorylated tau at serine 202 and threonine 205 (ptau 202/205)-containing neurofibrillary tangle loads in the inferior temporal gyrus (ITG ptau 202/205 NFT) between the individuals with NCI and those with MCI. However, for the cohort used to study Aβ(40)*56 (**a**), the ITG ptau 202/205 NFT of the individuals with AD dementia are higher than those of the individuals with NCI as well as MCI; for the cohort used to study Aβ(42)*56 (**b**), the ITG ptau 202/205 NFT of the individuals with AD dementia are higher than those of the individuals with NCI. The scatter plots represent data of individuals plus group median ± interquartile range. The numbers of individuals with NCI, MCI, and AD dementia used are shown in parentheses. a.u., arbitrary units.

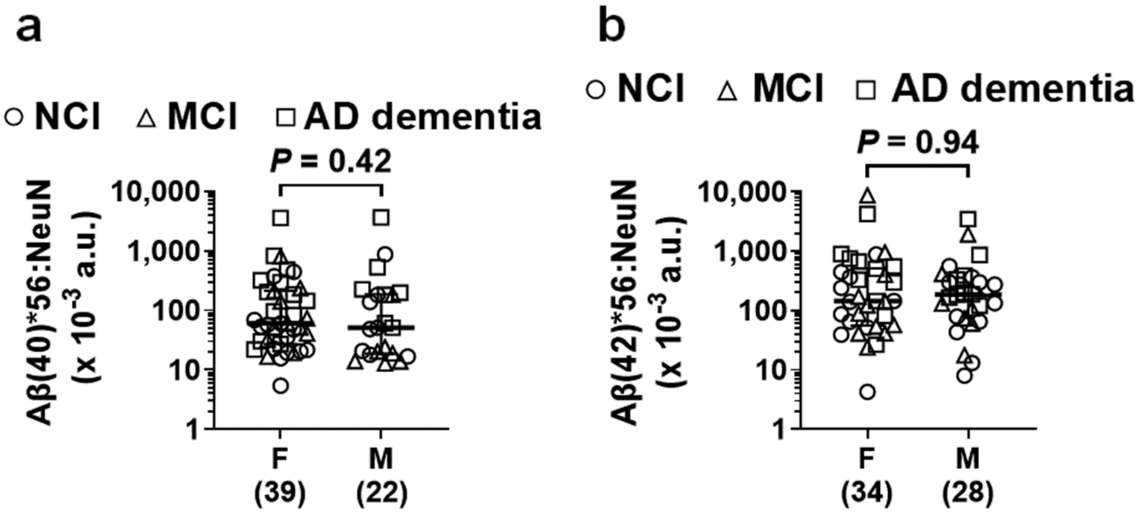
a
○ NCI △ MCI □ AD dementia
P = 0.42
Aβ(40)*56:NeuN
(x 10^-3 a.u.)
10,000
1,000
100
10
1
F
(39)
M
(22)
b
○ NCI △ MCI □ AD dementia
P = 0.94
Aβ(42)*56:NeuN
(x 10^-3 a.u.)
10,000
1,000
100
10
1
F
(34)
M
(28)

**Figure S16. Levels of Aβ*56 variants, normalized to levels of neuronal marker NeuN, are comparable between male and female individuals.** (**a**) For Aβ*(40)56, $U = 374$, $P = 0.42$, two-tailed Mann-Whitney *U* test. (**b**) For Aβ(42)*56, $U = 470$, $P = 0.94$, two-tailed Mann-Whitney *U* test. The scatter plots represent data of individuals plus group median ± interquartile range. The y-axes in figures are displayed on log scale. The numbers of male and female individuals used for the analyses are shown in parentheses. NCI, no cognitive impairment; MCI, mild cognitive impairment; AD dementia, Alzheimer's disease dementia. F, female; M, male. a.u., arbitrary units.

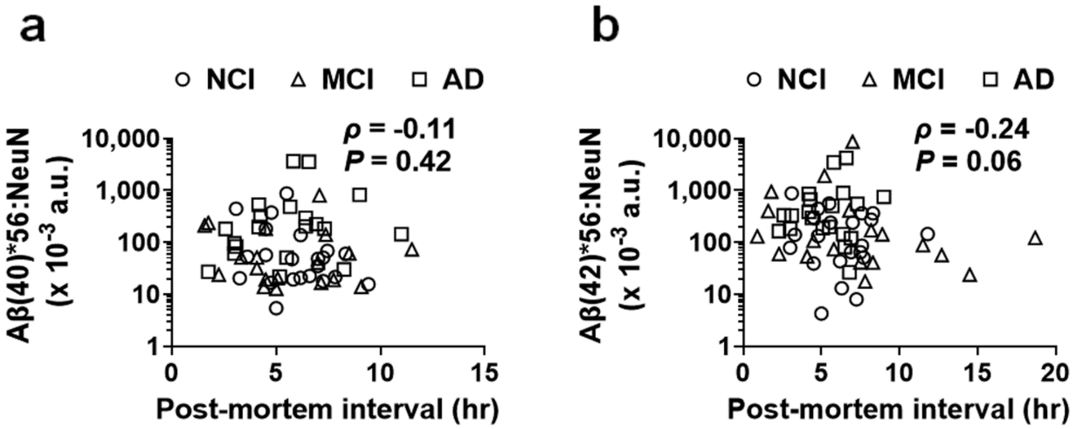
a
NCI MCI AD
ρ = -0.11
P = 0.42
Aβ(40)*56:NeuN (x 10^-3 a.u.)
10,000
1,000
100
10
1
0 5 10 15
Post-mortem interval (hr)
b
NCI MCI AD
ρ = -0.24
P = 0.06
Aβ(42)*56:NeuN (x 10^-3 a.u.)
10,000
1,000
100
10
1
0 5 10 15 20
Post-mortem interval (hr)

**Figure S17. Levels of Aβ(40)*56 (a) and Aβ(42)*56 (b), normalized to levels of neuronal marker NeuN, do not correlate to time intervals from death of individuals to autopsy.** Spearman's rank-order correlations were used for statistical analyses. The scatter plots represent data of individuals. The y-axes in figures are displayed on log scale. NCI, no cognitive impairment; MCI, mild cognitive impairment; AD dementia, Alzheimer's disease dementia. hr, hours. a.u., arbitrary units.

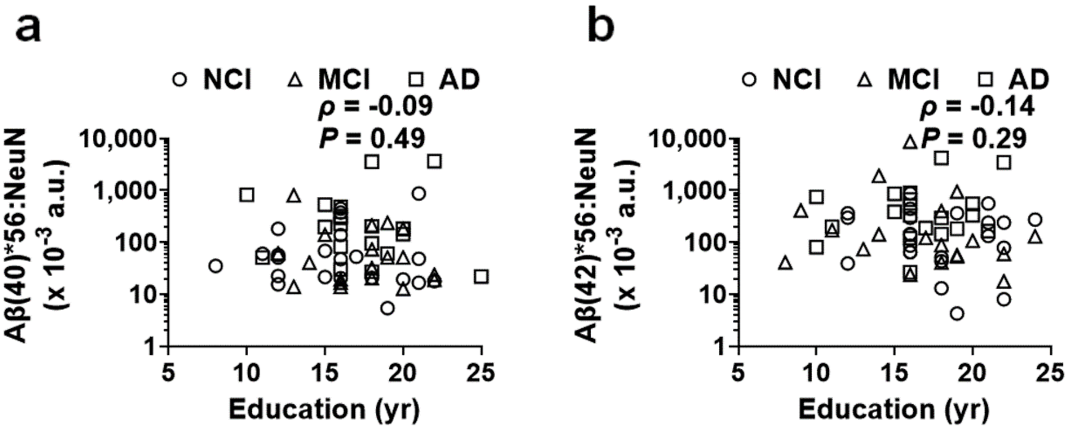
a
NCI MCI AD
ρ = -0.09
P = 0.49
Aβ(40)*56:NeuN (x 10-3 a.u.)
10,000
1,000
100
10
1
5 10 15 20 25
Education (yr)
b
NCI MCI AD
ρ = -0.14
P = 0.29
Aβ(42)*56:NeuN (x 10-3 a.u.)
10,000
1,000
100
10
1
5 10 15 20 25
Education (yr)

**Figure S18. Levels of Aβ(40)*56 (a) and Aβ(42)*56 (b), normalized to levels of neuronal marker NeuN, do not correlate to the education-receiving time of individuals.** Spearman's rank-order correlations were used for statistical analyses. The scatter plots represent data of individuals. The y-axes in figures are displayed on log scale. NCI, no cognitive impairment; MCI, mild cognitive impairment; AD dementia, Alzheimer's disease dementia. yr, years. a.u., arbitrary units.

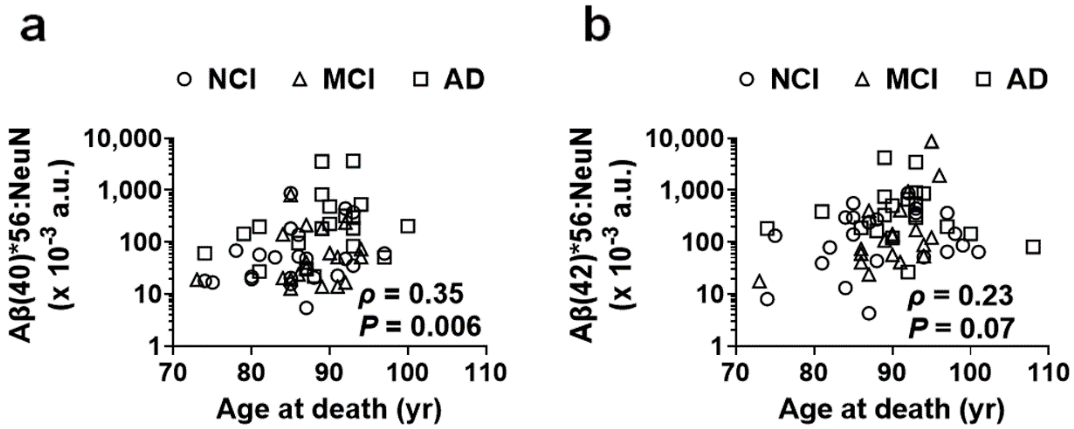
a
b
NCI MCI AD
Aβ(40)*56:NeuN (x 10^-3 a.u.)
Aβ(42)*56:NeuN (x 10^-3 a.u.)
ρ = 0.35
P = 0.006
ρ = 0.23
P = 0.07
Age at death (yr)

**Figure S19. Levels of Aβ(40)*56 (a) but not Aβ(42)*56 (b), normalized to levels of neuronal marker NeuN, correlate to the ages of individuals at death.** Spearman's rank-order correlations were used for statistical analyses. The scatter plots represent data of individuals. The y-axes in figures are displayed on log scale. NCI, no cognitive impairment; MCI, mild cognitive impairment; AD dementia, Alzheimer's disease dementia. yr, years. a.u., arbitrary units.

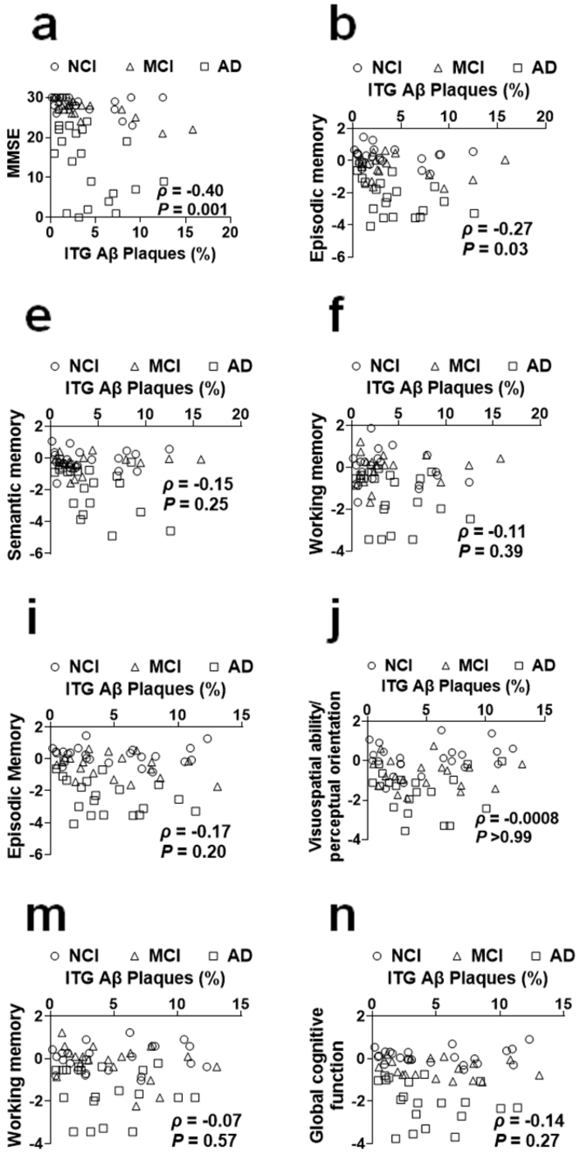
a
NCI MCI AD
MMSE
ITG Aβ Plaques (%)
ρ = -0.40
P = 0.001
b
Episodic memory
ρ = -0.27
P = 0.03
c
Visuospatial ability/ perceptual orientation
ρ = -0.08
P = 0.53
d
Perceptual speed
ρ = -0.17
P = 0.18
e
Semantic memory
ρ = -0.15
P = 0.25
f
Working memory
ρ = -0.11
P = 0.39
g
Global cognitive function
ρ = -0.23
P = 0.08
h
MMSE
ρ = -0.17
P = 0.20
i
Episodic Memory
ρ = -0.17
P = 0.20
j
Visuospatial ability/ perceptual orientation
ρ = -0.0008
P >0.99
k
Perceptual speed
ρ = -0.05
P = 0.69
l
Semantic memory
ρ = 0.03
P = 0.81
m
Working memory
ρ = -0.07
P = 0.57
n
Global cognitive function
ρ = -0.14
P = 0.27

**Figure S20. The correlation of immunohistologically determined β-amyloid plaque loads in the inferior temporal gyrus (ITG Aβ Plaques) and degrees of cognitive and memory impairment.** (**a, h**) Correlations of ITG Aβ plaque loads and the mini-mental state examination (MMSE) scores for the Aβ(40)*56 cohort (a) and the Aβ(42)*56 cohort (h). (**b, i**) Correlations of ITG Aβ plaque loads and the z scores of composite measures of episodic memory for the Aβ(40)*56 cohort (b) and the Aβ(42)*56 cohort (i). (**c, j**) Correlations of ITG Aβ plaque loads and the z scores of composite measures of visuospatial ability/perceptual orientation for the Aβ(40)*56 cohort (c) and the Aβ(42)*56 cohort (j). (**d, k**) Correlations of ITG Aβ plaque loads and the z scores of composite measures of perceptual speed for the Aβ(40)*56 cohort (d) and the Aβ(42)*56 cohort (k). (**e, l**) Correlations of ITG Aβ plaque loads and the z scores of composite measures of semantic memory for the Aβ(40)*56 cohort (e) and the Aβ(42)*56 cohort (l). (**f, m**) Correlations of ITG Aβ plaque loads and the z scores of composite measures of working memory for the Aβ(40)*56 cohort (f) and the Aβ(42)*56 cohort (m). (**g, n**) Correlations of ITG Aβ plaque loads and the z scores of composite measures of global cognitive function for the Aβ(40)*56 cohort (g) and the Aβ(42)*56 cohort (n). For Figures a-n, Spearman's rank-order correlations were used, and the scatter plots represent data of individuals. NCI, no cognitive impairment; MCI, mild cognitive impairment; AD dementia, Alzheimer's disease dementia.

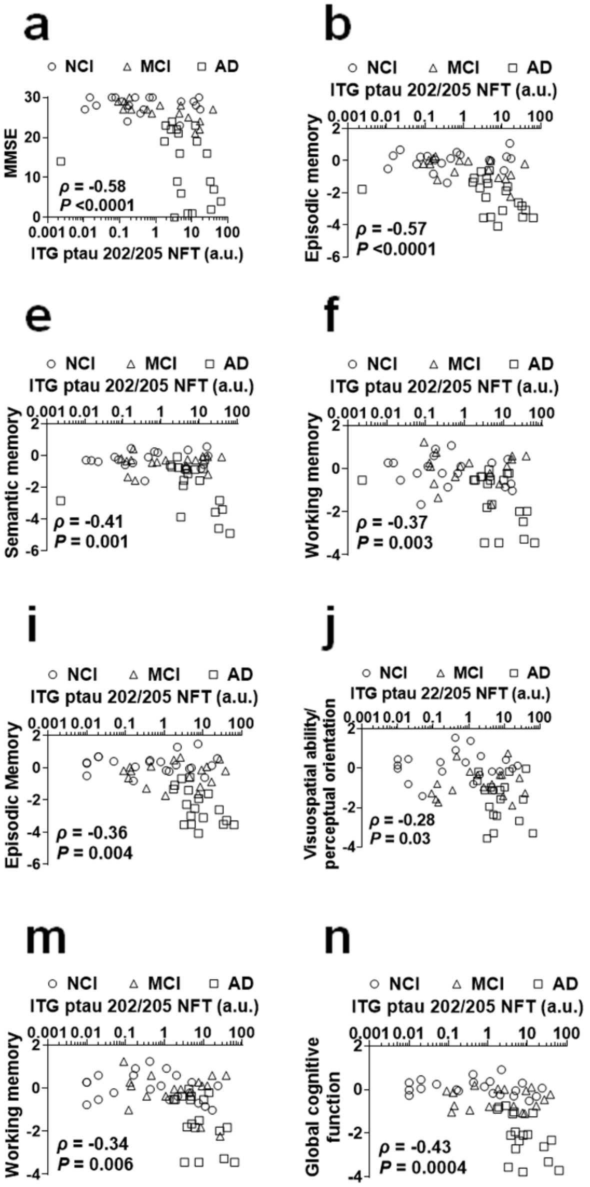
a
NCI MCI AD
MMSE
ITG ptau 202/205 NFT (a.u.)
ρ = -0.58
P <0.0001
b
Episodic memory
ρ = -0.57
P <0.0001
c
Visuospatial ability/ perceptual orientation
ρ = -0.30
P = 0.02
d
Perceptual speed
ρ = -0.47
P = 0.0001
e
Semantic memory
ρ = -0.41
P = 0.001
f
Working memory
ρ = -0.37
P = 0.003
g
Global cognitive function
ρ = -0.57
P <0.0001
h
MMSE
ρ = -0.39
P = 0.002
i
Episodic Memory
ρ = -0.36
P = 0.004
j
ITG ptau 22/205 NFT (a.u.)
Visuospatial ability/ perceptual orientation
ρ = -0.28
P = 0.03
k
Perceptual speed
ρ = -0.43
P = 0.0004
l
Semantic memory
ρ = -0.29
P = 0.02
m
Working memory
ρ = -0.34
P = 0.006
n
Global cognitive function
ρ = -0.43
P = 0.0004

**Figure S21. The correlation of immunohistologically determined neurofibrillary tangle loads using the AT8 antibody recognizing phosphorylated tau at serine 202 and threonine 205 in the inferior temporal gyrus (ITG ptau 202/205 NFT) and degrees of cognitive and memory impairment.** (**a, h**) Correlations of ITG ptau 202/205 NFT and the mini-mental state examination (MMSE) scores for the Aβ(40)*56 cohort (a) and the Aβ(42)*56 cohort (h). (**b, i**) Correlations of ITG ptau 202/205 NFT and the z scores of composite measures of episodic memory for the Aβ(40)*56 cohort (b) and the Aβ(42)*56 cohort (i). (**c, j**) Correlations of ITG ptau 202/205 NFT and the z scores of composite measures of visuospatial ability/perceptual orientation for the Aβ(40)*56 cohort (c) and the Aβ(42)*56 cohort (j). (**d, k**) Correlations of ITG ptau 202/205 NFT and the z scores of composite measures of perceptual speed for the Aβ(40)*56 cohort (d) and the Aβ(42)*56 cohort (k). (**e, l**) Correlations of ITG ptau 202/205 NFT and the z scores of composite measures of semantic memory for the Aβ(40)*56 cohort (e) and the Aβ(42)*56 cohort (l). (**f, m**) Correlations of ITG ptau 202/205 NFT and the z scores of composite measures of working memory for the Aβ(40)*56 cohort (f) and the Aβ(42)*56 cohort (m). (**g, n**) Correlations of ITG ptau 202/205 NFT and the z scores of composite measures of global cognitive function for the Aβ(40)*56 cohort (g) and the Aβ(42)*56 cohort (n). For Figures a-n, Spearman's rank-order correlations were used, the scatter plots represent data of individuals, and the x-axes in figures are displayed on log scale. a.u., arbitrary units. NCI, no cognitive impairment; MCI, mild cognitive impairment; AD dementia, Alzheimer's disease dementia.

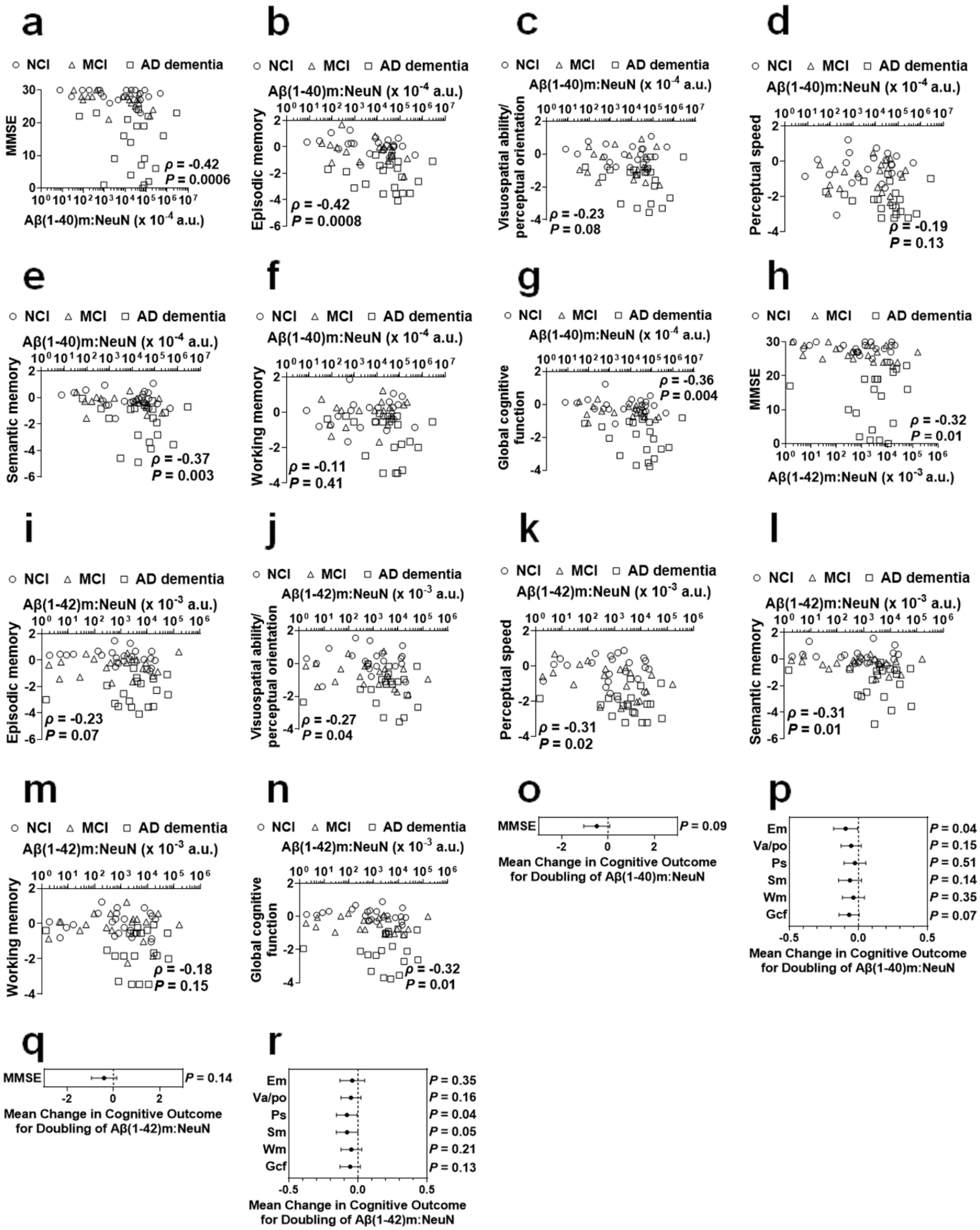
a
○ NCI △ MCI □ AD dementia
MMSE
ρ = -0.42
P = 0.0006
Aβ(1-40)m:NeuN (x 10^-4 a.u.)
b
○ NCI △ MCI □ AD dementia
Aβ(1-40)m:NeuN (x 10^-4 a.u.)
Episodic memory
ρ = -0.42
P = 0.0008
c
○ NCI △ MCI □ AD dementia
Aβ(1-40)m:NeuN (x 10^-4 a.u.)
Visuospatial ability/ perceptual orientation
ρ = -0.23
P = 0.08
d
○ NCI △ MCI □ AD dementia
Aβ(1-40)m:NeuN (x 10^-4 a.u.)
Perceptual speed
ρ = -0.19
P = 0.13
e
○ NCI △ MCI □ AD dementia
Aβ(1-40)m:NeuN (x 10^-4 a.u.)
Semantic memory
ρ = -0.37
P = 0.003
f
○ NCI △ MCI □ AD dementia
Aβ(1-40)m:NeuN (x 10^-4 a.u.)
Working memory
ρ = -0.11
P = 0.41
g
○ NCI △ MCI □ AD dementia
Aβ(1-40)m:NeuN (x 10^-4 a.u.)
Global cognitive function
ρ = -0.36
P = 0.004
h
○ NCI △ MCI □ AD dementia
MMSE
ρ = -0.32
P = 0.01
Aβ(1-42)m:NeuN (x 10^-3 a.u.)
i
○ NCI △ MCI □ AD dementia
Aβ(1-42)m:NeuN (x 10^-3 a.u.)
Episodic memory
ρ = -0.23
P = 0.07
j
○ NCI △ MCI □ AD dementia
Aβ(1-42)m:NeuN (x 10^-3 a.u.)
Visuospatial ability/ perceptual orientation
ρ = -0.27
P = 0.04
k
○ NCI △ MCI □ AD dementia
Aβ(1-42)m:NeuN (x 10^-3 a.u.)
Perceptual speed
ρ = -0.31
P = 0.02
l
○ NCI △ MCI □ AD dementia
Aβ(1-42)m:NeuN (x 10^-3 a.u.)
Semantic memory
ρ = -0.31
P = 0.01
m
○ NCI △ MCI □ AD dementia
Aβ(1-42)m:NeuN (x 10^-3 a.u.)
Working memory
ρ = -0.18
P = 0.15
n
○ NCI △ MCI □ AD dementia
Aβ(1-42)m:NeuN (x 10^-3 a.u.)
Global cognitive function
ρ = -0.32
P = 0.01
o
MMSE
P = 0.09
Mean Change in Cognitive Outcome for Doubling of Aβ(1-40)m:NeuN
p
Em P = 0.04
Va/po P = 0.15
Ps P = 0.51
Sm P = 0.14
Wm P = 0.35
Gcf P = 0.07
Mean Change in Cognitive Outcome for Doubling of Aβ(1-40)m:NeuN
q
MMSE
P = 0.14
Mean Change in Cognitive Outcome for Doubling of Aβ(1-42)m:NeuN
r
Em P = 0.35
Va/po P = 0.16
Ps P = 0.04
Sm P = 0.05
Wm P = 0.21
Gcf P = 0.13
Mean Change in Cognitive Outcome for Doubling of Aβ(1-42)m:NeuN

**Figure S22. The correlation of levels of monomeric Aβ and degrees of cognitive and memory impairment.** (**a, h**) Inverse correlations of levels of monomeric Aβ(1-40) (a) and monomeric Aβ(1-42) (h) (normalized to levels of NeuN) to the mini-mental state examination (MMSE) scores. (**b, i**) Correlations of NeuN-normalized levels of monomeric Aβ(1-40) (b) and monomeric Aβ(1-42) (i) to z scores of composite measures of episodic memory. (**c, j**) Correlations of NeuN-normalized levels of monomeric Aβ(1-40) (c) and monomeric Aβ(1-42) (j) to z scores of composite measures of visuospatial ability/perceptual orientation. (**d, k**) Correlations of NeuN-normalized levels of monomeric Aβ(1-40) (d) and monomeric Aβ(1-42) (k) to z scores of composite measures of perceptual speed. (**e, l**) Correlations of NeuN-normalized levels of monomeric Aβ(1-40) (e) and monomeric Aβ(1-42) (l) to z scores of composite measures of semantic memory. (**f, m**) Correlations of NeuN-normalized levels of monomeric Aβ(1-40) (f) and monomeric Aβ(1-42) (m) to z scores of composite measures of working memory. (**g, n**) Correlations of NeuN-normalized levels of monomeric Aβ(1-40) (g) and monomeric Aβ(1-42) (n) to z scores of composite measures of global cognitive function. (**o, q**) Following adjustment for age at death, sex, years of education, and log-transformed Aβ plaque loads in the inferior temporal gyrus (ITG) using multiple linear regression (MLR), NeuN-normalized levels of monomeric Aβ(1-40) (o) and monomeric Aβ(1-42) (q) do not correlate to MMSE scores. (**p, r**) Following adjustment for age at death, sex, years of education, and log-transformed Aβ plaque loads in the ITG using MLR, NeuN-normalized levels of monomeric Aβ(1-40) (q) and monomeric Aβ(1-42) (r) do not correlate to degrees of cognitive or memory impairment except for weak correlations between NeuN-normalized levels of monomeric Aβ(1-40) and z scores of composite measures of episodic memory and between NeuN-normalized levels of monomeric Aβ(1-42) and z scores of composite measures of perceptual speed. For Figures a-n, Spearman's rank-order correlations were used, the scatter plots represent data of individuals, and the x-axes are displayed on log scale. a.u., arbitrary units. NCI, no cognitive impairment; MCI, mild cognitive impairment; AD dementia, Alzheimer's disease dementia. Em, episodic memory; Va/po, visuospatial ability/perceptual orientation; Ps, perceptual speed; Sm, semantic memory; Wm, working memory; Gcf, global cognitive function.

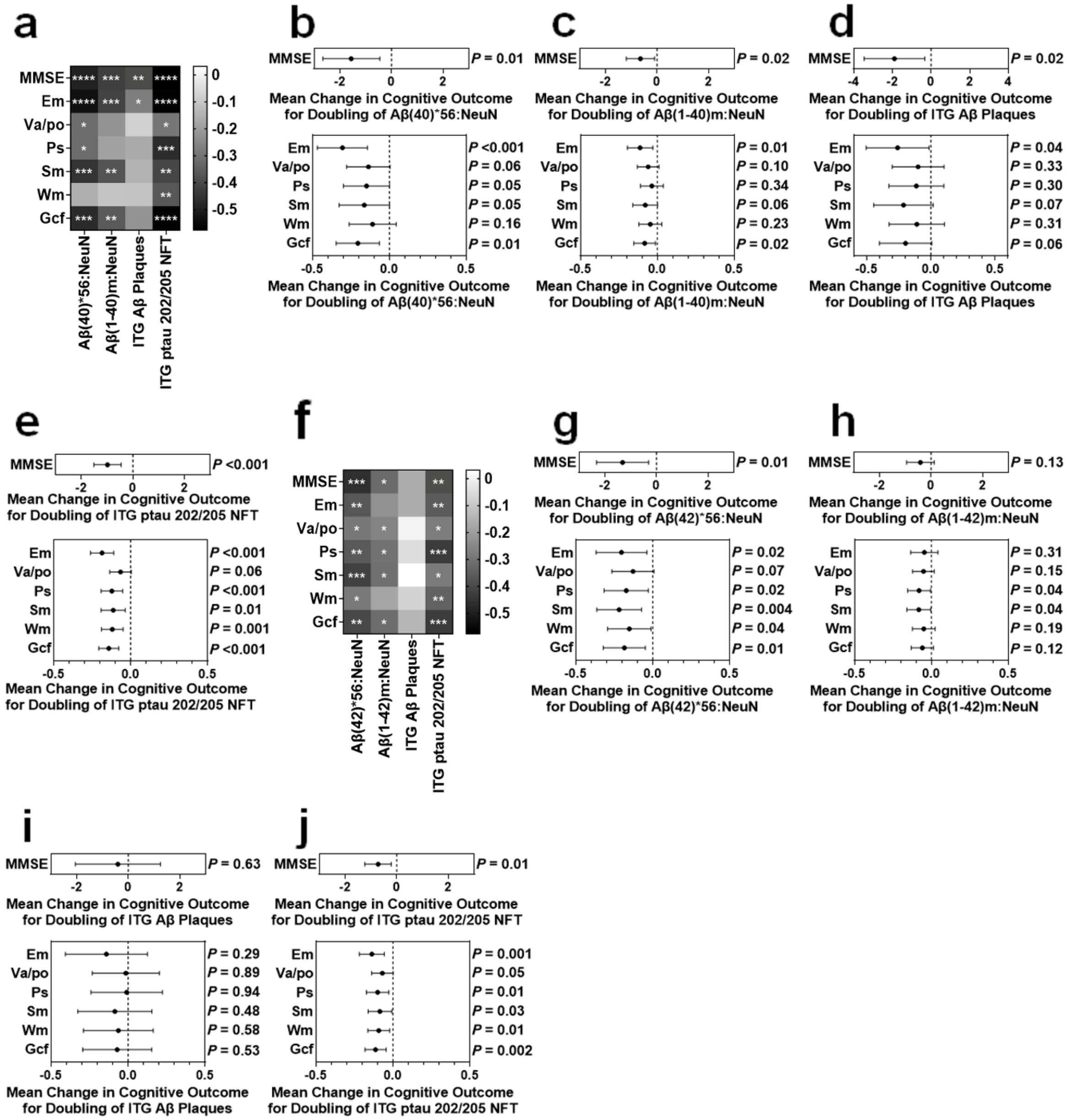
a
MMSE
Em
Va/po
Ps
Sm
Wm
Gcf
Aβ(40)*56:NeuN
Aβ(1-40)m:NeuN
ITG Aβ Plaques
ITG ptau 202/205 NFT
b
MMSE P = 0.01
Mean Change in Cognitive Outcome for Doubling of Aβ(40)*56:NeuN
Em P <0.001
Va/po P = 0.06
Ps P = 0.05
Sm P = 0.05
Wm P = 0.16
Gcf P = 0.01
Mean Change in Cognitive Outcome for Doubling of Aβ(40)*56:NeuN
c
MMSE P = 0.02
Mean Change in Cognitive Outcome for Doubling of Aβ(1-40)m:NeuN
Em P = 0.01
Va/po P = 0.10
Ps P = 0.34
Sm P = 0.06
Wm P = 0.23
Gcf P = 0.02
Mean Change in Cognitive Outcome for Doubling of Aβ(1-40)m:NeuN
d
MMSE P = 0.02
Mean Change in Cognitive Outcome for Doubling of ITG Aβ Plaques
Em P = 0.04
Va/po P = 0.33
Ps P = 0.30
Sm P = 0.07
Wm P = 0.31
Gcf P = 0.06
Mean Change in Cognitive Outcome for Doubling of ITG Aβ Plaques
e
MMSE P <0.001
Mean Change in Cognitive Outcome for Doubling of ITG ptau 202/205 NFT
Em P <0.001
Va/po P = 0.06
Ps P <0.001
Sm P = 0.01
Wm P = 0.001
Gcf P <0.001
Mean Change in Cognitive Outcome for Doubling of ITG ptau 202/205 NFT
f
MMSE
Em
Va/po
Ps
Sm
Wm
Gcf
Aβ(42)*56:NeuN
Aβ(1-42)m:NeuN
ITG Aβ Plaques
ITG ptau 202/205 NFT
g
MMSE P = 0.01
Mean Change in Cognitive Outcome for Doubling of Aβ(42)*56:NeuN
Em P = 0.02
Va/po P = 0.07
Ps P = 0.02
Sm P = 0.004
Wm P = 0.04
Gcf P = 0.01
Mean Change in Cognitive Outcome for Doubling of Aβ(42)*56:NeuN
h
MMSE P = 0.13
Mean Change in Cognitive Outcome for Doubling of Aβ(1-42)m:NeuN
Em P = 0.31
Va/po P = 0.15
Ps P = 0.04
Sm P = 0.04
Wm P = 0.19
Gcf P = 0.12
Mean Change in Cognitive Outcome for Doubling of Aβ(1-42)m:NeuN
i
MMSE P = 0.63
Mean Change in Cognitive Outcome for Doubling of ITG Aβ Plaques
Em P = 0.29
Va/po P = 0.89
Ps P = 0.94
Sm P = 0.48
Wm P = 0.58
Gcf P = 0.53
Mean Change in Cognitive Outcome for Doubling of ITG Aβ Plaques
j
MMSE P = 0.01
Mean Change in Cognitive Outcome for Doubling of ITG ptau 202/205 NFT
Em P = 0.001
Va/po P = 0.05
Ps P = 0.01
Sm P = 0.03
Wm P = 0.01
Gcf P = 0.002
Mean Change in Cognitive Outcome for Doubling of ITG ptau 202/205 NFT

**Figure S23. The associations between four neuropathological and biochemical measures and seven types of neuropsychological scores.** (**a**) A heat map illustrating in the inferior temporal gyrus (ITG) the correlations of NeuN-normalized levels of Aβ(40)*56 (Aβ(40)*56:NeuN), NeuN-normalized levels of monomeric Aβ(1-40) (Aβ(1-40)m:NeuN), Aβ plaque loads (ITG Aβ Plaques), and phosphorylated tau at serine 202 and threonine 205 (ptau 202/205)-containing neurofibrillary tangle loads (ITG ptau 202/205 NFT) to z scores of composite measures of seven neuropsychological tests: mini-mental state examination (MMSE), episodic memory (Em), visuospatial ability/perceptual orientation (Va/po), perceptual speed (Ps), semantic memory (Sm), working memory (Wm), and global cognitive function (Gcf). The data shown in the columns Aβ(40)*56:NeuN, Aβ(1-40)m:NeuN, ITG Aβ Plaques, and ITG ptau 202/205 NFT are summarized from Figures 4a-g, Figures S22a-g, Figures S20a-g, and Figures S21a-g, respectively. (**b**) Following adjustment for age at death, sex, and years of education using multiple linear regression (MLR), Aβ(40)*56:NeuN in the ITG remain inversely correlated to MMSE scores (upper panels) and remain correlated to z scores of composite measures of episodic memory and global cognitive function (lower panels). (**c**) Following adjustment for age at death, sex, and years of education using MLR, Aβ(1-40)m:NeuN in the ITG remain inversely correlated to MMSE scores (upper panels) and remain correlated to z scores of composite measures of episodic memory and global cognitive function (lower panels). (**d**) Following adjustment for age at death, sex, and years of education using MLR, ITG Aβ Plaques of the cohort for measuring Aβ(40)*56 remain inversely correlated to MMSE scores (upper panels) and remain correlated to z scores of composite measures of episodic memory (lower panels). (**e**) Following adjustment for age at death, sex, and years of education using MLR, ITG ptau 202/205 NFT of the cohort for measuring Aβ(40)*56 remain inversely correlated to MMSE scores (upper panels) and remain correlated to z scores of composite measures of episodic memory, perceptual speed, semantic memory, working memory, and global cognitive function (lower panels). (**f**) A heat map illustrating in the ITG the correlations of NeuN-normalized levels of Aβ(42)*56 (Aβ(42)*56:NeuN), NeuN-normalized levels of monomeric Aβ(1-42) (Aβ(1-42)m:NeuN), ITG Aβ Plaques, and ITG ptau 202/205 NFT to z scores of composite measures of the seven neuropsychological tests. The data shown in the columns Aβ(42)*56:NeuN, Aβ(1-42)m:NeuN, ITG Aβ Plaques, and ITG ptau 202/205 NFT are summarized from Figures 4h-n, Figures S22h-n, Figures S20h-n, and Figures S21h-n, respectively. (**g**) Following adjustment for age at death, sex, and years of education using MLR, Aβ(42)*56:NeuN in the ITG remain inversely correlated to MMSE scores (upper panels) and remain correlated to z scores of composite measures of episodic memory, perceptual speed, semantic memory, working memory, and global cognitive function (lower panels). (**h**) Following adjustment for age at death, sex, and years of education using MLR, Aβ(1-42)m:NeuN in the ITG are not correlated to MMSE scores (upper panels) but remain correlated to z scores of composite measures of perceptual speed and semantic memory (lower panels). (**i**) Following adjustment for age at death, sex, and years of education using MLR, ITG Aβ Plaques of the cohort for measuring Aβ(42)*56 are not correlated to MMSE scores (upper panels) or other neuropsychological test scores (lower panels). (**j**) Following adjustment for age at death, sex, and years of education using MLR, ITG ptau 202/205 NFT of the cohort for measuring Aβ(42)*56 remain inversely correlated to MMSE scores (upper panels) and remain correlated to z scores of composite measures of episodic memory, perceptual speed, semantic memory, working memory, and global cognitive function (Lower panels). For Figures a and f, the gray scales represent Spearman's rank-order correlation coefficients. **** $P$ <0.0001; *** $P$ <0.001, ** $P$ <0.01, * $P$ <0.05. For Figures b-e and g-j, mean changes in cognitive outcome for doubling of the four neuropathological and biochemical measures and corresponding *P* values are shown.

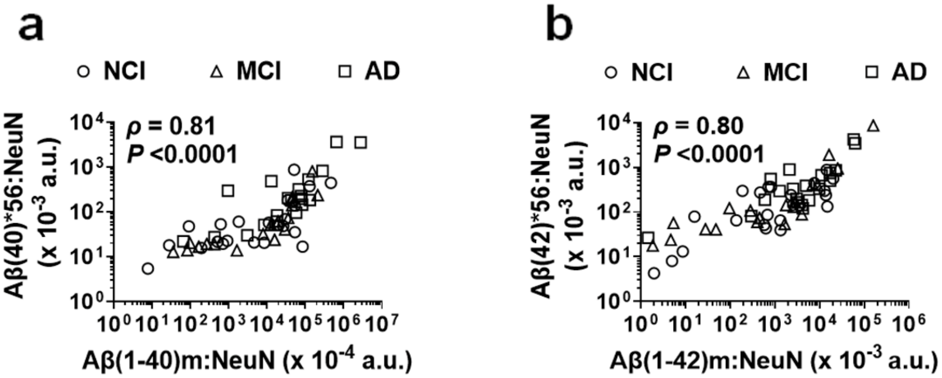
a
NCI
MCI
AD
ρ = 0.81
P <0.0001
Aβ(40)*56:NeuN (x 10-3 a.u.)
Aβ(1-40)m:NeuN (x 10-4 a.u.)
b
NCI
MCI
AD
ρ = 0.80
P <0.0001
Aβ(42)*56:NeuN (x 10-3 a.u.)
Aβ(1-42)m:NeuN (x 10-3 a.u.)

**Figure S24. Levels of Aβ(40)*56 (a) and Aβ(42)*56 (b), normalized to levels of neuronal marker NeuN, correlate to the levels of correspondingly normalized monomeric canonical Aβ.** Spearman's rank-order correlations were used for statistical analyses. The scatter plots represent data of individuals. The x- and y-axes in figures are displayed on log scale. NCI, no cognitive impairment; MCI, mild cognitive impairment; AD dementia, Alzheimer's disease dementia. Aβ(1-40)m, monomeric Aβ(1-40); Aβ(1-42)m, monomeric Aβ(1-42). a.u., arbitrary units.

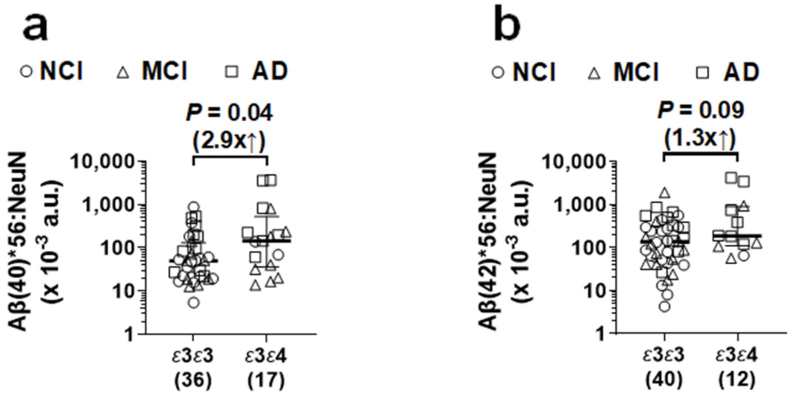
a
NCI
MCI
AD
P = 0.04
(2.9x↑)
Aβ(40)*56:NeuN
(x 10-3 a.u.)
10,000
1,000
100
10
1
ε3ε3
(36)
ε3ε4
(17)
b
NCI
MCI
AD
P = 0.09
(1.3x↑)
Aβ(42)*56:NeuN
(x 10-3 a.u.)
10,000
1,000
100
10
1
ε3ε3
(40)
ε3ε4
(12)

**Figure S25. Levels of Aβ*56 variants, normalized to levels of neuronal marker NeuN, are higher in individuals with apolipoprotein E (*APOE*) *ε3ε4* alleles than those with *APOE ε3ε3* alleles.** Levels of normalized Aβ(40)*56 (**a**) and Aβ(42)*56 (**b**) in *APOE ε3ε4* individuals are 2.9- and 1.3- fold higher than *APOE ε3ε3* individuals, respectively. Two-tailed Mann-Whitney *U* tests were used for the analyses. For Aβ(40)*56, $U = 200$, $P = 0.04$; for Aβ(42)*56, $U = 161$, $P = 0.09$. The scatter plots represent data of individuals plus group median ± interquartile range. The y-axes in figures are displayed on log scale. The numbers of individuals used for the analyses are shown in parentheses. NCI, no cognitive impairment; MCI, mild cognitive impairment; AD dementia, Alzheimer's disease dementia. a.u., arbitrary units.

**Table S1 Detailed demographic and neuropathological characteristics of individuals for measuring Aβ(40)*56**

| ID | Diagnosis[a] | Age[b] (yr[c]) | Sex[d] | PMI[e] (hr[f]) | Education (yr[c]) | *APOE*[g] genotype | Br. Wt. (g)[h] | Amyloid-ITG[i] | Tangle-ITG[j] | CERAD[k] | BST[l] | NIA-Reagan[m] |
|---|---|---|---|---|---|---|---|---|---|---|---|---|
| 1 | MCI | 91 | M | 9.1 | 13 | *ε3ε3* | 1080 | 1.3 | 0.2 | 1 | 1 | 1 |
| 2 | AD dementia | 94 | M | 4.2 | 15 | *ε3ε3* | 1170 | 3.5 | 3.8 | 3 | 5 | 3 |
| 3 | NCI | 83 | M | 7.3 | 12 | *ε3ε3* | 1425 | 1.8 | 0.1 | 2 | 0 | 1 |
| 4 | NCI | 88 | M | 6.2 | 18 | *ε3ε3* | 1170 | 0.9 | 0.01 | 0 | 1 | 1 |
| 5 | NCI | 87 | M | 7.0 | 21 | *ε3ε3* | 1160 | 7.1 | 17.1 | 2 | 5 | 2 |
| 6 | MCI | 84 | M | 5.1 | 18 | *ε3ε4* | 1424 | 2.1 | 0.2 | 2 | 4 | 2 |
| 7 | MCI | 85 | M | 5.0 | 20 | *ε3ε3* | 1480 | 1.0 | 0.2 | 2 | 3 | 2 |
| 8 | MCI | 86 | M | 2.3 | 22 | *ε2ε4* | 1220 | 7.9 | 38.8 | 1 | 5 | 1 |
| 9 | NCI | 74 | M | 7.3 | 22 | *ε3ε3* | n/a[n] | 0.4 | 0 | 0 | 2 | 1 |
| 10 | AD dementia | 81 | M | 4.2 | 15 | *ε3ε4* | 1200 | 1.8 | 7.7 | 2 | 4 | 2 |
| 11 | MCI | 89 | M | 4.5 | 20 | *ε3ε4* | 1300 | 2.4 | 4.5 | 2 | 4 | 2 |
| 12 | AD dementia | 89 | M | 2.6 | 20 | *ε2ε3* | 1230 | 4.1 | 2.8 | 3 | 4 | 2 |
| 13 | MCI | 73 | M | 7.8 | 22 | *ε3ε3* | 1380 | 0.5 | 0 | 0 | 2 | 1 |
| 14 | NCI | 75 | M | 4.8 | 21 | *ε3ε3* | 1470 | 0.2 | 0 | 2 | 1 | 1 |
| 15 | AD dementia | 74 | M | 3.0 | 19 | *ε3ε4* | 1390 | 2.4 | 0.002 | 1 | 2 | 1 |
| 16 | NCI | 86 | M | 6.2 | 16 | *ε3ε4* | 1460 | 4.3 | 0.5 | 2 | 4 | 2 |
| 17 | MCI | 90 | F | 8.5 | 12 | *ε2ε3* | 1102 | 2.7 | 0.8 | 2 | 4 | 2 |
| 18 | NCI | 85 | M | 5.5 | 21 | *ε3ε3* | 1050 | 3.1 | 5.1 | 2 | 4 | 2 |
| 19 | AD dementia | 93 | M | 5.8 | 22 | *ε3ε4* | 1300 | 3.4 | 26.4 | 3 | 5 | 3 |
| 20 | AD dementia | 93 | F | 3.1 | 16 | *ε3ε3* | 1080 | 6.5 | 64.3 | 3 | 5 | 3 |
| 21 | MCI | 87 | F | 1.6 | 18 | *ε2ε3* | 1060 | 2.6 | 0.1 | 3 | 4 | 2 |
| 22 | MCI | 91 | F | 3.3 | 20 | *ε3ε3* | 1025 | 2.0 | 5.0 | 1 | 4 | 1 |
| 23 | AD dementia | 81 | F | 1.8 | 18 | *ε3ε3* | 1120 | 2.1 | 12.6 | 3 | 5 | 3 |
| 24 | AD dementia | 89 | F | 9.0 | 10 | *ε3ε4* | 1140 | 7.0 | 5.1 | 3 | 4 | 2 |
| 25 | AD dementia | 88 | F | 5.2 | 25 | *ε3ε3* | 1250 | 1.0 | 2.6 | 2 | 4 | 2 |
| 26 | AD dementia | 93 | F | 7.3 | 20 | *ε3ε3* | 916 | 4.2 | 34.3 | 3 | 5 | 3 |
| 27 | MCI | 87 | F | 4.1 | 18 | *ε3ε4* | 1107 | 4.4 | 0 | 3 | 3 | 2 |
| 28 | AD dementia | 90 | F | 5.7 | 16 | *ε3ε3* | 1140 | 2.8 | 4.3 | 2 | 4 | 2 |
| 29 | AD dementia | 100 | F | 6.4 | 18 | *ε3ε3* | n/a[n] | 1.3 | 1.8 | 2 | 4 | 2 |

| ID | Diagnosis[a] | Age[b] (yr[c]) | Sex[d] | PMI[e] (hr[f]) | Education (yr[c]) | *APOE*[g] genotype | Br. Wt. (g)[h] | Amyloid-ITG[i] | Tangle-ITG[j] | CERAD[k] | BST[l] | NIA-Reagan[m] |
|---|---|---|---|---|---|---|---|---|---|---|---|---|
| 30 | AD dementia | 92 | F | 4.3 | 16 | *ε3ε3* | 1070 | 3.2 | 3.3 | 2 | 3 | 2 |
| 31 | NCI | 87 | F | 5.0 | 19 | *ε3ε3* | 1350 | 0.5 | 0.8 | 2 | 4 | 2 |
| 32 | NCI | 85 | F | 3.3 | 16 | *ε2ε3* | 1060 | 2.2 | 0.02 | 2 | 3 | 2 |
| 33 | MCI | 85 | F | 4.5 | 16 | *ε3ε3* | 1050 | 3.5 | 0 | 1 | 2 | 1 |
| 34 | MCI | 94 | F | 4.1 | 19 | *ε3ε3* | 1230 | 0.9 | 0.1 | 3 | 5 | 3 |
| 35 | NCI | 80 | F | 5.8 | 20 | *ε3ε3* | 1095 | 2.1 | 0.1 | 2 | 4 | 2 |
| 36 | NCI | 92 | F | 3.1 | 16 | *ε2ε3* | n/a[n] | 7.2 | 11.2 | 2 | 4 | 2 |
| 37 | AD dementia | 79 | F | 11.0 | 20 | *ε3ε4* | 1040 | 4.5 | 3.9 | 3 | 4 | 2 |
| 38 | MCI | 92 | F | 1.8 | 19 | *ε3ε4* | 1150 | 3.4 | 16.8 | 3 | 5 | 3 |
| 39 | MCI | 94 | F | 11.5 | 18 | *ε3ε3* | 1060 | 1.5 | 0.1 | 3 | 4 | 2 |
| 40 | AD dementia | 89 | F | 6.6 | 18 | *ε3ε4* | 1029 | 1.0 | 1.9 | 3 | 4 | 2 |
| 41 | AD dementia | 86 | F | 3.0 | 18 | *ε3ε3* | 1256 | 9.5 | 40.7 | 3 | 5 | 3 |
| 42 | NCI | 97 | F | 8.3 | 11 | *ε3ε3* | 1179 | 0.7 | 0.4 | 1 | 4 | 1 |
| 43 | NCI | 93 | F | 4.8 | 16 | *ε3ε3* | 1110 | 2.8 | 0.2 | 2 | 3 | 2 |
| 44 | NCI | 81 | F | 4.5 | 12 | *ε3ε3* | 1150 | 1.4 | 0.01 | 0 | 1 | 1 |
| 45 | MCI | 92 | F | 7.2 | 16 | *ε3ε4* | 1184 | 2.1 | 1.3 | 1 | 2 | 1 |
| 46 | AD dementia | 90 | M | 6.9 | 16 | *ε3ε4* | 1320 | 8.5 | 13.9 | 3 | 4 | 2 |
| 47 | MCI | 84 | F | 7.4 | 15 | *ε3ε3* | 1211 | 0.9 | 0.6 | 1 | 4 | 1 |
| 48 | NCI | 91 | F | 6.6 | 12 | *ε3ε3* | 1152 | 2.8 | 13.0 | 2 | 4 | 2 |
| 49 | AD dementia | 93 | F | 6.4 | 16 | *ε2ε3* | 940 | 7.3 | 10.3 | 2 | 4 | 2 |
| 50 | NCI | 80 | F | 7.8 | 15 | *ε3ε3* | 930 | 0.7 | 0.1 | 0 | 3 | 1 |
| 51 | MCI | 89 | M | 4.4 | 16 | *ε3ε4* | 1210 | 12.5 | 12.4 | 3 | 4 | 2 |
| 52 | AD dementia | 97 | M | 5.5 | 11 | *ε2ε3* | 1400 | 0.5 | 4.7 | 2 | 4 | 1 |
| 53 | NCI | 86 | F | 3.6 | 17 | *ε3ε3* | 1180 | 2.1 | 0 | 0 | 1 | 1 |
| 54 | AD dementia | 87 | F | 8.3 | 16 | *ε3ε3* | 1056 | 12.6 | 32.5 | 3 | 5 | 3 |
| 55 | MCI | 87 | F | 7.0 | 14 | *ε3ε4* | 1200 | 9.5 | 8.0 | 2 | 5 | 2 |
| 56 | MCI | 85 | F | 7.1 | 13 | *ε3ε4* | 1100 | 15.8 | 16.5 | 2 | 5 | 2 |
| 57 | NCI | 85 | F | 9.4 | 12 | *ε3ε3* | 1140 | 1.1 | 0.3 | 1 | 4 | 1 |
| 58 | NCI | 92 | F | 5.8 | 16 | *ε3ε3* | 1180 | 12.5 | 15.8 | 3 | 5 | 3 |
| 59 | NCI | 85 | M | 4.5 | 12 | *ε3ε3* | 1300 | 8.1 | 0.1 | 2 | 3 | 2 |

| ID | Diagnosis[a] | Age[b] (yr[c]) | Sex[d] | PMI[e] (hr[f]) | Education (yr[c]) | *APOE*[g] genotype | Br. Wt. (g)[h] | Amyloid-ITG[i] | Tangle-ITG[j] | CERAD[k] | BST[l] | NIA-Reagan[m] |
|---|---|---|---|---|---|---|---|---|---|---|---|---|
| 60 | NCI | 93 | F | 7.0 | 8 | *ε3ε3* | 1050 | 9.1 | 4.6 | 3 | 4 | 2 |
| 61 | NCI | 78 | F | 7.5 | 15 | *ε3ε4* | 1500 | 9.0 | 0.7 | 2 | 3 | 2 |

[a]NCI, no cognitive impairment; MCI, mild cognitive impairment; AD dementia, Alzheimer's disease dementia.
[b]Age, age at death.
[c]yr, years.
[d]F, female; M, male.
[e]PMI, post-mortem interval.
[f]hr, hours.
[g]*APOE*, apolipoprotein E gene.
[h]Br. Wt., brain weight; g, grams.
[i]Averaged percent area occupied by Aβ proteins that are identified by molecularly-specific immunohistochemistry and quantified by image analysis in the inferior temporal gyrus (ITG) region [1].
[j]Averaged densities of neuronal neurofibrillary tangles that are identified by molecularly specific immunohistochemistry (AT8 antibody directed against phosphorylated tau at serine 202 and threonine 205) in the inferior temporal gyrus (ITG) region [2].
[k]The consortium to establish a registry for Alzheimer's disease (CERAD) scores for densities of neocortical neuritic plaques [3].
[l]BST, Braak Stages [4].
[m]The modified scores of National Institute on Aging-Reagan (NIA-Reagan) diagnosis of Alzheimer's disease (AD) [5]. The NIA-Reagan diagnosis is based on consensus recommendations for post-mortem diagnosis of AD. The criteria rely on both neurofibrillary tangles (Braak Stages) and neuritic plaques (CERAD).
[n]n/a = not applicable.

**Table S2 Detailed demographic and neuropathological characteristics of individuals for measuring Aβ(42)*56**

| ID | Diagnosis[a] | Age[b] (yr[c]) | Sex[d] | PMI[e] (hr[f]) | Education (yr[c]) | *APOE*[g] genotype | Br. Wt. (g)[h] | Amyloid-ITG[i] | Tangle-ITG[j] | CERAD[k] | BST[l] | NIA-Reagan[m] |
|---|---|---|---|---|---|---|---|---|---|---|---|---|
| 1 | NCI | 101 | F | 7.5 | 16 | *ε3ε3* | 1165 | 11.05 | 4.54 | 2 | 3 | 2 |
| 2 | AD dementia | 94 | M | 4.2 | 15 | *ε3ε3* | 1170 | 3.55 | 3.81 | 3 | 5 | 3 |
| 3 | NCI | 88 | M | 6.2 | 18 | *ε3ε3* | 1170 | 0.94 | 0.01 | 0 | 1 | 1 |
| 4 | NCI | 87 | M | 7.0 | 21 | *ε3ε3* | 1160 | 7.14 | 17.07 | 2 | 5 | 2 |
| 5 | MCI | 86 | M | 2.3 | 22 | *ε2ε4* | 1220 | 7.93 | 38.78 | 1 | 5 | 1 |
| 6 | NCI | 74 | M | 7.25 | 22 | *ε3ε3* | n/a[n] | 0.44 | 0.00 | 0 | 2 | 1 |
| 7 | NCI | 82 | M | 3.0 | 22 | *ε3ε3* | 1280 | 1.00 | 0.43 | 2 | 3 | 2 |
| 8 | AD dementia | 81 | M | 4.2 | 15 | *ε3ε4* | 1200 | 1.83 | 7.70 | 2 | 4 | 2 |
| 9 | MCI | 89 | M | 4.5 | 20 | *ε3ε4* | 1300 | 2.37 | 4.52 | 2 | 4 | 2 |
| 10 | AD dementia | 88 | M | 2.3 | 21 | *ε3ε3* | 1380 | 1.07 | 0.00 | 3 | 3 | 2 |
| 11 | AD dementia | 89 | M | 2.6 | 20 | *ε2ε3* | 1230 | 4.08 | 2.84 | 3 | 4 | 2 |
| 12 | MCI | 73 | M | 7.8 | 22 | *ε3ε3* | 1380 | 0.50 | 0.00 | 0 | 2 | 1 |
| 13 | NCI | 75 | M | 4.8 | 21 | *ε3ε3* | 1470 | 0.18 | 0.00 | 2 | 1 | 1 |
| 14 | AD dementia | 74 | M | 3.0 | 19 | *ε3ε4* | 1390 | 2.43 | 0.00 | 1 | 2 | 1 |
| 15 | NCI | 85 | M | 5.5 | 21 | *ε3ε3* | 1050 | 3.07 | 5.06 | 2 | 4 | 2 |
| 16 | AD dementia | 93 | M | 5.8 | 22 | *ε3ε4* | 1300 | 3.43 | 26.44 | 3 | 5 | 3 |
| 17 | NCI | 84 | M | 6.3 | 18 | *ε3ε3* | 1250 | 1.56 | 0.05 | 3 | 4 | 2 |
| 18 | NCI | 88 | M | 8.2 | 24 | *ε3ε3* | 1088 | 6.44 | 23.82 | 2 | 5 | 2 |
| 19 | AD dementia | 93 | F | 3.1 | 16 | *ε3ε3* | 1080 | 6.48 | 64.25 | 3 | 5 | 3 |
| 20 | MCI | 87 | F | 1.6 | 18 | *ε2ε3* | 1060 | 2.56 | 0.14 | 3 | 4 | 2 |
| 21 | MCI | 90 | F | 12.7 | 19 | *ε3ε4* | 962 | 10.82 | 9.29 | 3 | 5 | 3 |
| 22 | AD dementia | 89 | F | 9.0 | 10 | *ε3ε4* | 1140 | 6.99 | 5.05 | 3 | 4 | 2 |
| 23 | AD dementia | 93 | F | 7.3 | 20 | *ε3ε3* | 916 | 4.17 | 34.33 | 3 | 5 | 3 |
| 24 | AD dementia | 90 | F | 5.7 | 16 | *ε3ε3* | 1140 | 2.85 | 4.27 | 2 | 4 | 2 |
| 25 | AD dementia | 100 | F | 6.4 | 18 | *ε3ε3* | n/a[n] | 1.28 | 1.75 | 2 | 4 | 2 |
| 26 | AD dementia | 92 | F | 4.3 | 16 | *ε3ε3* | 1070 | 3.19 | 3.29 | 2 | 3 | 2 |
| 27 | NCI | 87 | F | 5.0 | 19 | *ε3ε3* | 1350 | 0.50 | 0.82 | 2 | 4 | 2 |
| 28 | NCI | 85 | F | 3.3 | 16 | *ε2ε3* | 1060 | 2.18 | 0.02 | 2 | 3 | 2 |
| 29 | MCI | 95 | F | 18.7 | 17 | *ε3ε3* | 1110 | 1.14 | 0.46 | 2 | 4 | 2 |

| ID | Diagnosis[a] | Age[b] (yr[c]) | Sex[d] | PMI[e] (hr[f]) | Education (yr[c]) | *APOE*[g] genotype | Br. Wt. (g)[h] | Amyloid-ITG[i] | Tangle-ITG[j] | CERAD[k] | BST[l] | NIA-Reagan[m] |
|---|---|---|---|---|---|---|---|---|---|---|---|---|
| 30 | MCI | 94 | F | 4.1 | 19 | *ε3ε3* | 1230 | 0.93 | 0.09 | 3 | 5 | 3 |
| 31 | NCI | 98 | F | 11.8 | 16 | *ε3ε3* | 1140 | 2.76 | 0.01 | 2 | 0 | 1 |
| 32 | NCI | 92 | F | 3.1 | 16 | *ε2ε3* | n/a[n] | 7.16 | 11.19 | 2 | 4 | 2 |
| 33 | MCI | 95 | F | 7.0 | 16 | *ε2ε4* | 1200 | 3.01 | 2.64 | 2 | 4 | 2 |
| 34 | AD dementia | 86 | M | 5.1 | 17 | *ε3ε4* | 1040 | 11.38 | 41.14 | 2 | 5 | 2 |
| 35 | MCI | 92 | F | 1.8 | 19 | *ε3ε4* | 1150 | 3.37 | 16.81 | 3 | 5 | 3 |
| 36 | MCI | 94 | F | 11.5 | 18 | *ε3ε3* | 1060 | 1.54 | 0.13 | 3 | 4 | 2 |
| 37 | AD dementia | 89 | F | 6.6 | 18 | *ε3ε4* | 1029 | 0.98 | 1.89 | 3 | 4 | 2 |
| 38 | MCI | 91 | F | 7.5 | 8 | *ε3ε3* | 1275 | 4.97 | 0.35 | 2 | 4 | 2 |
| 39 | MCI | 90 | M | 0.9 | 24 | *ε3ε4* | 1450 | 5.61 | 12.74 | 2 | 5 | 2 |
| 40 | NCI | 97 | M | 6.9 | 16 | *ε3ε4* | 1147 | 10.96 | 0.02 | 1 | 1 | 1 |
| 41 | NCI | 93 | F | 4.8 | 16 | *ε3ε3* | 1110 | 2.83 | 0.18 | 2 | 3 | 2 |
| 42 | MCI | 91 | M | 6.8 | 9 | *ε3ε3* | 950 | 7.89 | 0.12 | 2 | 3 | 2 |
| 43 | AD dementia | 92 | F | 6.8 | 16 | *ε3ε3* | 1120 | 2.23 | 4.95 | 2 | 4 | 2 |
| 44 | NCI | 81 | F | 4.5 | 12 | *ε3ε3* | 1150 | 1.38 | 0.01 | 0 | 1 | 1 |
| 45 | MCI | 90 | F | 8.9 | 14 | *ε3ε3* | 1100 | 6.75 | 26.95 | 3 | 5 | 3 |
| 46 | AD dementia | 90 | M | 6.9 | 16 | *ε3ε4* | 1320 | 8.48 | 13.90 | 3 | 4 | 2 |
| 47 | AD dementia | 93 | F | 6.4 | 16 | *ε2ε3* | 940 | 7.29 | 10.32 | 2 | 4 | 2 |
| 48 | AD dementia | 97 | M | 5.5 | 11 | *ε2ε3* | 1400 | 0.45 | 4.65 | 2 | 4 | 1 |
| 49 | MCI | 87 | F | 14.5 | 16 | *ε3ε3* | 1050 | 1.97 | 0.00 | 2 | 3 | 2 |
| 50 | NCI | 94 | F | 7.7 | 18 | *ε3ε3* | 1105 | 2.80 | 7.54 | 3 | 4 | 2 |
| 51 | NCI | 84 | M | 4.5 | 16 | *ε3ε3* | 1340 | 6.26 | 0.42 | 2 | 3 | 2 |
| 52 | AD dementia | 93 | F | 4.4 | 18 | *ε3ε3* | 1039 | 10.06 | 6.22 | 3 | 5 | 3 |
| 53 | NCI | 85 | M | 4.5 | 12 | *ε3ε3* | 1300 | 8.01 | 0.16 | 2 | 3 | 2 |
| 54 | MCI | 96 | M | 5.2 | 14 | *ε3ε3* | 1470 | 8.61 | 8.56 | 3 | 2 | 2 |
| 55 | AD dementia | 108 | F | 6.5 | 10 | *ε3ε3* | 950 | 5.41 | 7.88 | 2 | 4 | 2 |
| 56 | MCI | 86 | M | 5.8 | 13 | *ε3ε3* | 1444 | 13.09 | 1.08 | 2 | 5 | 2 |
| 57 | NCI | 87 | F | 5.6 | 22 | *ε3ε3* | 1246 | 12.30 | 2.21 | 3 | 4 | 2 |
| 58 | NCI | 99 | F | 7.6 | 16 | *ε3ε3* | 917 | 8.24 | 2.04 | 2 | 5 | 2 |
| 59 | NCI | 97 | F | 8.3 | 12 | *ε2ε3* | 1164 | 4.57 | 1.35 | 3 | 5 | 3 |

| ID | Diagnosis[a] | Age[b] (yr[c]) | Sex[d] | PMI[e] (hr[f]) | Education (yr[c]) | *APOE*[g] genotype | Br. Wt. (g)[h] | Amyloid-ITG[i] | Tangle-ITG[j] | CERAD[k] | BST[l] | NIA-Reagan[m] |
|---|---|---|---|---|---|---|---|---|---|---|---|---|
| 60 | MCI | 93 | F | 8.2 | 11 | *ε3ε3* | 1128 | 6.30 | 7.55 | 3 | 5 | 3 |
| 61 | NCI | 93 | M | 7.6 | 19 | n/a[n] | 1259 | 10.52 | 1.13 | 2 | 3 | 2 |
| 62 | MCI | 86 | F | 8.3 | 18 | *ε3ε3* | 1366 | 4.55 | 1.80 | 2 | 2 | 1 |

[a]NCI, no cognitive impairment; MCI, mild cognitive impairment; AD dementia, Alzheimer's disease dementia.
[b]Age, age at death.
[c]yr, years.
[d]F, female; M, male.
[e]PMI, post-mortem interval.
[f]hr, hours.
[g]*APOE*, apolipoprotein E gene.
[h]Br. Wt., brain weight; g, grams.
[i]Averaged percent area occupied by Aβ proteins that are identified by molecularly-specific immunohistochemistry and quantified by image analysis in the inferior temporal cortex (ITG) region [1].
[j]Averaged densities of neuronal neurofibrillary tangles that are identified by molecularly specific immunohistochemistry (AT8 antibody directed against phosphorylated tau at serine 202 and threonine 205) in the inferior temporal cortex (ITG) region [2].
[k]The consortium to establish a registry for Alzheimer's disease (CERAD) scores for densities of neocortical neuritic plaques [3].
[l]BST, Braak stages [4].
[m]The modified scores of National Institute on Aging-Reagan (NIA-Reagan) diagnosis of Alzheimer's disease (AD) [5]. The NIA-Reagan diagnosis is based on consensus recommendations for post-mortem diagnosis of AD. The criteria rely on both neurofibrillary tangles (Braak Stages) and neuritic plaques (CERAD).
[n]n/a = not applicable.

**Table S3 Biological samples and reagents used in immunoprecipitation, immunoaffinity purification, and western blotting**

| Figure | ID | Brain Extracts | Capturing Reagent | Matrix[b] | Detecting Reagent |
|---|---|---|---|---|---|
| 1b, lane 1, upper panel | 2 and 5 (Table S4) | 100 µg each, pooled | D8Q7I (3.1µg) | DynaG (50 µL) | Biotin-conjugated mouse monoclonal anti-Aβ(1-x) antibody 82E1 (IBL America, Cat. No.: 10326, RRID: AB_10705565; 1:1,000; 100 ng/mL[c]) / Neutravidin-HRP (Thermo Fisher Scientific, Cat. No.: A2664; 1:5,000) |
| 1b, lane 2, upper panel | 1, 3, 4, and 6 (Table S4) | 50 µg each, pooled | D8Q7I (3.1µg) | DynaG (50 µL) | |
| 1b, lane 3, upper panel | not applicable | not applicable | D8Q7I (3.1µg) | DynaG (50 µL) | |
| 1b, lane 4, upper panel | 59 (Table S1) | 1,400 µg | D8Q7I (3.1µg) | DynaG (50 µL) | |
| 1b, lane 5, upper panel | 14 (Table S1) | 1,400 µg | D8Q7I (3.1µg) | DynaG (50 µL) | |
| 1b, lane 6, upper panel | 34 (Table S1) | 1,400 µg | D8Q7I (3.1µg) | DynaG (50 µL) | |
| 1b, lane 7, upper panel | 39 (Table S1) | 1,400 µg | D8Q7I (3.1µg) | DynaG (50 µL) | |
| 1b, lane 8, upper panel | 46 (Table S1) | 1,400 µg | D8Q7I (3.1µg) | DynaG (50 µL) | |
| 1b, lane 9, upper panel | 19 (Table S1) | 1,400 µg | D8Q7I (3.1µg) | DynaG (50 µL) | |
| 1b, lane 10, upper panel | 14, 19, 34, 39, 46, and 59 (Table S1) | 233 µg each, pooled | RbIgG (3.1µg) | DynaG (50 µL) | |
| 1b, lane 11, upper panel | not applicable | not applicable | not applicable | not applicable | |
| 1b, lane 1, middle panel | 2 and 5 (Table S4) | 25 µg each, pooled[a] | not applicable | not applicable | Rabbit monoclonal anti-NeuN antibody 27-4 (MilliporeSigma, Cat. No.: MABN140; RRID: AB_2571567; 1:5,000) / HRP-conjugated ImmunoPure goat-anti-rabbit IgG (Thermo Fisher Scientific, Cat. No.: 31463; 1:200,000) |
| 1b, lane 2, middle panel | 1, 3, 4, and 6 (Table S4) | 12.5 µg each, pooled[a] | not applicable | not applicable | |
| 1b, lane 3, middle panel | not applicable | not applicable | not applicable | not applicable | |
| 1b, lane 4, middle panel | 59 (Table S1) | 50 µg[a] | not applicable | not applicable | |
| 1b, lane 5, middle panel | 14 (Table S1) | 50 µg[a] | not applicable | not applicable | |
| 1b, lane 6, middle panel | 34 (Table S1) | 50 µg[a] | not applicable | not applicable | |
| 1b, lane 7, middle panel | 39 (Table S1) | 50 µg[a] | not applicable | not applicable | |
| 1b, lane 8, middle panel | 46 (Table S1) | 50 µg[a] | not applicable | not applicable | |
| 1b, lane 9, middle panel | 19 (Table S1) | 50 µg[a] | not applicable | not applicable | |
| 1b, lane 10, middle panel | 14, 19, 34, 39, 46, and 59 (Table S1) | 8.3 µg each, pooled[a] | not applicable | not applicable | |

| | | | | | |
|---|---|---|---|---|---|
| 1b, lane 11, middle panel | not applicable | not applicable | not applicable | not applicable | |
| 1b, lane 1, lower panel | 2 and 5 (Table S4) | 25 µg each, pooled[a] | not applicable | not applicable | Rabbit monoclonal anti-GAPDH antibody 14C10 (Cell Signaling Technology, Cat. No.: 2118S; RRID: AB_561053; 1:5,000; 8.4 ng/mL[c]) / HRP-conjugated ImmunoPure goat-anti-rabbit IgG (Thermo Fisher Scientific, Cat. No.: 31463; 1:200,000) |
| 1b, lane 2, lower panel | 1, 3, 4, and 6 (Table S4) | 12.5 µg each, pooled[a] | not applicable | not applicable | |
| 1b, lane 3, lower panel | not applicable | not applicable | not applicable | not applicable | |
| 1b, lane 4, lower panel | 59 (Table S1) | 50 µg[a] | not applicable | not applicable | |
| 1b, lane 5, lower panel | 14 (Table S1) | 50 µg[a] | not applicable | not applicable | |
| 1b, lane 6, lower panel | 34 (Table S1) | 50 µg[a] | not applicable | not applicable | |
| 1b, lane 7, lower panel | 39 (Table S1) | 50 µg[a] | not applicable | not applicable | |
| 1b, lane 8, lower panel | 46 (Table S1) | 50 µg[a] | not applicable | not applicable | |
| 1b, lane 9, lower panel | 19 (Table S1) | 50 µg[a] | not applicable | not applicable | |
| 1b, lane 10, lower panel | 14, 19, 34, 39, 46, and 59 (Table S1) | 8.3 µg each, pooled[a] | not applicable | not applicable | |
| 1b, lane 11, lower panel | not applicable | not applicable | not applicable | not applicable | |
| 1d, lane 1, upper panel | 2 and 5 (Table S4) | 50 µg each, pooled | D3E10 (3.1µg) | DynaG (50 µL) | Biotin-conjugated mouse monoclonal anti-Aβ(1-x) antibody 82E1 (IBL America, Cat. No.: 10326, RRID: AB_10705565; 1:1,000; 100 ng/mL[c]) / Neutravidin-HRP (Thermo Fisher Scientific, Cat. No.: A2664; 1:5,000) |
| 1d, lane 2, upper panel | 1, 3, 4, and 6 (Table S4) | 50 µg each, pooled | D3E10 (3.1µg) | DynaG (50 µL) | |
| 1d, lane 3, upper panel | not applicable | not applicable | D3E10 (3.1µg) | DynaG (50 µL) | |
| 1d, lane 4, upper panel | 13 (Table S2) | 1,400 µg | D3E10 (3.1µg) | DynaG (50 µL) | |
| 1d, lane 5, upper panel | 28 (Table S2) | 1,400 µg | D3E10 (3.1µg) | DynaG (50 µL) | |
| 1d, lane 6, upper panel | 62 (Table S2) | 1,400 µg | D3E10 (3.1µg) | DynaG (50 µL) | |
| 1d, lane 7, upper panel | 36 (Table S2) | 1,400 µg | D3E10 (3.1µg) | DynaG (50 µL) | |
| 1d, lane 8, upper panel | 34 (Table S2) | 1,400 µg | D3E10 (3.1µg) | DynaG (50 µL) | |
| 1d, lane 9, upper panel | 16 (Table S2) | 1,400 µg | D3E10 (3.1µg) | DynaG (50 µL) | |
| 1d, lane 10, upper panel | 13, 16, 28, 34, 36, and 62 (Table S2) | 233 µg each, pooled | RbIgG (3.1µg) | DynaG (50 µL) | |
| 1d, lane 11, upper panel | not applicable | not applicable | not applicable | not applicable | |

| | | | | | |
|---|---|---|---|---|---|
| 1d, lane 1, middle panel | 2 and 5 (Table S4) | 25 µg each, pooled[a] | not applicable | not applicable | Rabbit monoclonal anti-NeuN antibody 27-4 (MilliporeSigma, Cat. No.: MABN140; RRID: AB_2571567; 1:5,000) / HRP-conjugated ImmunoPure goat-anti-rabbit IgG (Thermo Fisher Scientific, Cat. No.: 31463; 1:200,000) |
| 1d, lane 2, middle panel | 1, 3, 4, and 6 (Table S4) | 12.5 µg each, pooled[a] | not applicable | not applicable | |
| 1d, lane 3, middle panel | not applicable | not applicable | not applicable | not applicable | |
| 1d, lane 4, middle panel | 13 (Table S2) | 50 µg[a] | not applicable | not applicable | |
| 1d, lane 5, middle panel | 28 (Table S2) | 50 µg[a] | not applicable | not applicable | |
| 1d, lane 6, middle panel | 62 (Table S2) | 50 µg[a] | not applicable | not applicable | |
| 1d, lane 7, middle panel | 36 (Table S2) | 50 µg[a] | not applicable | not applicable | |
| 1d, lane 8, middle panel | 34 (Table S2) | 50 µg[a] | not applicable | not applicable | |
| 1d, lane 9, middle panel | 16 (Table S2) | 50 µg[a] | not applicable | not applicable | |
| 1d, lane 10, middle panel | 13, 16, 28, 34, 36, and 62 (Table S2) | 8.3 µg each, pooled[a] | not applicable | not applicable | |
| 1d, lane 11, middle panel | not applicable | not applicable | not applicable | not applicable | |
| 1d, lane 1, lower panel | 2 and 5 (Table S4) | 25 µg each, pooled[a] | not applicable | not applicable | Rabbit monoclonal anti-GAPDH antibody 14C10 (Cell Signaling Technology, Cat. No.: 2118S; RRID: AB_561053; 1:5,000; 8.4 ng/mL[c]) / HRP-conjugated ImmunoPure goat-anti-rabbit IgG (Thermo Fisher Scientific, Cat. No.: 31463; 1:200,000) |
| 1d, lane 2, lower panel | 1, 3, 4, and 6 (Table S4) | 12.5 µg each, pooled[a] | not applicable | not applicable | |
| 1d, lane 3, lower panel | not applicable | not applicable | not applicable | not applicable | |
| 1d, lane 4, lower panel | 13 (Table S2) | 50 µg[a] | not applicable | not applicable | |
| 1d, lane 5, lower panel | 28 (Table S2) | 50 µg[a] | not applicable | not applicable | |
| 1d, lane 6, lower panel | 62 (Table S2) | 50 µg[a] | not applicable | not applicable | |
| 1d, lane 7, lower panel | 36 (Table S2) | 50 µg[a] | not applicable | not applicable | |
| 1d, lane 8, lower panel | 34 (Table S2) | 50 µg[a] | not applicable | not applicable | |
| 1d, lane 9, lower panel | 16 (Table S2) | 50 µg[a] | not applicable | not applicable | |
| 1d, lane 10, lower panel | 13, 16, 28, 34, 36, and 62 (Table S2) | 8.3 µg each, pooled[a] | not applicable | not applicable | |
| 1d, lane 11, lower panel | not applicable | not applicable | not applicable | not applicable | |

| | | | | | |
|---|---|---|---|---|---|
| 1f, lane 1, Tris buffer | 2, 10, 12, 19, 20, 24, 26, 28, 29, and 52 (Table S1) | 150 µg each, pooled | D8Q7I (31µg) | DynaG (500 µL) | Biotin-conjugated rabbit polyclonal anti-oligomer antibody A11 (StressMarq Biosciences, Victoria, BC; Cat. No.: SPC-506D-BI; Lot.: MA333444; RRID: AB_10962958; 1:1,000) / Neutravidin-HRP (Thermo Fisher Scientific, Cat. No.: A2664; 1:5,000) |
| 1f, lane 3, 6M GuHCl | 2, 10, 12, 19, 20, 24, 26, 28, 29, and 52 (Table S1) | 150 µg each, pooled | D8Q7I (31µg) | DynaG (500 µL) | |
| 1g, lane 1, Tris buffer | 2, 8, 11, 16, 19, 22, 23, 24, 25, and 48 (Table S2) | 150 µg each, pooled | D3E10 (31µg) | DynaG (500 µL) | |
| 1g, lane 3, 6M GuHCl | 2, 8, 11, 16, 19, 22, 23, 24, 25, and 48 (Table S2) | 150 µg each, pooled | D3E10 (31µg) | DynaG (500 µL) | |
| 1h, lane "input" | 2, 10, 12, 19, 20, 24, 26, 28, 29, and 52 (Table S1) | 7.5 µg each, pooled | not applicable | not applicable | Biotin-conjugated anti-Aβ(1-x) antibody 82E1 (IBL America, Cat. No.: 10326, RRID: AB_10705565; 1:1,000; 100 ng/mL[c]) / Neutravidin-HRP (Thermo Fisher Scientific, Cat. No.: A2664; 1:5,000) |
| 1h, lane "fractions" | 2, 10, 12, 19, 20, 24, 26, 28, 29, and 52 (Table S1) | 75 µg each, pooled | not applicable | not applicable | |
| S2a, lane 1, Tris buffer | 2, 10, 12, 19, 20, 24, 26, 28, 29, and 52 (Table S1) | 150 µg each, pooled | D8Q7I (31µg) | DynaG (500 µL) | Neutravidin-HRP (Thermo Fisher |

| | | | | | |
|---|---|---|---|---|---|
| S2a, lane 2, 6M GuHCl | 2, 10, 12, 19, 20, 24, 26, 28, 29, and 52 (Table S1) | 150 µg each, pooled | D8Q7I (31µg) | DynaG (500 µL) | Scientific, Cat. No.: A2664; 1:5,000) |
| S2b, lane 1, Tris buffer | 2, 8, 11, 16, 19, 22, 23, 24, 25, and 48 (Table S2) | 150 µg each, pooled | D3E10 (31µg) | DynaG (500 µL) | |
| S2b, lane 2, 6M GuHCl | 2, 8, 11, 16, 19, 22, 23, 24, 25, and 48 (Table S2) | 150 µg each, pooled | D3E10 (31µg) | DynaG (500 µL) | |
| S3, lane "input" | 2, 10, 12, 19, 20, 24, 26, 28, 29, and 52 (Table S1) | 7.5 µg each, pooled | not applicable | not applicable | Neutravidin-HRP (Thermo Fisher Scientific, Cat. No.: A2664; 1:5,000) |
| S3, lane "fractions" | 2, 10, 12, 19, 20, 24, 26, 28, 29, and 52 (Table S1) | 75 µg each, pooled | not applicable | not applicable | |
| S4a, lane 1 | 14 (Table S1) | 50 µg[a] | not applicable | not applicable | Rabbit monoclonal anti-GAPDH antibody 14C10 (Cell Signaling Technology, Cat. No.: 2118S; RRID: AB_561053; 1:5,000; 8.4 ng/mL[c]) / HRP-conjugated ImmunoPure goat-anti-rabbit IgG (Thermo Fisher Scientific, Cat. No.: 31463; 1:200,000) |
| S4a, lane 2 | 13 (Table S1) | 50 µg[a] | not applicable | not applicable | |
| S4a, lane 3 | 5 (Table S1) | 50 µg[a] | not applicable | not applicable | |
| S4a, lane 4 | 8 (Table S1) | 50 µg[a] | not applicable | not applicable | |
| S4a, lane 5 | 9 (Table S1) | 50 µg[a] | not applicable | not applicable | |
| S4a, lane 6 | 40 (Table S1) | 50 µg[a] | not applicable | not applicable | |
| S4a, lane 7 | 52 (Table S1) | 50 µg[a] | not applicable | not applicable | |
| S4a, lane 8 | 31 (Table S1) | 50 µg[a] | not applicable | not applicable | |
| S4a, lane 9 | 10 (Table S1) | 50 µg[a] | not applicable | not applicable | |
| S4a, lane 10 | 7, 8, 9, and 10 (Table S4) | 12.5 µg each, pooled[a] | not applicable | not applicable | |
| S4b, lane 1 | 44 (Table S1) | 50 µg[a] | not applicable | not applicable | |
| S4b, lane 2 | 34 (Table S1) | 50 µg[a] | not applicable | not applicable | |
| S4b, lane 3 | 59 (Table S1) | 50 µg[a] | not applicable | not applicable | |

| | | | | |
|---|---|---|---|---|
| S4b, lane 4 | 46 (Table S1) | 50 µg[a] | not applicable | not applicable |
| S4b, lane 5 | 4 (Table S1) | 50 µg[a] | not applicable | not applicable |
| S4b, lane 6 | 39 (Table S1) | 50 µg[a] | not applicable | not applicable |
| S4b, lane 7 | 32 (Table S1) | 50 µg[a] | not applicable | not applicable |
| S4b, lane 8 | 11 (Table S1) | 50 µg[a] | not applicable | not applicable |
| S4b, lane 9 | 15 (Table S1) | 50 µg[a] | not applicable | not applicable |
| S4b, lane 10 | 7, 8, 9, and 10 (Table S4) | 12.5 µg each, pooled[a] | not applicable | not applicable |
| S4c, lane 1 | 21 (Table S1) | 50 µg[a] | not applicable | not applicable |
| S4c, lane 2 | 28 (Table S1) | 50 µg[a] | not applicable | not applicable |
| S4c, lane 3 | 18 (Table S1) | 50 µg[a] | not applicable | not applicable |
| S4c, lane 4 | 12 (Table S1) | 50 µg[a] | not applicable | not applicable |
| S4c, lane 5 | 38 (Table S1) | 50 µg[a] | not applicable | not applicable |
| S4c, lane 6 | 19 (Table S1) | 50 µg[a] | not applicable | not applicable |
| S4c, lane 7 | 36 (Table S1) | 50 µg[a] | not applicable | not applicable |
| S4c, lane 8 | 20 (Table S1) | 50 µg[a] | not applicable | not applicable |
| S4c, lane 9 | empty | empty | not applicable | not applicable |
| S4c, lane 10 | 7, 8, 9, and 10 (Table S4) | 12.5 µg each, pooled[a] | not applicable | not applicable |
| S4d, lane 1 | 29 (Table S1) | 50 µg[a] | not applicable | not applicable |
| S4d, lane 2 | 43 (Table S1) | 50 µg[a] | not applicable | not applicable |
| S4d, lane 3 | 2 (Table S1) | 50 µg[a] | not applicable | not applicable |
| S4d, lane 4 | not relevant | not applicable | not applicable | not applicable |
| S4d, lane 5 | 26 (Table S1) | 50 µg[a] | not applicable | not applicable |
| S4d, lane 6 | 30 (Table S1) | 50 µg[a] | not applicable | not applicable |
| S4d, lane 7 | 24 (Table S1) | 50 µg[a] | not applicable | not applicable |

| | | | | |
|---|---|---|---|---|
| S4d, lane 8 | 49 (Table S1) | 50 μg[a] | not applicable | not applicable |
| S4d, lane 9 | empty | empty | not applicable | not applicable |
| S4d, lane 10 | 7, 8, 9, and 10 (Table S4) | 12.5 μg each, pooled[a] | not applicable | not applicable |
| S4e, lane 1 | 57 (Table S1) | 50 μg[a] | not applicable | not applicable |
| S4e, lane 2 | 61 (Table S1) | 50 μg[a] | not applicable | not applicable |
| S4e, lane 3 | 7 (Table S1) | 50 μg[a] | not applicable | not applicable |
| S4e, lane 4 | 1 (Table S1) | 50 μg[a] | not applicable | not applicable |
| S4e, lane 5 | 55 (Table S1) | 50 μg[a] | not applicable | not applicable |
| S4e, lane 6 | 25 (Table S1) | 50 μg[a] | not applicable | not applicable |
| S4e, lane 7 | 53 (Table S1) | 50 μg[a] | not applicable | not applicable |
| S4e, lane 8 | 60 (Table S1) | 50 μg[a] | not applicable | not applicable |
| S4e, lane 9 | 6 (Table S1) | 50 μg[a] | not applicable | not applicable |
| S4e, lane 10 | 7, 8, 9, and 10 (Table S4) | 12.5 μg each, pooled[a] | not applicable | not applicable |
| S4f, lane 1 | 45 (Table S1) | 50 μg[a] | not applicable | not applicable |
| S4f, lane 2 | 23 (Table S1) | 50 μg[a] | not applicable | not applicable |
| S4f, lane 3 | 41 (Table S1) | 50 μg[a] | not applicable | not applicable |
| S4f, lane 4 | 16 (Table S1) | 50 μg[a] | not applicable | not applicable |
| S4f, lane 5 | 58 (Table S1) | 50 μg[a] | not applicable | not applicable |
| S4f, lane 6 | 27 (Table S1) | 50 μg[a] | not applicable | not applicable |
| S4f, lane 7 | 51 (Table S1) | 50 μg[a] | not applicable | not applicable |
| S4f, lane 8 | 37 (Table S1) | 50 μg[a] | not applicable | not applicable |
| S4f, lane 9 | 54 (Table S1) | 50 μg[a] | not applicable | not applicable |
| S4f, lane 10 | 7, 8, 9, and 10 (Table S4) | 12.5 μg each, pooled[a] | not applicable | not applicable |

| | | | | | |
|---|---|---|---|---|---|
| S4g, lane 1 | 50 (Table S1) | 50 μg[a] | not applicable | not applicable | |
| S4g, lane 2 | 48 (Table S1) | 50 μg[a] | not applicable | not applicable | |
| S4g, lane 3 | 47 (Table S1) | 50 μg[a] | not applicable | not applicable | |
| S4g, lane 4 | 17 (Table S1) | 50 μg[a] | not applicable | not applicable | |
| S4g, lane 5 | 56 (Table S1) | 50 μg[a] | not applicable | not applicable | |
| S4g, lane 6 | 42 (Table S1) | 50 μg[a] | not applicable | not applicable | |
| S4g, lane 7 | 3 (Table S1) | 50 μg[a] | not applicable | not applicable | |
| S4g, lane 8 | 35 (Table S1) | 50 μg[a] | not applicable | not applicable | |
| S4g, lane 9 | 22 (Table S1) | 50 μg[a] | not applicable | not applicable | |
| S4g, lane 10 | 33 (Table S1) | 50 μg[a] | not applicable | not applicable | |
| S4g, lane 11 | 7, 8, 9, and 10 (Table S4) | 12.5 μg each, pooled[a] | not applicable | not applicable | |
| S5a, lane 1 | 13 (Table S2) | 50 μg[a] | not applicable | not applicable | Rabbit monoclonal anti-GAPDH antibody 14C10 (Cell Signaling Technology, Cat. No.: 2118S; RRID: AB_561053; 1:5,000; 8.4 ng/mL[c]) / HRP-conjugated ImmunoPure goat-anti-rabbit IgG (Thermo Fisher Scientific, Cat. No.: 31463; 1:200,000) |
| S5a, lane 2 | 12 (Table S2) | 50 μg[a] | not applicable | not applicable | |
| S5a, lane 3 | 4 (Table S2) | 50 μg[a] | not applicable | not applicable | |
| S5a, lane 4 | 5 (Table S2) | 50 μg[a] | not applicable | not applicable | |
| S5a, lane 5 | 6 (Table S2) | 50 μg[a] | not applicable | not applicable | |
| S5a, lane 6 | 37 (Table S2) | 50 μg[a] | not applicable | not applicable | |
| S5a, lane 7 | 48 (Table S2) | 50 μg[a] | not applicable | not applicable | |
| S5a, lane 8 | 27 (Table S2) | 50 μg[a] | not applicable | not applicable | |
| S5a, lane 9 | 8 (Table S2) | 50 μg[a] | not applicable | not applicable | |
| S5a, lane 10 | 7, 8, 9, and 10 (Table S4) | 12.5 μg each, pooled[a] | not applicable | not applicable | |
| S5b, lane 1 | 44 (Table S2) | 50 μg[a] | not applicable | not applicable | |
| S5b, lane 2 | 30 (Table S2) | 50 μg[a] | not applicable | not applicable | |
| S5b, lane 3 | 53 (Table S2) | 50 μg[a] | not applicable | not applicable | |

| | | | | |
|---|---|---|---|---|
| S5b, lane 4 | 46 (Table S2) | 50 µg[a] | not applicable | not applicable |
| S5b, lane 5 | 3 (Table S2) | 50 µg[a] | not applicable | not applicable |
| S5b, lane 6 | 36 (Table S2) | 50 µg[a] | not applicable | not applicable |
| S5b, lane 7 | 28 (Table S2) | 50 µg[a] | not applicable | not applicable |
| S5b, lane 8 | 9 (Table S2) | 50 µg[a] | not applicable | not applicable |
| S5b, lane 9 | 14 (Table S2) | 50 µg[a] | not applicable | not applicable |
| S5b, lane 10 | 7, 8, 9, and 10 (Table S4) | 12.5 µg each, pooled[a] | not applicable | not applicable |
| S5c, lane 1 | 20 (Table S2) | 50 µg[a] | not applicable | not applicable |
| S5c, lane 2 | 24 (Table S2) | 50 µg[a] | not applicable | not applicable |
| S5c, lane 3 | 15 (Table S2) | 50 µg[a] | not applicable | not applicable |
| S5c, lane 4 | 11 (Table S2) | 50 µg[a] | not applicable | not applicable |
| S5c, lane 5 | 35 (Table S2) | 50 µg[a] | not applicable | not applicable |
| S5c, lane 6 | 16 (Table S2) | 50 µg[a] | not applicable | not applicable |
| S5c, lane 7 | 32 (Table S2) | 50 µg[a] | not applicable | not applicable |
| S5c, lane 8 | 19 (Table S2) | 50 µg[a] | not applicable | not applicable |
| S5c, lane 9 | empty | empty | not applicable | not applicable |
| S5c, lane 10 | 7, 8, 9, and 10 (Table S4) | 12.5 µg each, pooled[a] | not applicable | not applicable |
| S5d, lane 1 | 25 (Table S2) | 50 µg[a] | not applicable | not applicable |
| S5d, lane 2 | 41 (Table S2) | 50 µg[a] | not applicable | not applicable |
| S5d, lane 3 | 2 (Table S2) | 50 µg[a] | not applicable | not applicable |
| S5d, lane 4 | 33 (Table S2) | 50 µg[a] | not applicable | not applicable |
| S5d, lane 5 | 23 (Table S2) | 50 µg[a] | not applicable | not applicable |
| S5d, lane 6 | 26 (Table S2) | 50 µg[a] | not applicable | not applicable |
| S5d, lane 7 | 22 (Table S2) | 50 µg[a] | not applicable | not applicable |

| | | | | |
|---|---|---|---|---|
| S5d, lane 8 | 47 (Table S2) | 50 µg[a] | not applicable | not applicable |
| S5d, lane 9 | empty | empty | not applicable | not applicable |
| S5d, lane 10 | 7, 8, 9, and 10 (Table S4) | 12.5 µg each, pooled[a] | not applicable | not applicable |
| S5e, lane 1 | 7 (Table S2) | 50 µg[a] | not applicable | not applicable |
| S5e, lane 2 | 58 (Table S2) | 50 µg[a] | not applicable | not applicable |
| S5e, lane 3 | 29 (Table S2) | 50 µg[a] | not applicable | not applicable |
| S5e, lane 4 | 42 (Table S2) | 50 µg[a] | not applicable | not applicable |
| S5e, lane 5 | 54 (Table S2) | 50 µg[a] | not applicable | not applicable |
| S5e, lane 6 | 10 (Table S2) | 50 µg[a] | not applicable | not applicable |
| S5e, lane 7 | 17 (Table S2) | 50 µg[a] | not applicable | not applicable |
| S5e, lane 8 | 61 (Table S2) | 50 µg[a] | not applicable | not applicable |
| S5e, lane 9 | 49 (Table S2) | 50 µg[a] | not applicable | not applicable |
| S5e, lane 10 | 7, 8, 9, and 10 (Table S4) | 12.5 µg each, pooled[a] | not applicable | not applicable |
| S5f, lane 1 | 21 (Table S2) | 50 µg[a] | not applicable | not applicable |
| S5f, lane 2 | 43 (Table S2) | 50 µg[a] | not applicable | not applicable |
| S5f, lane 3 | 52 (Table S2) | 50 µg[a] | not applicable | not applicable |
| S5f, lane 4 | 59 (Table S2) | 50 µg[a] | not applicable | not applicable |
| S5f, lane 5 | 1 (Table S2) | 50 µg[a] | not applicable | not applicable |
| S5f, lane 6 | 38 (Table S2) | 50 µg[a] | not applicable | not applicable |
| S5f, lane 7 | 55 (Table S2) | 50 µg[a] | not applicable | not applicable |
| S5f, lane 8 | 34 (Table S2) | 50 µg[a] | not applicable | not applicable |
| S5f, lane 9 | 51 (Table S2) | 50 µg[a] | not applicable | not applicable |
| S5f, lane 10 | 7, 8, 9, and 10 (Table S4) | 12.5 µg each, pooled[a] | not applicable | not applicable |

| | | | | | |
|---|---|---|---|---|---|
| S5g, lane 1 | 18 (Table S2) | 50 µg[a] | not applicable | not applicable | |
| S5g, lane 2 | 39 (Table S2) | 50 µg[a] | not applicable | not applicable | |
| S5g, lane 3 | 60 (Table S2) | 50 µg[a] | not applicable | not applicable | |
| S5g, lane 4 | 45 (Table S2) | 50 µg[a] | not applicable | not applicable | |
| S5g, lane 5 | 31 (Table S2) | 50 µg[a] | not applicable | not applicable | |
| S5g, lane 6 | 50 (Table S2) | 50 µg[a] | not applicable | not applicable | |
| S5g, lane 7 | 40 (Table S2) | 50 µg[a] | not applicable | not applicable | |
| S5g, lane 8 | 57 (Table S2) | 50 µg[a] | not applicable | not applicable | |
| S5g, lane 9 | 62 (Table S2) | 50 µg[a] | not applicable | not applicable | |
| S5g, lane 10 | 56 (Table S2) | 50 µg[a] | not applicable | not applicable | |
| S5g, lane 11 | 7, 8, 9, and 10 (Table S4) | 12.5 µg each, pooled[a] | not applicable | not applicable | |
| S7a, lane 1 | 14 (Table S1) | 50 µg[a] | not applicable | not applicable | Rabbit monoclonal anti-NeuN antibody 27-4 (MilliporeSigma, Cat. No.: MABN140; RRID: AB_2571567; 1:5,000) / HRP-conjugated ImmunoPure goat-anti-rabbit IgG (Thermo Fisher Scientific, Cat. No.: 31463; 1:200,000) |
| S7a, lane 2 | 13 (Table S1) | 50 µg[a] | not applicable | not applicable | |
| S7a, lane 3 | 5 (Table S1) | 50 µg[a] | not applicable | not applicable | |
| S7a, lane 4 | 8 (Table S1) | 50 µg[a] | not applicable | not applicable | |
| S7a, lane 5 | 9 (Table S1) | 50 µg[a] | not applicable | not applicable | |
| S7a, lane 6 | 40 (Table S1) | 50 µg[a] | not applicable | not applicable | |
| S7a, lane 7 | 52 (Table S1) | 50 µg[a] | not applicable | not applicable | |
| S7a, lane 8 | 31 (Table S1) | 50 µg[a] | not applicable | not applicable | |
| S7a, lane 9 | 10 (Table S1) | 50 µg[a] | not applicable | not applicable | |
| S7a, lane 10 | 7, 8, 9, and 10 (Table S4) | 12.5 µg each, pooled[a] | not applicable | not applicable | |
| S7b, lane 1 | 44 (Table S1) | 50 µg[a] | not applicable | not applicable | |
| S7b, lane 2 | 34 (Table S1) | 50 µg[a] | not applicable | not applicable | |
| S7b, lane 3 | 59 (Table S1) | 50 µg[a] | not applicable | not applicable | |

| | | | | |
|---|---|---|---|---|
| S7b, lane 4 | 46 (Table S1) | 50 µg[a] | not applicable | not applicable |
| S7b, lane 5 | 4 (Table S1) | 50 µg[a] | not applicable | not applicable |
| S7b, lane 6 | 39 (Table S1) | 50 µg[a] | not applicable | not applicable |
| S7b, lane 7 | 32 (Table S1) | 50 µg[a] | not applicable | not applicable |
| S7b, lane 8 | 11 (Table S1) | 50 µg[a] | not applicable | not applicable |
| S7b, lane 9 | 15 (Table S1) | 50 µg[a] | not applicable | not applicable |
| S7b, lane 10 | 7, 8, 9, and 10 (Table S4) | 12.5 µg each, pooled[a] | not applicable | not applicable |
| S7c, lane 1 | 21 (Table S1) | 50 µg[a] | not applicable | not applicable |
| S7c, lane 2 | 28 (Table S1) | 50 µg[a] | not applicable | not applicable |
| S7c, lane 3 | 18 (Table S1) | 50 µg[a] | not applicable | not applicable |
| S7c, lane 4 | 12 (Table S1) | 50 µg[a] | not applicable | not applicable |
| S7c, lane 5 | 38 (Table S1) | 50 µg[a] | not applicable | not applicable |
| S7c, lane 6 | 19 (Table S1) | 50 µg[a] | not applicable | not applicable |
| S7c, lane 7 | 36 (Table S1) | 50 µg[a] | not applicable | not applicable |
| S7c, lane 8 | 20 (Table S1) | 50 µg[a] | not applicable | not applicable |
| S7c, lane 9 | empty | empty | not applicable | not applicable |
| S7c, lane 10 | 7, 8, 9, and 10 (Table S4) | 12.5 µg each, pooled[a] | not applicable | not applicable |
| S7d, lane 1 | 29 (Table S1) | 50 µg[a] | not applicable | not applicable |
| S7d, lane 2 | 43 (Table S1) | 50 µg[a] | not applicable | not applicable |
| S7d, lane 3 | 2 (Table S1) | 50 µg[a] | not applicable | not applicable |
| S7d, lane 4 | not relevant | not applicable | not applicable | not applicable |
| S7d, lane 5 | 26 (Table S1) | 50 µg[a] | not applicable | not applicable |
| S7d, lane 6 | 30 (Table S1) | 50 µg[a] | not applicable | not applicable |
| S7d, lane 7 | 24 (Table S1) | 50 µg[a] | not applicable | not applicable |

| | | | | |
|---|---|---|---|---|
| S7d, lane 8 | 49 (Table S1) | 50 μg[a] | not applicable | not applicable |
| S7d, lane 9 | empty | empty | not applicable | not applicable |
| S7d, lane 10 | 7, 8, 9, and 10 (Table S4) | 12.5 μg each, pooled[a] | not applicable | not applicable |
| S7e, lane 1 | 57 (Table S1) | 50 μg[a] | not applicable | not applicable |
| S7e, lane 2 | 61 (Table S1) | 50 μg[a] | not applicable | not applicable |
| S7e, lane 3 | 7 (Table S1) | 50 μg[a] | not applicable | not applicable |
| S7e, lane 4 | 1 (Table S1) | 50 μg[a] | not applicable | not applicable |
| S7e, lane 5 | 55 (Table S1) | 50 μg[a] | not applicable | not applicable |
| S7e, lane 6 | 25 (Table S1) | 50 μg[a] | not applicable | not applicable |
| S7e, lane 7 | 53 (Table S1) | 50 μg[a] | not applicable | not applicable |
| S7e, lane 8 | 60 (Table S1) | 50 μg[a] | not applicable | not applicable |
| S7e, lane 9 | 6 (Table S1) | 50 μg[a] | not applicable | not applicable |
| S7e, lane 10 | 7, 8, 9, and 10 (Table S4) | 12.5 μg each, pooled[a] | not applicable | not applicable |
| S7f, lane 1 | 45 (Table S1) | 50 μg[a] | not applicable | not applicable |
| S7f, lane 2 | 23 (Table S1) | 50 μg[a] | not applicable | not applicable |
| S7f, lane 3 | 41 (Table S1) | 50 μg[a] | not applicable | not applicable |
| S7f, lane 4 | 16 (Table S1) | 50 μg[a] | not applicable | not applicable |
| S7f, lane 5 | 58 (Table S1) | 50 μg[a] | not applicable | not applicable |
| S7f, lane 6 | 27 (Table S1) | 50 μg[a] | not applicable | not applicable |
| S7f, lane 7 | 51 (Table S1) | 50 μg[a] | not applicable | not applicable |
| S7f, lane 8 | 37 (Table S1) | 50 μg[a] | not applicable | not applicable |
| S7f, lane 9 | 54 (Table S1) | 50 μg[a] | not applicable | not applicable |
| S7f, lane 10 | 7, 8, 9, and 10 (Table S4) | 12.5 μg each, pooled[a] | not applicable | not applicable |

| | | | | | |
|---|---|---|---|---|---|
| S7g, lane 1 | 50 (Table S1) | 50 μg[a] | not applicable | not applicable | |
| S7g, lane 2 | 48 (Table S1) | 50 μg[a] | not applicable | not applicable | |
| S7g, lane 3 | 47 (Table S1) | 50 μg[a] | not applicable | not applicable | |
| S7g, lane 4 | 17 (Table S1) | 50 μg[a] | not applicable | not applicable | |
| S7g, lane 5 | 56 (Table S1) | 50 μg[a] | not applicable | not applicable | |
| S7g, lane 6 | 42 (Table S1) | 50 μg[a] | not applicable | not applicable | |
| S7g, lane 7 | 3 (Table S1) | 50 μg[a] | not applicable | not applicable | |
| S7g, lane 8 | 35 (Table S1) | 50 μg[a] | not applicable | not applicable | |
| S7g, lane 9 | 22 (Table S1) | 50 μg[a] | not applicable | not applicable | |
| S7g, lane 10 | 33 (Table S1) | 50 μg[a] | not applicable | not applicable | |
| S7g, lane 11 | 7, 8, 9, and 10 (Table S4) | 12.5 μg each, pooled[a] | not applicable | not applicable | |
| S8a, lane 1 | 13 (Table S2) | 50 μg[a] | not applicable | not applicable | Rabbit monoclonal anti-NeuN antibody 27-4 (MilliporeSigma, Cat. No.: MABN140; RRID: AB_2571567; 1:5,000) / HRP-conjugated ImmunoPure goat-anti-rabbit IgG (Thermo Fisher Scientific, Cat. No.: 31463; 1:200,000) |
| S8a, lane 2 | 12 (Table S2) | 50 μg[a] | not applicable | not applicable | |
| S8a, lane 3 | 4 (Table S2) | 50 μg[a] | not applicable | not applicable | |
| S8a, lane 4 | 5 (Table S2) | 50 μg[a] | not applicable | not applicable | |
| S8a, lane 5 | 6 (Table S2) | 50 μg[a] | not applicable | not applicable | |
| S8a, lane 6 | 37 (Table S2) | 50 μg[a] | not applicable | not applicable | |
| S8a, lane 7 | 48 (Table S2) | 50 μg[a] | not applicable | not applicable | |
| S8a, lane 8 | 27 (Table S2) | 50 μg[a] | not applicable | not applicable | |
| S8a, lane 9 | 8 (Table S2) | 50 μg[a] | not applicable | not applicable | |
| S8a, lane 10 | 7, 8, 9, and 10 (Table S4) | 12.5 μg each, pooled[a] | not applicable | not applicable | |
| S8b, lane 1 | 44 (Table S2) | 50 μg[a] | not applicable | not applicable | |
| S8b, lane 2 | 30 (Table S2) | 50 μg[a] | not applicable | not applicable | |
| S8b, lane 3 | 53 (Table S2) | 50 μg[a] | not applicable | not applicable | |

| | | | | |
|---|---|---|---|---|
| S8b, lane 4 | 46 (Table S2) | 50 µg[a] | not applicable | not applicable |
| S8b, lane 5 | 3 (Table S2) | 50 µg[a] | not applicable | not applicable |
| S8b, lane 6 | 36 (Table S2) | 50 µg[a] | not applicable | not applicable |
| S8b, lane 7 | 28 (Table S2) | 50 µg[a] | not applicable | not applicable |
| S8b, lane 8 | 9 (Table S2) | 50 µg[a] | not applicable | not applicable |
| S8b, lane 9 | 14 (Table S2) | 50 µg[a] | not applicable | not applicable |
| S8b, lane 10 | 7, 8, 9, and 10 (Table S4) | 12.5 µg each, pooled[a] | not applicable | not applicable |
| S8c, lane 1 | 20 (Table S2) | 50 µg[a] | not applicable | not applicable |
| S8c, lane 2 | 24 (Table S2) | 50 µg[a] | not applicable | not applicable |
| S8c, lane 3 | 15 (Table S2) | 50 µg[a] | not applicable | not applicable |
| S8c, lane 4 | 11 (Table S2) | 50 µg[a] | not applicable | not applicable |
| S8c, lane 5 | 35 (Table S2) | 50 µg[a] | not applicable | not applicable |
| S8c, lane 6 | 16 (Table S2) | 50 µg[a] | not applicable | not applicable |
| S8c, lane 7 | 32 (Table S2) | 50 µg[a] | not applicable | not applicable |
| S8c, lane 8 | 19 (Table S2) | 50 µg[a] | not applicable | not applicable |
| S8c, lane 9 | empty | empty | not applicable | not applicable |
| S8c, lane 10 | 7, 8, 9, and 10 (Table S4) | 12.5 µg each, pooled[a] | not applicable | not applicable |
| S8d, lane 1 | 25 (Table S2) | 50 µg[a] | not applicable | not applicable |
| S8d, lane 2 | 41 (Table S2) | 50 µg[a] | not applicable | not applicable |
| S8d, lane 3 | 2 (Table S2) | 50 µg[a] | not applicable | not applicable |
| S8d, lane 4 | 33 (Table S2) | 50 µg[a] | not applicable | not applicable |
| S8d, lane 5 | 23 (Table S2) | 50 µg[a] | not applicable | not applicable |
| S8d, lane 6 | 26 (Table S2) | 50 µg[a] | not applicable | not applicable |
| S8d, lane 7 | 22 (Table S2) | 50 µg[a] | not applicable | not applicable |

| | | | | |
|---|---|---|---|---|
| S8d, lane 8 | 47 (Table S2) | 50 µg[a] | not applicable | not applicable |
| S8d, lane 9 | empty | empty | not applicable | not applicable |
| S8d, lane 10 | 7, 8, 9, and 10 (Table S4) | 12.5 µg each, pooled[a] | not applicable | not applicable |
| S8e, lane 1 | 7 (Table S2) | 50 µg[a] | not applicable | not applicable |
| S8e, lane 2 | 58 (Table S2) | 50 µg[a] | not applicable | not applicable |
| S8e, lane 3 | 29 (Table S2) | 50 µg[a] | not applicable | not applicable |
| S8e, lane 4 | 42 (Table S2) | 50 µg[a] | not applicable | not applicable |
| S8e, lane 5 | 54 (Table S2) | 50 µg[a] | not applicable | not applicable |
| S8e, lane 6 | 10 (Table S2) | 50 µg[a] | not applicable | not applicable |
| S8e, lane 7 | 17 (Table S2) | 50 µg[a] | not applicable | not applicable |
| S8e, lane 8 | 61 (Table S2) | 50 µg[a] | not applicable | not applicable |
| S8e, lane 9 | 49 (Table S2) | 50 µg[a] | not applicable | not applicable |
| S8e, lane 10 | 7, 8, 9, and 10 (Table S4) | 12.5 µg each, pooled[a] | not applicable | not applicable |
| S8f, lane 1 | 21 (Table S2) | 50 µg[a] | not applicable | not applicable |
| S8f, lane 2 | 43 (Table S2) | 50 µg[a] | not applicable | not applicable |
| S8f, lane 3 | 52 (Table S2) | 50 µg[a] | not applicable | not applicable |
| S8f, lane 4 | 59 (Table S2) | 50 µg[a] | not applicable | not applicable |
| S8f, lane 5 | 1 (Table S2) | 50 µg[a] | not applicable | not applicable |
| S8f, lane 6 | 38 (Table S2) | 50 µg[a] | not applicable | not applicable |
| S8f, lane 7 | 55 (Table S2) | 50 µg[a] | not applicable | not applicable |
| S8f, lane 8 | 34 (Table S2) | 50 µg[a] | not applicable | not applicable |
| S8f, lane 9 | 51 (Table S2) | 50 µg[a] | not applicable | not applicable |
| S8f, lane 10 | 7, 8, 9, and 10 (Table S4) | 12.5 µg each, pooled[a] | not applicable | not applicable |

| | | | | | |
|---|---|---|---|---|---|
| S8g, lane 1 | 18 (Table S2) | 50 µg[a] | not applicable | not applicable | |
| S8g, lane 2 | 39 (Table S2) | 50 µg[a] | not applicable | not applicable | |
| S8g, lane 3 | 60 (Table S2) | 50 µg[a] | not applicable | not applicable | |
| S8g, lane 4 | 45 (Table S2) | 50 µg[a] | not applicable | not applicable | |
| S8g, lane 5 | 31 (Table S2) | 50 µg[a] | not applicable | not applicable | |
| S8g, lane 6 | 50 (Table S2) | 50 µg[a] | not applicable | not applicable | |
| S8g, lane 7 | 40 (Table S2) | 50 µg[a] | not applicable | not applicable | |
| S8g, lane 8 | 57 (Table S2) | 50 µg[a] | not applicable | not applicable | |
| S8g, lane 9 | 62 (Table S2) | 50 µg[a] | not applicable | not applicable | |
| S8g, lane 10 | 56 (Table S2) | 50 µg[a] | not applicable | not applicable | |
| S8g, lane 11 | 7, 8, 9, and 10 (Table S4) | 12.5 µg each, pooled[a] | not applicable | not applicable | |
| S10a, lane 1 | 2 and 5 (Table S4) | 100 µg each, pooled | D8Q7I (3.1µg) | DynaG (50 µL) | Biotin-conjugated mouse monoclonal anti-Aβ(1-x) antibody 82E1 (IBL America, Cat. No.: 10326, RRID: AB_10705565; 1:1,000; 100 ng/mL[c]) / Neutravidin-HRP (Thermo Fisher Scientific, Cat. No.: A2664; 1:5,000) |
| S10a, lane 2 | 1, 3, 4, and 6 (Table S4) | 50 µg each, pooled | D8Q7I (3.1µg) | DynaG (50 µL) | |
| S10a, lane 3 | not applicable | not applicable | D8Q7I (3.1µg) | DynaG (50 µL) | |
| S10a, lane 4 | 14 (Table S1) | 1,400 µg | D8Q7I (3.1µg) | DynaG (50 µL) | |
| S10a, lane 5 | 13 (Table S1) | 1,400 µg | D8Q7I (3.1µg) | DynaG (50 µL) | |
| S10a, lane 6 | 5 (Table S1) | 1,400 µg | D8Q7I (3.1µg) | DynaG (50 µL) | |
| S10a, lane 7 | 8 (Table S1) | 1,400 µg | D8Q7I (3.1µg) | DynaG (50 µL) | |
| S10a, lane 8 | not applicable | not applicable | not applicable | not applicable | |
| S10b, lane 1 | not applicable | not applicable | D8Q7I (3.1µg) | DynaG (50 µL) | |
| S10b, lane 2 | 2 and 5 (Table S4) | 100 µg each, pooled | D8Q7I (3.1µg) | DynaG (50 µL) | |
| S10b, lane 3 | 1, 3, 4, and 6 (Table S4) | 50 µg each, pooled | D8Q7I (3.1µg) | DynaG (50 µL) | |
| S10b, lane 4 | 9 (Table S1) | 1,400 µg | D8Q7I (3.1µg) | DynaG (50 µL) | |

| | | | | |
|---|---|---|---|---|
| S10b, lane 5 | 40 (Table S1) | 1,400 µg | D8Q7I (3.1µg) | DynaG (50 µL) |
| S10b, lane 6 | 52 (Table S1) | 1,400 µg | D8Q7I (3.1µg) | DynaG (50 µL) |
| S10b, lane 7 | 31 (Table S1) | 1,400 µg | D8Q7I (3.1µg) | DynaG (50 µL) |
| S10b, lane 8 | 10 (Table S1) | 1,400 µg | D8Q7I (3.1µg) | DynaG (50 µL) |
| S10b, lane 9 | not applicable | not applicable | not applicable | not applicable |
| S10c, lane 1 | not applicable | not applicable | D8Q7I (3.1µg) | DynaG (50 µL) |
| S10c, lane 2 | 2 and 5 (Table S4) | 100 µg each, pooled | D8Q7I (3.1µg) | DynaG (50 µL) |
| S10c, lane 3 | 1, 3, 4, and 6 (Table S4) | 50 µg each, pooled | D8Q7I (3.1µg) | DynaG (50 µL) |
| S10c, lane 4 | 44 (Table S1) | 1,400 µg | D8Q7I (3.1µg) | DynaG (50 µL) |
| S10c, lane 5 | 34 (Table S1) | 1,400 µg | D8Q7I (3.1µg) | DynaG (50 µL) |
| S10c, lane 6 | 29 (Table S1) | 1,400 µg | D8Q7I (3.1µg) | DynaG (50 µL) |
| S10c, lane 7 | 59 (Table S1) | 1,400 µg | D8Q7I (3.1µg) | DynaG (50 µL) |
| S10c, lane 8 | 46 (Table S1) | 1,400 µg | D8Q7I (3.1µg) | DynaG (50 µL) |
| S10c, lane 9 | not applicable | not applicable | not applicable | not applicable |
| S10d, lane 1 | not applicable | not applicable | D8Q7I (3.1µg) | DynaG (50 µL) |
| S10d, lane 2 | 2 and 5 (Table S4) | 100 µg each, pooled | D8Q7I (3.1µg) | DynaG (50 µL) |
| S10d, lane 3 | 1, 3, 4, and 6 (Table S4) | 50 µg each, pooled | D8Q7I (3.1µg) | DynaG (50 µL) |
| S10d, lane 4 | 4 (Table S1) | 1,400 µg | D8Q7I (3.1µg) | DynaG (50 µL) |
| S10d, lane 5 | 39 (Table S1) | 1,400 µg | D8Q7I (3.1µg) | DynaG (50 µL) |
| S10d, lane 6 | 32 (Table S1) | 1,400 µg | D8Q7I (3.1µg) | DynaG (50 µL) |
| S10d, lane 7 | 11 (Table S1) | 1,400 µg | D8Q7I (3.1µg) | DynaG (50 µL) |
| S10d, lane 8 | 15 (Table S1) | 1,400 µg | D8Q7I (3.1µg) | DynaG (50 µL) |
| S10d, lane 9 | not applicable | not applicable | not applicable | not applicable |
| S10e, lane 1 | not applicable | not applicable | D8Q7I (3.1µg) | DynaG (50 µL) |

| | | | | |
|---|---|---|---|---|
| S10e, lane 2 | 2 and 5 (Table S4) | 100 µg each, pooled | D8Q7I (3.1µg) | DynaG (50 µL) |
| S10e, lane 3 | 1, 3, 4, and 6 (Table S4) | 50 µg each, pooled | D8Q7I (3.1µg) | DynaG (50 µL) |
| S10e, lane 4 | 21 (Table S1) | 1,400 µg | D8Q7I (3.1µg) | DynaG (50 µL) |
| S10e, lane 5 | 28 (Table S1) | 1,400 µg | D8Q7I (3.1µg) | DynaG (50 µL) |
| S10e, lane 6 | 18 (Table S1) | 1,400 µg | D8Q7I (3.1µg) | DynaG (50 µL) |
| S10e, lane 7 | 12 (Table S1) | 1,400 µg | D8Q7I (3.1µg) | DynaG (50 µL) |
| S10e, lane 8 | 2 (Table S1) | 1,400 µg | D8Q7I (3.1µg) | DynaG (50 µL) |
| S10e, lane 9 | not applicable | not applicable | not applicable | not applicable |
| S10f, lane 1 | not applicable | not applicable | D8Q7I (3.1µg) | DynaG (50 µL) |
| S10f, lane 2 | 2 and 5 (Table S4) | 100 µg each, pooled | D8Q7I (3.1µg) | DynaG (50 µL) |
| S10f, lane 3 | 1, 3, 4, and 6 (Table S4) | 50 µg each, pooled | D8Q7I (3.1µg) | DynaG (50 µL) |
| S10f, lane 4 | 43 (Table S1) | 1,400 µg | D8Q7I (3.1µg) | DynaG (50 µL) |
| S10f, lane 5 | 38 (Table S1) | 1,400 µg | D8Q7I (3.1µg) | DynaG (50 µL) |
| S10f, lane 6 | 19 (Table S1) | 1,400 µg | D8Q7I (3.1µg) | DynaG (50 µL) |
| S10f, lane 7 | 36 (Table S1) | 1,400 µg | D8Q7I (3.1µg) | DynaG (50 µL) |
| S10f, lane 8 | 20 (Table S1) | 1,400 µg | D8Q7I (3.1µg) | DynaG (50 µL) |
| S10f, lane 9 | not applicable | not applicable | not applicable | not applicable |
| S10g, lane 1 | not applicable | not applicable | D8Q7I (3.1µg) | DynaG (50 µL) |
| S10g, lane 2 | 2 and 5 (Table S4) | 100 µg each, pooled | D8Q7I (3.1µg) | DynaG (50 µL) |
| S10g, lane 3 | 1, 3, 4, and 6 (Table S4) | 50 µg each, pooled | D8Q7I (3.1µg) | DynaG (50 µL) |
| S10g, lane 4 | 26 (Table S1) | 1,400 µg | D8Q7I (3.1µg) | DynaG (50 µL) |
| S10g, lane 5 | 30 (Table S1) | 1,400 µg | D8Q7I (3.1µg) | DynaG (50 µL) |
| S10g, lane 6 | 24 (Table S1) | 1,400 µg | D8Q7I (3.1µg) | DynaG (50 µL) |

| | | | | |
|---|---|---|---|---|
| S10g, lane 7 | 49 (Table S1) | 1,400 µg | D8Q7I (3.1µg) | DynaG (50 µL) |
| S10g, lane 8 | not applicable | not applicable | not applicable | not applicable |
| S10h, lane 1 | not applicable | not applicable | not applicable | not applicable |
| S10h, lane 2 | 2 and 5 (Table S4) | 100 µg each, pooled | D8Q7I (3.1µg) | DynaG (50 µL) |
| S10h, lane 3 | 1, 3, 4, and 6 (Table S4) | 50 µg each, pooled | D8Q7I (3.1µg) | DynaG (50 µL) |
| S10h, lane 4 | 57 (Table S1) | 1,400 µg | D8Q7I (3.1µg) | DynaG (50 µL) |
| S10h, lane 5 | 61 (Table S1) | 1,400 µg | D8Q7I (3.1µg) | DynaG (50 µL) |
| S10h, lane 6 | 7 (Table S1) | 1,400 µg | D8Q7I (3.1µg) | DynaG (50 µL) |
| S10h, lane 7 | 1 (Table S1) | 1,400 µg | D8Q7I (3.1µg) | DynaG (50 µL) |
| S10h, lane 8 | 55 (Table S1) | 1,400 µg | D8Q7I (3.1µg) | DynaG (50 µL) |
| S10h, lane 9 | 25 (Table S1) | 1,400 µg | D8Q7I (3.1µg) | DynaG (50 µL) |
| S10h, lane 10 | not applicable | not applicable | D8Q7I (3.1µg) | DynaG (50 µL) |
| S10i, lane 1 | not applicable | not applicable | not applicable | not applicable |
| S10i, lane 2 | 2 and 5 (Table S4) | 100 µg each, pooled | D8Q7I (3.1µg) | DynaG (50 µL) |
| S10i, lane 3 | 1, 3, 4, and 6 (Table S4) | 50 µg each, pooled | D8Q7I (3.1µg) | DynaG (50 µL) |
| S10i, lane 4 | not applicable | not applicable | D8Q7I (3.1µg) | DynaG (50 µL) |
| S10i, lane 5 | 53 (Table S1) | 1,400 µg | D8Q7I (3.1µg) | DynaG (50 µL) |
| S10i, lane 6 | 60 (Table S1) | 1,400 µg | D8Q7I (3.1µg) | DynaG (50 µL) |
| S10i, lane 7 | 6 (Table S1) | 1,400 µg | D8Q7I (3.1µg) | DynaG (50 µL) |
| S10i, lane 8 | 45 (Table S1) | 1,400 µg | D8Q7I (3.1µg) | DynaG (50 µL) |
| S10i, lane 9 | 23 (Table S1) | 1,400 µg | D8Q7I (3.1µg) | DynaG (50 µL) |
| S10i, lane 10 | 41 (Table S1) | 1,400 µg | D8Q7I (3.1µg) | DynaG (50 µL) |
| S10j, lane 1 | not applicable | not applicable | not applicable | not applicable |
| S10j, lane 2 | 2 and 5 (Table S4) | 100 µg each, pooled | D8Q7I (3.1µg) | DynaG (50 µL) |

| | | | | |
|---|---|---|---|---|
| S10j, lane 3 | 1, 3, 4, and 6 (Table S4) | 50 µg each, pooled | D8Q7I (3.1µg) | DynaG (50 µL) |
| S10j, lane 4 | not applicable | not applicable | D8Q7I (3.1µg) | DynaG (50 µL) |
| S10j, lane 5 | 16 (Table S1) | 1,400 µg | D8Q7I (3.1µg) | DynaG (50 µL) |
| S10j, lane 6 | 58 (Table S1) | 1,400 µg | D8Q7I (3.1µg) | DynaG (50 µL) |
| S10j, lane 7 | 27 (Table S1) | 1,400 µg | D8Q7I (3.1µg) | DynaG (50 µL) |
| S10j, lane 8 | 51 (Table S1) | 1,400 µg | D8Q7I (3.1µg) | DynaG (50 µL) |
| S10j, lane 9 | 37 (Table S1) | 1,400 µg | D8Q7I (3.1µg) | DynaG (50 µL) |
| S10j, lane 10 | 54 (Table S1) | 1,400 µg | D8Q7I (3.1µg) | DynaG (50 µL) |
| S10k, lane 1 | 2 and 5 (Table S4) | 100 µg each, pooled | D8Q7I (3.1µg) | DynaG (50 µL) |
| S10k, lane 2 | 1, 3, 4, and 6 (Table S4) | 50 µg each, pooled | D8Q7I (3.1µg) | DynaG (50 µL) |
| S10k, lane 3 | not applicable | not applicable | D8Q7I (3.1µg) | DynaG (50 µL) |
| S10k, lane 4 | 50 (Table S1) | 1,400 µg | D8Q7I (3.1µg) | DynaG (50 µL) |
| S10k, lane 5 | 48 (Table S1) | 1,400 µg | D8Q7I (3.1µg) | DynaG (50 µL) |
| S10k, lane 6 | 47 (Table S1) | 1,400 µg | D8Q7I (3.1µg) | DynaG (50 µL) |
| S10k, lane 7 | 17 (Table S1) | 1,400 µg | D8Q7I (3.1µg) | DynaG (50 µL) |
| S10k, lane 8 | 56 (Table S1) | 1,400 µg | D8Q7I (3.1µg) | DynaG (50 µL) |
| S10k, lane 9 | not applicable | not applicable | not applicable | not applicable |
| S10l, lane 1 | 2 and 5 (Table S4) | 100 µg each, pooled | D8Q7I (3.1µg) | DynaG (50 µL) |
| S10l, lane 2 | 1, 3, 4, and 6 (Table S4) | 50 µg each, pooled | D8Q7I (3.1µg) | DynaG (50 µL) |
| S10l, lane 3 | not applicable | not applicable | D8Q7I (3.1µg) | DynaG (50 µL) |
| S10l, lane 4 | 42 (Table S1) | 1,400 µg | D8Q7I (3.1µg) | DynaG (50 µL) |
| S10l, lane 5 | 3 (Table S1) | 1,400 µg | D8Q7I (3.1µg) | DynaG (50 µL) |
| S10l, lane 6 | 35 (Table S1) | 1,400 µg | D8Q7I (3.1µg) | DynaG (50 µL) |

| | | | | | |
|---|---|---|---|---|---|
| S10l, lane 7 | 22 (Table S1) | 1,400 µg | D8Q7I (3.1µg) | DynaG (50 µL) | |
| S10l, lane 8 | 33 (Table S1) | 1,400 µg | D8Q7I (3.1µg) | DynaG (50 µL) | |
| S10l, lane 9 | not applicable | not applicable | not applicable | not applicable | |
| S11a, lane 1 | 2 and 5 (Table S4) | 100 µg each, pooled | D3E10 (3.1µg) | DynaG (50 µL) | Biotin-conjugated mouse monoclonal anti-Aβ(1-x) antibody 82E1 (IBL America, Cat. No.: 10326, RRID: AB_10705565; 1:1,000; 100 ng/mL[c]) / Neutravidin-HRP (Thermo Fisher Scientific, Cat. No.: A2664; 1:5,000) |
| S11a, lane 2 | 1, 3, 4, and 6 (Table S4) | 50 µg each, pooled | D3E10 (3.1µg) | DynaG (50 µL) | |
| S11a, lane 3 | not applicable | not applicable | D3E10 (3.1µg) | DynaG (50 µL) | |
| S11a, lane 4 | 13 (Table S2) | 1,400 µg | D3E10 (3.1µg) | DynaG (50 µL) | |
| S11a, lane 5 | 12 (Table S2) | 1,400 µg | D3E10 (3.1µg) | DynaG (50 µL) | |
| S11a, lane 6 | 4 (Table S2) | 1,400 µg | D3E10 (3.1µg) | DynaG (50 µL) | |
| S11a, lane 7 | 5 (Table S2) | 1,400 µg | D3E10 (3.1µg) | DynaG (50 µL) | |
| S11a, lane 8 | not applicable | not applicable | not applicable | not applicable | |
| S11a, lane 9 | not applicable | not applicable | not applicable | not applicable | |
| S11b, lane 1 | not applicable | not applicable | D3E10 (3.1µg) | DynaG (50 µL) | |
| S11b, lane 2 | 2 and 5 (Table S4) | 100 µg each, pooled | D3E10 (3.1µg) | DynaG (50 µL) | |
| S11b, lane 3 | 1, 3, 4, and 6 (Table S4) | 50 µg each, pooled | D3E10 (3.1µg) | DynaG (50 µL) | |
| S11b, lane 4 | 6 (Table S2) | 1,400 µg | D3E10 (3.1µg) | DynaG (50 µL) | |
| S11b, lane 5 | 37 (Table S2) | 1,400 µg | D3E10 (3.1µg) | DynaG (50 µL) | |
| S11b, lane 6 | 48 (Table S2) | 1,400 µg | D3E10 (3.1µg) | DynaG (50 µL) | |
| S11b, lane 7 | 27 (Table S2) | 1,400 µg | D3E10 (3.1µg) | DynaG (50 µL) | |
| S11b, lane 8 | 8 (Table S2) | 1,400 µg | D3E10 (3.1µg) | DynaG (50 µL) | |
| S11b, lane 9 | not applicable | not applicable | not applicable | not applicable | |
| S11c, lane 1 | not applicable | not applicable | D3E10 (3.1µg) | DynaG (50 µL) | |
| S11c, lane 2 | 2 and 5 (Table S4) | 100 µg each, pooled | D3E10 (3.1µg) | DynaG (50 µL) | |

| | | | | |
|---|---|---|---|---|
| S11c, lane 3 | 1, 3, 4, and 6 (Table S4) | 50 µg each, pooled | D3E10 (3.1µg) | DynaG (50 µL) |
| S11c, lane 4 | 44 (Table S2) | 1,400 µg | D3E10 (3.1µg) | DynaG (50 µL) |
| S11c, lane 5 | 30 (Table S2) | 1,400 µg | D3E10 (3.1µg) | DynaG (50 µL) |
| S11c, lane 6 | 25 (Table S2) | 1,400 µg | D3E10 (3.1µg) | DynaG (50 µL) |
| S11c, lane 7 | 53 (Table S2) | 1,400 µg | D3E10 (3.1µg) | DynaG (50 µL) |
| S11c, lane 8 | 46 (Table S2) | 1,400 µg | D3E10 (3.1µg) | DynaG (50 µL) |
| S11c, lane 9 | not applicable | not applicable | not applicable | not applicable |
| S11d, lane 1 | not applicable | not applicable | D3E10 (3.1µg) | DynaG (50 µL) |
| S11d, lane 2 | 2 and 5 (Table S4) | 100 µg each, pooled | D3E10 (3.1µg) | DynaG (50 µL) |
| S11d, lane 3 | 1, 3, 4, and 6 (Table S4) | 50 µg each, pooled | D3E10 (3.1µg) | DynaG (50 µL) |
| S11d, lane 4 | 3 (Table S2) | 1,400 µg | D3E10 (3.1µg) | DynaG (50 µL) |
| S11d, lane 5 | 36 (Table S2) | 1,400 µg | D3E10 (3.1µg) | DynaG (50 µL) |
| S11d, lane 6 | 28 (Table S2) | 1,400 µg | D3E10 (3.1µg) | DynaG (50 µL) |
| S11d, lane 7 | 9 (Table S2) | 1,400 µg | D3E10 (3.1µg) | DynaG (50 µL) |
| S11d, lane 8 | 14 (Table S2) | 1,400 µg | D3E10 (3.1µg) | DynaG (50 µL) |
| S11d, lane 9 | not applicable | not applicable | not applicable | not applicable |
| S11e, lane 1 | not applicable | not applicable | D3E10 (3.1µg) | DynaG (50 µL) |
| S11e, lane 2 | 2 and 5 (Table S4) | 100 µg each, pooled | D3E10 (3.1µg) | DynaG (50 µL) |
| S11e, lane 3 | 1, 3, 4, and 6 (Table S4) | 50 µg each, pooled | D3E10 (3.1µg) | DynaG (50 µL) |
| S11e, lane 4 | 20 (Table S2) | 1,400 µg | D3E10 (3.1µg) | DynaG (50 µL) |
| S11e, lane 5 | 24 (Table S2) | 1,400 µg | D3E10 (3.1µg) | DynaG (50 µL) |
| S11e, lane 6 | 15 (Table S2) | 1,400 µg | D3E10 (3.1µg) | DynaG (50 µL) |
| S11e, lane 7 | 11 (Table S2) | 1,400 µg | D3E10 (3.1µg) | DynaG (50 µL) |

| | | | | |
|---|---|---|---|---|
| S11e, lane 8 | 2 (Table S2) | 1,400 µg | D3E10 (3.1µg) | DynaG (50 µL) |
| S11e, lane 9 | not applicable | not applicable | not applicable | not applicable |
| S11f, lane 1 | not applicable | not applicable | D3E10 (3.1µg) | DynaG (50 µL) |
| S11f, lane 2 | 2 and 5 (Table S4) | 100 µg each, pooled | D3E10 (3.1µg) | DynaG (50 µL) |
| S11f, lane 3 | 1, 3, 4, and 6 (Table S4) | 50 µg each, pooled | D3E10 (3.1µg) | DynaG (50 µL) |
| S11f, lane 4 | 41 (Table S2) | 1,400 µg | D3E10 (3.1µg) | DynaG (50 µL) |
| S11f, lane 5 | 35 (Table S2) | 1,400 µg | D3E10 (3.1µg) | DynaG (50 µL) |
| S11f, lane 6 | 16 (Table S2) | 1,400 µg | D3E10 (3.1µg) | DynaG (50 µL) |
| S11f, lane 7 | 32 (Table S2) | 1,400 µg | D3E10 (3.1µg) | DynaG (50 µL) |
| S11f, lane 8 | 19 (Table S2) | 1,400 µg | D3E10 (3.1µg) | DynaG (50 µL) |
| S11f, lane 9 | not applicable | not applicable | not applicable | not applicable |
| S11g, lane 1 | not applicable | not applicable | D3E10 (3.1µg) | DynaG (50 µL) |
| S11g, lane 2 | 2 and 5 (Table S4) | 100 µg each, pooled | D3E10 (3.1µg) | DynaG (50 µL) |
| S11g, lane 3 | 1, 3, 4, and 6 (Table S4) | 50 µg each, pooled | D3E10 (3.1µg) | DynaG (50 µL) |
| S11g, lane 4 | 33 (Table S2) | 1,400 µg | D3E10 (3.1µg) | DynaG (50 µL) |
| S11g, lane 5 | 23 (Table S2) | 1,400 µg | D3E10 (3.1µg) | DynaG (50 µL) |
| S11g, lane 6 | 26 (Table S2) | 1,400 µg | D3E10 (3.1µg) | DynaG (50 µL) |
| S11g, lane 7 | 22 (Table S2) | 1,400 µg | D3E10 (3.1µg) | DynaG (50 µL) |
| S11g, lane 8 | 47 (Table S2) | 1,400 µg | D3E10 (3.1µg) | DynaG (50 µL) |
| S11g, lane 9 | not applicable | not applicable | not applicable | not applicable |
| S11h, lane 1 | not applicable | not applicable | not applicable | not applicable |
| S11h, lane 2 | 2 and 5 (Table S4) | 100 µg each, pooled | D3E10 (3.1µg) | DynaG (50 µL) |
| S11h, lane 3 | 1, 3, 4, and 6 (Table S4) | 50 µg each, pooled | D3E10 (3.1µg) | DynaG (50 µL) |

| | | | | |
|---|---|---|---|---|
| S11h, lane 4 | not applicable | not applicable | D3E10 (3.1µg) | DynaG (50 µL) |
| S11h, lane 5 | 7 (Table S2) | 1,400 µg | D3E10 (3.1µg) | DynaG (50 µL) |
| S11h, lane 6 | 58 (Table S2) | 1,400 µg | D3E10 (3.1µg) | DynaG (50 µL) |
| S11h, lane 7 | 29 (Table S2) | 1,400 µg | D3E10 (3.1µg) | DynaG (50 µL) |
| S11h, lane 8 | 42 (Table S2) | 1,400 µg | D3E10 (3.1µg) | DynaG (50 µL) |
| S11h, lane 9 | 54 (Table S2) | 1,400 µg | D3E10 (3.1µg) | DynaG (50 µL) |
| S11h, lane 10 | 10 (Table S2) | 1,400 µg | D3E10 (3.1µg) | DynaG (50 µL) |
| S11i, lane 1 | not applicable | not applicable | not applicable | not applicable |
| S11i, lane 2 | 2 and 5 (Table S4) | 100 µg each, pooled | D3E10 (3.1µg) | DynaG (50 µL) |
| S11i, lane 3 | 1, 3, 4, and 6 (Table S4) | 50 µg each, pooled | D3E10 (3.1µg) | DynaG (50 µL) |
| S11i, lane 4 | not applicable | not applicable | D3E10 (3.1µg) | DynaG (50 µL) |
| S11i, lane 5 | 17 (Table S2) | 1,400 µg | D3E10 (3.1µg) | DynaG (50 µL) |
| S11i, lane 6 | 61 (Table S2) | 1,400 µg | D3E10 (3.1µg) | DynaG (50 µL) |
| S11i, lane 7 | 49 (Table S2) | 1,400 µg | D3E10 (3.1µg) | DynaG (50 µL) |
| S11i, lane 8 | 21 (Table S2) | 1,400 µg | D3E10 (3.1µg) | DynaG (50 µL) |
| S11i, lane 9 | 43 (Table S2) | 1,400 µg | D3E10 (3.1µg) | DynaG (50 µL) |
| S11i, lane 10 | 52 (Table S2) | 1,400 µg | D3E10 (3.1µg) | DynaG (50 µL) |
| S11j, lane 1 | 2 and 5 (Table S4) | 100 µg each, pooled | D3E10 (3.1µg) | DynaG (50 µL) |
| S11j, lane 2 | 1, 3, 4, and 6 (Table S4) | 50 µg each, pooled | D3E10 (3.1µg) | DynaG (50 µL) |
| S11j, lane 3 | not applicable | not applicable | D3E10 (3.1µg) | DynaG (50 µL) |
| S11j, lane 4 | 59 (Table S2) | 1,400 µg | D3E10 (3.1µg) | DynaG (50 µL) |
| S11j, lane 5 | 1 (Table S2) | 1,400 µg | D3E10 (3.1µg) | DynaG (50 µL) |
| S11j, lane 6 | 38 (Table S2) | 1,400 µg | D3E10 (3.1µg) | DynaG (50 µL) |
| S11j, lane 7 | 55 (Table S2) | 1,400 µg | D3E10 (3.1µg) | DynaG (50 µL) |

| | | | | | |
|---|---|---|---|---|---|
| S11j, lane 8 | 34 (Table S2) | 1,400 µg | D3E10 (3.1µg) | DynaG (50 µL) | |
| S11j, lane 9 | not applicable | not applicable | not applicable | not applicable | |
| S11k, lane 1 | 2 and 5 (Table S4) | 100 µg each, pooled | D3E10 (3.1µg) | DynaG (50 µL) | |
| S11k, lane 2 | 1, 3, 4, and 6 (Table S4) | 50 µg each, pooled | D3E10 (3.1µg) | DynaG (50 µL) | |
| S11k, lane 3 | not applicable | not applicable | D3E10 (3.1µg) | DynaG (50 µL) | |
| S11k, lane 4 | 51 (Table S2) | 1,400 µg | D3E10 (3.1µg) | DynaG (50 µL) | |
| S11k, lane 5 | 18 (Table S2) | 1,400 µg | D3E10 (3.1µg) | DynaG (50 µL) | |
| S11k, lane 6 | 39 (Table S2) | 1,400 µg | D3E10 (3.1µg) | DynaG (50 µL) | |
| S11k, lane 7 | 60 (Table S2) | 1,400 µg | D3E10 (3.1µg) | DynaG (50 µL) | |
| S11k, lane 8 | 45 (Table S2) | 1,400 µg | D3E10 (3.1µg) | DynaG (50 µL) | |
| S11k, lane 9 | not applicable | not applicable | not applicable | not applicable | |
| S11l, lane 1 | not applicable | not applicable | not applicable | not applicable | |
| S11l, lane 2 | 2 and 5 (Table S4) | 100 µg each, pooled | D3E10 (3.1µg) | DynaG (50 µL) | |
| S11l, lane 3 | 1, 3, 4, and 6 (Table S4) | 50 µg each, pooled | D3E10 (3.1µg) | DynaG (50 µL) | |
| S11l, lane 4 | not applicable | not applicable | D3E10 (3.1µg) | DynaG (50 µL) | |
| S11l, lane 5 | 31 (Table S2) | 1,400 µg | D3E10 (3.1µg) | DynaG (50 µL) | |
| S11l, lane 6 | 50 (Table S2) | 1,400 µg | D3E10 (3.1µg) | DynaG (50 µL) | |
| S11l, lane 7 | 40 (Table S2) | 1,400 µg | D3E10 (3.1µg) | DynaG (50 µL) | |
| S11l, lane 8 | 57 (Table S2) | 1,400 µg | D3E10 (3.1µg) | DynaG (50 µL) | |
| S11l, lane 9 | 62 (Table S2) | 1,400 µg | D3E10 (3.1µg) | DynaG (50 µL) | |
| S11l, lane 10 | 56 (Table S2) | 1,400 µg | D3E10 (3.1µg) | DynaG (50 µL) | |
| S12a, lane 1 | 2 and 5 (Table S4) | 100 µg each, pooled | D8Q7I (3.1µg) | DynaG (50 µL) | Biotin-conjugated mouse monoclonal anti-Aβ(1-x) |
| S12a, lane 2 | 1, 3, 4, and 6 (Table S4) | 50 µg each, pooled | D8Q7I (3.1µg) | DynaG (50 µL) | |

| | | | | | |
|---|---|---|---|---|---|
| S12a, lane 3 | not applicable | not applicable | D8Q7I (3.1µg) | DynaG (50 µL) | antibody 82E1 (IBL America, Cat. No.: 10326, RRID: AB_10705565; 1:1,000; 100 ng/mL[c]) / Neutravidin-HRP (Thermo Fisher Scientific, Cat. No.: A2664; 1:5,000) |
| S12a, lane 4 | 14 (Table S1) | 1,400 µg | D8Q7I (3.1µg) | DynaG (50 µL) | |
| S12a, lane 5 | 13 (Table S1) | 1,400 µg | D8Q7I (3.1µg) | DynaG (50 µL) | |
| S12a, lane 6 | 5 (Table S1) | 1,400 µg | D8Q7I (3.1µg) | DynaG (50 µL) | |
| S12a, lane 7 | 8 (Table S1) | 1,400 µg | D8Q7I (3.1µg) | DynaG (50 µL) | |
| S12a, lane 8 | not applicable | not applicable | not applicable | not applicable | |
| S12b, lane 1 | not applicable | not applicable | D8Q7I (3.1µg) | DynaG (50 µL) | |
| S12b, lane 2 | 2 and 5 (Table S4) | 100 µg each, pooled | D8Q7I (3.1µg) | DynaG (50 µL) | |
| S12b, lane 3 | 1, 3, 4, and 6 (Table S4) | 50 µg each, pooled | D8Q7I (3.1µg) | DynaG (50 µL) | |
| S12b, lane 4 | 9 (Table S1) | 1,400 µg | D8Q7I (3.1µg) | DynaG (50 µL) | |
| S12b, lane 5 | 40 (Table S1) | 1,400 µg | D8Q7I (3.1µg) | DynaG (50 µL) | |
| S12b, lane 6 | 52 (Table S1) | 1,400 µg | D8Q7I (3.1µg) | DynaG (50 µL) | |
| S12b, lane 7 | 31 (Table S1) | 1,400 µg | D8Q7I (3.1µg) | DynaG (50 µL) | |
| S12b, lane 8 | 10 (Table S1) | 1,400 µg | D8Q7I (3.1µg) | DynaG (50 µL) | |
| S12b, lane 9 | not applicable | not applicable | not applicable | not applicable | |
| S12c, lane 1 | not applicable | not applicable | D8Q7I (3.1µg) | DynaG (50 µL) | |
| S12c, lane 2 | 2 and 5 (Table S4) | 100 µg each, pooled | D8Q7I (3.1µg) | DynaG (50 µL) | |
| S12c, lane 3 | 1, 3, 4, and 6 (Table S4) | 50 µg each, pooled | D8Q7I (3.1µg) | DynaG (50 µL) | |
| S12c, lane 4 | 44 (Table S1) | 1,400 µg | D8Q7I (3.1µg) | DynaG (50 µL) | |
| S12c, lane 5 | 34 (Table S1) | 1,400 µg | D8Q7I (3.1µg) | DynaG (50 µL) | |
| S12c, lane 6 | 29 (Table S1) | 1,400 µg | D8Q7I (3.1µg) | DynaG (50 µL) | |
| S12c, lane 7 | 59 (Table S1) | 1,400 µg | D8Q7I (3.1µg) | DynaG (50 µL) | |
| S12c, lane 8 | 46 (Table S1) | 1,400 µg | D8Q7I (3.1µg) | DynaG (50 µL) | |
| S12c, lane 9 | not applicable | not applicable | not applicable | not applicable | |

| | | | | |
|---|---|---|---|---|
| S12d, lane 1 | not applicable | not applicable | D8Q7I (3.1µg) | DynaG (50 µL) |
| S12d, lane 2 | 2 and 5 (Table S4) | 100 µg each, pooled | D8Q7I (3.1µg) | DynaG (50 µL) |
| S12d, lane 3 | 1, 3, 4, and 6 (Table S4) | 50 µg each, pooled | D8Q7I (3.1µg) | DynaG (50 µL) |
| S12d, lane 4 | 4 (Table S1) | 1,400 µg | D8Q7I (3.1µg) | DynaG (50 µL) |
| S12d, lane 5 | 39 (Table S1) | 1,400 µg | D8Q7I (3.1µg) | DynaG (50 µL) |
| S12d, lane 6 | 32 (Table S1) | 1,400 µg | D8Q7I (3.1µg) | DynaG (50 µL) |
| S12d, lane 7 | 11 (Table S1) | 1,400 µg | D8Q7I (3.1µg) | DynaG (50 µL) |
| S12d, lane 8 | 15 (Table S1) | 1,400 µg | D8Q7I (3.1µg) | DynaG (50 µL) |
| S12d, lane 9 | not applicable | not applicable | not applicable | not applicable |
| S12e, lane 1 | not applicable | not applicable | D8Q7I (3.1µg) | DynaG (50 µL) |
| S12e, lane 2 | 2 and 5 (Table S4) | 100 µg each, pooled | D8Q7I (3.1µg) | DynaG (50 µL) |
| S12e, lane 3 | 1, 3, 4, and 6 (Table S4) | 50 µg each, pooled | D8Q7I (3.1µg) | DynaG (50 µL) |
| S12e, lane 4 | 21 (Table S1) | 1,400 µg | D8Q7I (3.1µg) | DynaG (50 µL) |
| S12e, lane 5 | 28 (Table S1) | 1,400 µg | D8Q7I (3.1µg) | DynaG (50 µL) |
| S12e, lane 6 | 18 (Table S1) | 1,400 µg | D8Q7I (3.1µg) | DynaG (50 µL) |
| S12e, lane 7 | 12 (Table S1) | 1,400 µg | D8Q7I (3.1µg) | DynaG (50 µL) |
| S12e, lane 8 | 2 (Table S1) | 1,400 µg | D8Q7I (3.1µg) | DynaG (50 µL) |
| S12e, lane 9 | not applicable | not applicable | not applicable | not applicable |
| S12f, lane 1 | not applicable | not applicable | D8Q7I (3.1µg) | DynaG (50 µL) |
| S12f, lane 2 | 2 and 5 (Table S4) | 100 µg each, pooled | D8Q7I (3.1µg) | DynaG (50 µL) |
| S12f, lane 3 | 1, 3, 4, and 6 (Table S4) | 50 µg each, pooled | D8Q7I (3.1µg) | DynaG (50 µL) |
| S12f, lane 4 | 43 (Table S1) | 1,400 µg | D8Q7I (3.1µg) | DynaG (50 µL) |
| S12f, lane 5 | 38 (Table S1) | 1,400 µg | D8Q7I (3.1µg) | DynaG (50 µL) |

| | | | | |
|---|---|---|---|---|
| S12f, lane 6 | 19 (Table S1) | 1,400 µg | D8Q7I (3.1µg) | DynaG (50 µL) |
| S12f, lane 7 | 36 (Table S1) | 1,400 µg | D8Q7I (3.1µg) | DynaG (50 µL) |
| S12f, lane 8 | 20 (Table S1) | 1,400 µg | D8Q7I (3.1µg) | DynaG (50 µL) |
| S12f, lane 9 | not applicable | not applicable | not applicable | not applicable |
| S12g, lane 1 | not applicable | not applicable | D8Q7I (3.1µg) | DynaG (50 µL) |
| S12g, lane 2 | 2 and 5 (Table S4) | 100 µg each, pooled | D8Q7I (3.1µg) | DynaG (50 µL) |
| S12g, lane 3 | 1, 3, 4, and 6 (Table S4) | 50 µg each, pooled | D8Q7I (3.1µg) | DynaG (50 µL) |
| S12g, lane 4 | 26 (Table S1) | 1,400 µg | D8Q7I (3.1µg) | DynaG (50 µL) |
| S12g, lane 5 | 30 (Table S1) | 1,400 µg | D8Q7I (3.1µg) | DynaG (50 µL) |
| S12g, lane 6 | 24 (Table S1) | 1,400 µg | D8Q7I (3.1µg) | DynaG (50 µL) |
| S12g, lane 7 | 49 (Table S1) | 1,400 µg | D8Q7I (3.1µg) | DynaG (50 µL) |
| S12g, lane 8 | not applicable | not applicable | not applicable | not applicable |
| S12h, lane 1 | not applicable | not applicable | not applicable | not applicable |
| S12h, lane 2 | 2 and 5 (Table S4) | 100 µg each, pooled | D8Q7I (3.1µg) | DynaG (50 µL) |
| S12h, lane 3 | 1, 3, 4, and 6 (Table S4) | 50 µg each, pooled | D8Q7I (3.1µg) | DynaG (50 µL) |
| S12h, lane 4 | 57 (Table S1) | 1,400 µg | D8Q7I (3.1µg) | DynaG (50 µL) |
| S12h, lane 5 | 61 (Table S1) | 1,400 µg | D8Q7I (3.1µg) | DynaG (50 µL) |
| S12h, lane 6 | 7 (Table S1) | 1,400 µg | D8Q7I (3.1µg) | DynaG (50 µL) |
| S12h, lane 7 | 1 (Table S1) | 1,400 µg | D8Q7I (3.1µg) | DynaG (50 µL) |
| S12h, lane 8 | 55 (Table S1) | 1,400 µg | D8Q7I (3.1µg) | DynaG (50 µL) |
| S12h, lane 9 | 25 (Table S1) | 1,400 µg | D8Q7I (3.1µg) | DynaG (50 µL) |
| S12h, lane 10 | not applicable | not applicable | D8Q7I (3.1µg) | DynaG (50 µL) |
| S12i, lane 1 | not applicable | not applicable | not applicable | not applicable |
| S12i, lane 2 | 2 and 5 (Table S4) | 100 µg each, pooled | D8Q7I (3.1µg) | DynaG (50 µL) |

| | | | | |
|---|---|---|---|---|
| S12i, lane 3 | 1, 3, 4, and 6 (Table S4) | 50 µg each, pooled | D8Q7I (3.1µg) | DynaG (50 µL) |
| S12i, lane 4 | not applicable | not applicable | D8Q7I (3.1µg) | DynaG (50 µL) |
| S12i, lane 5 | 53 (Table S1) | 1,400 µg | D8Q7I (3.1µg) | DynaG (50 µL) |
| S12i, lane 6 | 60 (Table S1) | 1,400 µg | D8Q7I (3.1µg) | DynaG (50 µL) |
| S12i, lane 7 | 6 (Table S1) | 1,400 µg | D8Q7I (3.1µg) | DynaG (50 µL) |
| S12i, lane 8 | 45 (Table S1) | 1,400 µg | D8Q7I (3.1µg) | DynaG (50 µL) |
| S12i, lane 9 | 23 (Table S1) | 1,400 µg | D8Q7I (3.1µg) | DynaG (50 µL) |
| S12i, lane 10 | 41 (Table S1) | 1,400 µg | D8Q7I (3.1µg) | DynaG (50 µL) |
| S12j, lane 1 | not applicable | not applicable | not applicable | not applicable |
| S12j, lane 2 | 2 and 5 (Table S4) | 100 µg each, pooled | D8Q7I (3.1µg) | DynaG (50 µL) |
| S12j, lane 3 | 1, 3, 4, and 6 (Table S4) | 50 µg each, pooled | D8Q7I (3.1µg) | DynaG (50 µL) |
| S12j, lane 4 | not applicable | not applicable | D8Q7I (3.1µg) | DynaG (50 µL) |
| S12j, lane 5 | 16 (Table S1) | 1,400 µg | D8Q7I (3.1µg) | DynaG (50 µL) |
| S12j, lane 6 | 58 (Table S1) | 1,400 µg | D8Q7I (3.1µg) | DynaG (50 µL) |
| S12j, lane 7 | 27 (Table S1) | 1,400 µg | D8Q7I (3.1µg) | DynaG (50 µL) |
| S12j, lane 8 | 51 (Table S1) | 1,400 µg | D8Q7I (3.1µg) | DynaG (50 µL) |
| S12j, lane 9 | 37 (Table S1) | 1,400 µg | D8Q7I (3.1µg) | DynaG (50 µL) |
| S12j, lane 10 | 54 (Table S1) | 1,400 µg | D8Q7I (3.1µg) | DynaG (50 µL) |
| S12k, lane 1 | 2 and 5 (Table S4) | 100 µg each, pooled | D8Q7I (3.1µg) | DynaG (50 µL) |
| S12k, lane 2 | 1, 3, 4, and 6 (Table S4) | 50 µg each, pooled | D8Q7I (3.1µg) | DynaG (50 µL) |
| S12k, lane 3 | not applicable | not applicable | D8Q7I (3.1µg) | DynaG (50 µL) |
| S12k, lane 4 | 50 (Table S1) | 1,400 µg | D8Q7I (3.1µg) | DynaG (50 µL) |
| S12k, lane 5 | 48 (Table S1) | 1,400 µg | D8Q7I (3.1µg) | DynaG (50 µL) |

| | | | | | |
|---|---|---|---|---|---|
| S12k, lane 6 | 47 (Table S1) | 1,400 µg | D8Q7I (3.1µg) | DynaG (50 µL) | |
| S12k, lane 7 | 17 (Table S1) | 1,400 µg | D8Q7I (3.1µg) | DynaG (50 µL) | |
| S12k, lane 8 | 56 (Table S1) | 1,400 µg | D8Q7I (3.1µg) | DynaG (50 µL) | |
| S12k, lane 9 | not applicable | not applicable | not applicable | not applicable | |
| S12l, lane 1 | 2 and 5 (Table S4) | 100 µg each, pooled | D8Q7I (3.1µg) | DynaG (50 µL) | |
| S12l, lane 2 | 1, 3, 4, and 6 (Table S4) | 50 µg each, pooled | D8Q7I (3.1µg) | DynaG (50 µL) | |
| S12l, lane 3 | not applicable | not applicable | D8Q7I (3.1µg) | DynaG (50 µL) | |
| S12l, lane 4 | 42 (Table S1) | 1,400 µg | D8Q7I (3.1µg) | DynaG (50 µL) | |
| S12l, lane 5 | 3 (Table S1) | 1,400 µg | D8Q7I (3.1µg) | DynaG (50 µL) | |
| S12l, lane 6 | 35 (Table S1) | 1,400 µg | D8Q7I (3.1µg) | DynaG (50 µL) | |
| S12l, lane 7 | 22 (Table S1) | 1,400 µg | D8Q7I (3.1µg) | DynaG (50 µL) | |
| S12l, lane 8 | 33 (Table S1) | 1,400 µg | D8Q7I (3.1µg) | DynaG (50 µL) | |
| S12l, lane 9 | not applicable | not applicable | not applicable | not applicable | |
| S13a, lane 1 | 2 and 5 (Table S4) | 100 µg each, pooled | D3E10 (3.1µg) | DynaG (50 µL) | Biotin-conjugated mouse monoclonal anti-Aβ(1-x) antibody 82E1 (IBL America, Cat. No.: 10326, RRID: AB_10705565; 1:1,000; 100 ng/mL[c]) / Neutravidin-HRP (Thermo Fisher Scientific, Cat. No.: A2664; 1:5,000) |
| S13a, lane 2 | 1, 3, 4, and 6 (Table S4) | 50 µg each, pooled | D3E10 (3.1µg) | DynaG (50 µL) | |
| S13a, lane 3 | not applicable | not applicable | D3E10 (3.1µg) | DynaG (50 µL) | |
| S13a, lane 4 | 13 (Table S2) | 1,400 µg | D3E10 (3.1µg) | DynaG (50 µL) | |
| S13a, lane 5 | 12 (Table S2) | 1,400 µg | D3E10 (3.1µg) | DynaG (50 µL) | |
| S13a, lane 6 | 4 (Table S2) | 1,400 µg | D3E10 (3.1µg) | DynaG (50 µL) | |
| S13a, lane 7 | 5 (Table S2) | 1,400 µg | D3E10 (3.1µg) | DynaG (50 µL) | |
| S13a, lane 8 | not applicable | not applicable | not applicable | not applicable | |
| S13a, lane 9 | not applicable | not applicable | not applicable | not applicable | |
| S13b, lane 1 | not applicable | not applicable | D3E10 (3.1µg) | DynaG (50 µL) | |
| S13b, lane 2 | 2 and 5 (Table S4) | 100 µg each, pooled | D3E10 (3.1µg) | DynaG (50 µL) | |

| | | | | |
|---|---|---|---|---|
| S13b, lane 3 | 1, 3, 4, and 6 (Table S4) | 50 µg each, pooled | D3E10 (3.1µg) | DynaG (50 µL) |
| S13b, lane 4 | 6 (Table S2) | 1,400 µg | D3E10 (3.1µg) | DynaG (50 µL) |
| S13b, lane 5 | 37 (Table S2) | 1,400 µg | D3E10 (3.1µg) | DynaG (50 µL) |
| S13b, lane 6 | 48 (Table S2) | 1,400 µg | D3E10 (3.1µg) | DynaG (50 µL) |
| S13b, lane 7 | 27 (Table S2) | 1,400 µg | D3E10 (3.1µg) | DynaG (50 µL) |
| S13b, lane 8 | 8 (Table S2) | 1,400 µg | D3E10 (3.1µg) | DynaG (50 µL) |
| S13b, lane 9 | not applicable | not applicable | not applicable | not applicable |
| S13c, lane 1 | not applicable | not applicable | D3E10 (3.1µg) | DynaG (50 µL) |
| S13c, lane 2 | 2 and 5 (Table S4) | 100 µg each, pooled | D3E10 (3.1µg) | DynaG (50 µL) |
| S13c, lane 3 | 1, 3, 4, and 6 (Table S4) | 50 µg each, pooled | D3E10 (3.1µg) | DynaG (50 µL) |
| S13c, lane 4 | 44 (Table S2) | 1,400 µg | D3E10 (3.1µg) | DynaG (50 µL) |
| S13c, lane 5 | 30 (Table S2) | 1,400 µg | D3E10 (3.1µg) | DynaG (50 µL) |
| S13c, lane 6 | 25 (Table S2) | 1,400 µg | D3E10 (3.1µg) | DynaG (50 µL) |
| S13c, lane 7 | 53 (Table S2) | 1,400 µg | D3E10 (3.1µg) | DynaG (50 µL) |
| S13c, lane 8 | 46 (Table S2) | 1,400 µg | D3E10 (3.1µg) | DynaG (50 µL) |
| S13c, lane 9 | not applicable | not applicable | not applicable | not applicable |
| S13d, lane 1 | not applicable | not applicable | D3E10 (3.1µg) | DynaG (50 µL) |
| S13d, lane 2 | 2 and 5 (Table S4) | 100 µg each, pooled | D3E10 (3.1µg) | DynaG (50 µL) |
| S13d, lane 3 | 1, 3, 4, and 6 (Table S4) | 50 µg each, pooled | D3E10 (3.1µg) | DynaG (50 µL) |
| S13d, lane 4 | 3 (Table S2) | 1,400 µg | D3E10 (3.1µg) | DynaG (50 µL) |
| S13d, lane 5 | 36 (Table S2) | 1,400 µg | D3E10 (3.1µg) | DynaG (50 µL) |
| S13d, lane 6 | 28 (Table S2) | 1,400 µg | D3E10 (3.1µg) | DynaG (50 µL) |
| S13d, lane 7 | 9 (Table S2) | 1,400 µg | D3E10 (3.1µg) | DynaG (50 µL) |

| | | | | |
|---|---|---|---|---|
| S13d, lane 8 | 14 (Table S2) | 1,400 µg | D3E10 (3.1µg) | DynaG (50 µL) |
| S13d, lane 9 | not applicable | not applicable | not applicable | not applicable |
| S13e, lane 1 | not applicable | not applicable | D3E10 (3.1µg) | DynaG (50 µL) |
| S13e, lane 2 | 2 and 5 (Table S4) | 100 µg each, pooled | D3E10 (3.1µg) | DynaG (50 µL) |
| S13e, lane 3 | 1, 3, 4, and 6 (Table S4) | 50 µg each, pooled | D3E10 (3.1µg) | DynaG (50 µL) |
| S13e, lane 4 | 20 (Table S2) | 1,400 µg | D3E10 (3.1µg) | DynaG (50 µL) |
| S13e, lane 5 | 24 (Table S2) | 1,400 µg | D3E10 (3.1µg) | DynaG (50 µL) |
| S13e, lane 6 | 15 (Table S2) | 1,400 µg | D3E10 (3.1µg) | DynaG (50 µL) |
| S13e, lane 7 | 11 (Table S2) | 1,400 µg | D3E10 (3.1µg) | DynaG (50 µL) |
| S13e, lane 8 | 2 (Table S2) | 1,400 µg | D3E10 (3.1µg) | DynaG (50 µL) |
| S13e, lane 9 | not applicable | not applicable | not applicable | not applicable |
| S13f, lane 1 | not applicable | not applicable | D3E10 (3.1µg) | DynaG (50 µL) |
| S13f, lane 2 | 2 and 5 (Table S4) | 100 µg each, pooled | D3E10 (3.1µg) | DynaG (50 µL) |
| S13f, lane 3 | 1, 3, 4, and 6 (Table S4) | 50 µg each, pooled | D3E10 (3.1µg) | DynaG (50 µL) |
| S13f, lane 4 | 41 (Table S2) | 1,400 µg | D3E10 (3.1µg) | DynaG (50 µL) |
| S13f, lane 5 | 35 (Table S2) | 1,400 µg | D3E10 (3.1µg) | DynaG (50 µL) |
| S13f, lane 6 | 16 (Table S2) | 1,400 µg | D3E10 (3.1µg) | DynaG (50 µL) |
| S13f, lane 7 | 32 (Table S2) | 1,400 µg | D3E10 (3.1µg) | DynaG (50 µL) |
| S13f, lane 8 | 19 (Table S2) | 1,400 µg | D3E10 (3.1µg) | DynaG (50 µL) |
| S13f, lane 9 | not applicable | not applicable | not applicable | not applicable |
| S13g, lane 1 | not applicable | not applicable | D3E10 (3.1µg) | DynaG (50 µL) |
| S13g, lane 2 | 2 and 5 (Table S4) | 100 µg each, pooled | D3E10 (3.1µg) | DynaG (50 µL) |
| S13g, lane 3 | 1, 3, 4, and 6 (Table S4) | 50 µg each, pooled | D3E10 (3.1µg) | DynaG (50 µL) |

| | | | | |
|---|---|---|---|---|
| S13g, lane 4 | 33 (Table S2) | 1,400 µg | D3E10 (3.1µg) | DynaG (50 µL) |
| S13g, lane 5 | 23 (Table S2) | 1,400 µg | D3E10 (3.1µg) | DynaG (50 µL) |
| S13g, lane 6 | 26 (Table S2) | 1,400 µg | D3E10 (3.1µg) | DynaG (50 µL) |
| S13g, lane 7 | 22 (Table S2) | 1,400 µg | D3E10 (3.1µg) | DynaG (50 µL) |
| S13g, lane 8 | 47 (Table S2) | 1,400 µg | D3E10 (3.1µg) | DynaG (50 µL) |
| S13g, lane 9 | not applicable | not applicable | not applicable | not applicable |
| S13h, lane 1 | not applicable | not applicable | not applicable | not applicable |
| S13h, lane 2 | 2 and 5 (Table S4) | 100 µg each, pooled | D3E10 (3.1µg) | DynaG (50 µL) |
| S13h, lane 3 | 1, 3, 4, and 6 (Table S4) | 50 µg each, pooled | D3E10 (3.1µg) | DynaG (50 µL) |
| S13h, lane 4 | not applicable | not applicable | D3E10 (3.1µg) | DynaG (50 µL) |
| S13h, lane 5 | 7 (Table S2) | 1,400 µg | D3E10 (3.1µg) | DynaG (50 µL) |
| S13h, lane 6 | 58 (Table S2) | 1,400 µg | D3E10 (3.1µg) | DynaG (50 µL) |
| S13h, lane 7 | 29 (Table S2) | 1,400 µg | D3E10 (3.1µg) | DynaG (50 µL) |
| S13h, lane 8 | 42 (Table S2) | 1,400 µg | D3E10 (3.1µg) | DynaG (50 µL) |
| S13h, lane 9 | 54 (Table S2) | 1,400 µg | D3E10 (3.1µg) | DynaG (50 µL) |
| S13h, lane 10 | 10 (Table S2) | 1,400 µg | D3E10 (3.1µg) | DynaG (50 µL) |
| S13i, lane 1 | not applicable | not applicable | not applicable | not applicable |
| S13i, lane 2 | 2 and 5 (Table S4) | 100 µg each, pooled | D3E10 (3.1µg) | DynaG (50 µL) |
| S13i, lane 3 | 1, 3, 4, and 6 (Table S4) | 50 µg each, pooled | D3E10 (3.1µg) | DynaG (50 µL) |
| S13i, lane 4 | not applicable | not applicable | D3E10 (3.1µg) | DynaG (50 µL) |
| S13i, lane 5 | 17 (Table S2) | 1,400 µg | D3E10 (3.1µg) | DynaG (50 µL) |
| S13i, lane 6 | 61 (Table S2) | 1,400 µg | D3E10 (3.1µg) | DynaG (50 µL) |
| S13i, lane 7 | 49 (Table S2) | 1,400 µg | D3E10 (3.1µg) | DynaG (50 µL) |
| S13i, lane 8 | 21 (Table S2) | 1,400 µg | D3E10 (3.1µg) | DynaG (50 µL) |

| | | | | |
|---|---|---|---|---|
| S13i, lane 9 | 43 (Table S2) | 1,400 µg | D3E10 (3.1µg) | DynaG (50 µL) |
| S13i, lane 10 | 52 (Table S2) | 1,400 µg | D3E10 (3.1µg) | DynaG (50 µL) |
| S13j, lane 1 | 2 and 5 (Table S4) | 100 µg each, pooled | D3E10 (3.1µg) | DynaG (50 µL) |
| S13j, lane 2 | 1, 3, 4, and 6 (Table S4) | 50 µg each, pooled | D3E10 (3.1µg) | DynaG (50 µL) |
| S13j, lane 3 | not applicable | not applicable | D3E10 (3.1µg) | DynaG (50 µL) |
| S13j, lane 4 | 59 (Table S2) | 1,400 µg | D3E10 (3.1µg) | DynaG (50 µL) |
| S13j, lane 5 | 1 (Table S2) | 1,400 µg | D3E10 (3.1µg) | DynaG (50 µL) |
| S13j, lane 6 | 38 (Table S2) | 1,400 µg | D3E10 (3.1µg) | DynaG (50 µL) |
| S13j, lane 7 | 55 (Table S2) | 1,400 µg | D3E10 (3.1µg) | DynaG (50 µL) |
| S13j, lane 8 | 34 (Table S2) | 1,400 µg | D3E10 (3.1µg) | DynaG (50 µL) |
| S13j, lane 9 | not applicable | not applicable | not applicable | not applicable |
| S13k, lane 1 | 2 and 5 (Table S4) | 100 µg each, pooled | D3E10 (3.1µg) | DynaG (50 µL) |
| S13k, lane 2 | 1, 3, 4, and 6 (Table S4) | 50 µg each, pooled | D3E10 (3.1µg) | DynaG (50 µL) |
| S13k, lane 3 | not applicable | not applicable | D3E10 (3.1µg) | DynaG (50 µL) |
| S13k, lane 4 | 51 (Table S2) | 1,400 µg | D3E10 (3.1µg) | DynaG (50 µL) |
| S13k, lane 5 | 18 (Table S2) | 1,400 µg | D3E10 (3.1µg) | DynaG (50 µL) |
| S13k, lane 6 | 39 (Table S2) | 1,400 µg | D3E10 (3.1µg) | DynaG (50 µL) |
| S13k, lane 7 | 60 (Table S2) | 1,400 µg | D3E10 (3.1µg) | DynaG (50 µL) |
| S13k, lane 8 | 45 (Table S2) | 1,400 µg | D3E10 (3.1µg) | DynaG (50 µL) |
| S13k, lane 9 | not applicable | not applicable | not applicable | not applicable |
| S13l, lane 1 | not applicable | not applicable | not applicable | not applicable |
| S13l, lane 2 | 2 and 5 (Table S4) | 100 µg each, pooled | D3E10 (3.1µg) | DynaG (50 µL) |
| S13l, lane 3 | 1, 3, 4, and 6 (Table S4) | 50 µg each, pooled | D3E10 (3.1µg) | DynaG (50 µL) |

| | | | | |
|---|---|---|---|---|
| S13l, lane 4 | not applicable | not applicable | D3E10 (3.1µg) | DynaG (50 µL) |
| S13l, lane 5 | 31 (Table S2) | 1,400 µg | D3E10 (3.1µg) | DynaG (50 µL) |
| S13l, lane 6 | 50 (Table S2) | 1,400 µg | D3E10 (3.1µg) | DynaG (50 µL) |
| S13l, lane 7 | 40 (Table S2) | 1,400 µg | D3E10 (3.1µg) | DynaG (50 µL) |
| S13l, lane 8 | 57 (Table S2) | 1,400 µg | D3E10 (3.1µg) | DynaG (50 µL) |
| S13l, lane 9 | 62 (Table S2) | 1,400 µg | D3E10 (3.1µg) | DynaG (50 µL) |
| S13l, lane 10 | 56 (Table S2) | 1,400 µg | D3E10 (3.1µg) | DynaG (50 µL) |

[a]These are detergent-containing brain extracts prepared from pellets of detergent-free aqueous brain homogenates (see Materials and Methods for details). [b]The volumes listed in this column are the volumes of beads in slurry. The net volumes of beads are approximately 10% (DynaG) of the reported volumes. DynaG, Dynabeads Protein G. [c]The final concentration of the primary antibody is shown.

**Table S4 Mice used in this study**

| ID | Line | Tg[a] | Background | Age (Mo.)[b] | Gender[c] |
|---|---|---|---|---|---|
| 1 | APP/TTA | +/+ | B6 | 10.0 | M |
| 2 | APP/TTA | +/- | B6 | 10.6 | F |
| 3 | APP/TTA | +/+ | B6 | 10.6 | F |
| 4 | APP/TTA | +/+ | B6 | 10.7 | M |
| 5 | APP/TTA | +/- | B6 | 10.7 | M |
| 6 | APP/TTA | +/+ | B6 | 10.7 | M |
| 7 | Tg2576 | + | B6S | 4.0 | M |
| 8 | Tg2576 | + | B6S | 4.0 | M |
| 9 | Tg2576 | + | B6S | 5.3 | F |
| 10 | Tg2576 | + | B6S | 5.3 | F |

[a]+, mice that express the 695-amino-acid human amyloid precursor protein isoform with the Swedish (K670N, M671L) mutation; +/+, mice that express both a tetracycline transactivator and the 695-amino-acid mouse amyloid precursor protein isoform with a humanized Aβ region plus the Swedish (K670N, M671L) and Indiana (V717F) mutations; +/-, mice that express only a tetracycline transactivator but no amyloid precursor protein transgene. [b]Mo., months. [c]F, female; M, male.

**Table S5 Mediation analysis of the contribution of ITG ptau 202/205 NFT to the effects of Aβ(40)*56 on cognitive and memory impairment**

| | ACME[a] | | ADE[b] | | Total Effect[c] | | Proportion Mediated[d] | |
|---|---|---|---|---|---|---|---|---|
| | Estimate (95% CI[e]) | *P* | Estimate (95% CI[e]) | *P* | Estimate (95% CI[e]) | *P* | Estimate (95% CI[e]) | *P* |
| MMSE[f] | -0.477 (-1.06, -0.04) | 0.03 | -1.08 (-2.14, -0.048) | 0.04 | -1.56 (-2.64, -0.474) | 0.004 | 0.298 (0.0245, 0.904) | 0.03 |
| Ep[g] | -0.0888 (-0.185, -0.0104) | 0.02 | -0.216 (-0.364, -0.0727) | 0.002 | -0.305 (-0.464, -0.147) | <0.001 | 0.286 (0.0381, 0.625) | 0.02 |
| Va/po[h] | -0.0295 -0.0892, 0.0116) | 0.18 | -0.107 (-0.25, 0.034) | 0.14 | -0.137 (-0.276, 0.00347) | 0.06 | 0.193 (-0.41, 1.49) | 0.23 |
| Ps[i] | -0.0629 (-0.141, -0.00492) | 0.03 | -0.0858 (-0.229, 0.0545) | 0.23 | -0.149 (-0.295, -0.00157) | 0.05 | 0.408 (-0.0995, 2.03) | 0.07 |
| Sm[j] | -0.0573 (-0.137, -0.00122) | 0.04 | -0.106 (-0.267, 0.0535) | 0.19 | -0.163 (-0.324, -0.000557) | 0.05 | 0.332 (-0.137, 1.82) | 0.08 |
| Wm[k] | -0.0648 (-0.145, -0.00519) | 0.03 | -0.044 (-0.191, 0.0996) | 0.54 | -0.109 (-0.259, 0.0416) | 0.16 | 0.522 (-2.7, 4.27) | 0.17 |
| Gcf[l] | -0.0704 (-0.15, -0.00705) | 0.02 | -0.135 (-0.266, -0.00728) | 0.04 | -0.205 (-0.342, -0.0682) | 0.002 | 0.336 (0.037, 0.908) | 0.03 |

[a]ACME, average causal mediation effect that measures the indirect effect of the predictor (Aβ(40)*56) on the cognitive outcome through the mediator (ITG ptau 202/205 NFT). [b]ADE, average direct effect that measures the direct effect of the predictor on the cognitive outcome, not through the mediator. [c]Total Effect, sum of ACME and ADE. [d]Proportion Mediated, percentage of total effect mediated through the mediator. [e]CI, confidence interval. [f]MMSE, mini-mental state examination. [g]Em, episodic memory. [h]Va/po, visuospatial ability/perceptual orientation. [i]Ps, perceptual speed. [j]Sm, semantic memory. [k]Wm, working memory. [l]Gcf, global cognitive function. ITG ptau 202/205 NFT, immunohistologically determined neurofibrillary tangle loads in the inferior temporal gyrus using the antibody AT8 that recognizes phosphorylated tau at serine 202 and threonine 205.

**Table S6 Mediation analysis of the contribution of ITG ptau 202/205 NFT to the effects of Aβ(42)*56 on cognitive and memory impairment**

| | ACME[a] | | ADE[b] | | Total Effect[c] | | Proportion Mediated[d] | |
|---|---|---|---|---|---|---|---|---|
| | Estimate (95% CI[e]) | *P* | Estimate (95% CI[e]) | *P* | Estimate (95% CI[e]) | *P* | Estimate (95% CI[e]) | *P* |
| MMSE[f] | -0.381 (-0.911, -0.00649) | 0.05 | -0.931 (-1.98, 0.0928) | 0.08 | -1.31 (-2.3, -0.303) | 0.01 | 0.281 (-0.00747, 1.16) | 0.05 |
| Ep[g] | -0.0788 (-0.171, -0.012) | 0.01 | -0.124 (-0.291, 0.0391) | 0.14 | -0.203 (-0.366, -0.0391) | 0.01 | 0.381 (0.047, 1.53) | 0.03 |
| Va/po[h] | -0.0354 (-0.101, 0.0143) | 0.17 | -0.0929 (-0.237, 0.0481) | 0.19 | -0.128 (-0.261, 0.00787) | 0.06 | 0.251 (-0.728, 2.05) | 0.23 |
| Ps[i] | -0.0532 (-0.128, -0.000186) | 0.05 | -0.119 (-0.268, 0.0259) | 0.12 | -0.173 (-0.312, -0.0297) | 0.01 | 0.298 (-0.0177, 1.36) | 0.06 |
| Sm[j] | -0.0351 (-0.105, 0.0175) | 0.20 | -0.183 (-0.337, -0.0332) | 0.01 | -0.218 (-0.359, -0.0742) | <0.001 | 0.153 (-0.0914, 0.638) | 0.20 |
| Wm[k] | -0.0506 (-0.123, 0.00123) | 0.06 | -0.102 (-0.248, 0.0414) | 0.16 | -0.152 (-0.289, -0.0124) | 0.03 | 0.316 (-0.0875, 1.7) | 0.09 |
| Gcf[l] | -0.0621 (-0.137, -0.00767) | 0.02 | -0.123 (-0.262, 0.0132) | 0.08 | -0.185 (-0.319, -0.049) | 0.01 | 0.329 (0.0378, 1.16) | 0.02 |

[a]ACME, average causal mediation effect that measures the indirect effect of the predictor (Aβ(42)*56) on the cognitive outcome through the mediator (ITG ptau 202/205 NFT). [b]ADE, average direct effect that measures the direct effect of the predictor on the cognitive outcome, not through the mediator. [c]Total Effect, sum of ACME and ADE. [d]Proportion Mediated, percentage of total effect mediated through the mediator. [e]CI, confidence interval. [f]MMSE, mini-mental state examination. [g]Em, episodic memory. [h]Va/po, visuospatial ability/perceptual orientation. [i]Ps, perceptual speed. [j]Sm, semantic memory. [k]Wm, working memory. [l]Gcf, global cognitive function. ITG ptau 202/205 NFT, immunohistologically determined neurofibrillary tangle loads in the inferior temporal gyrus using the antibody AT8 that recognizes phosphorylated tau at serine 202 and threonine 205.

**Table S7 Mediation analysis of the contribution of ITG Aβ Plaques to the effects of Aβ(40)*56 on cognitive and memory impairment**

| | ACME[a] | | ADE[b] | | Total Effect[c] | | Proportion Mediated[d] | |
|---|---|---|---|---|---|---|---|---|
| | Estimate (95% CI[e]) | *P* | Estimate (95% CI[e]) | *P* | Estimate (95% CI[e]) | *P* | Estimate (95% CI[e]) | *P* |
| MMSE[f] | -0.281 (-0.762, 0.0366) | 0.10 | -1.28 (-2.39, -0.201) | 0.02 | -1.56 (-2.64, -0.473) | 0.003 | 0.168 (-0.0268, 0.672) | 0.11 |
| Ep[g] | -0.0303 (-0.0968, 0.0165) | 0.23 | -0.275 (-0.441, -0.115) | <0.001 | -0.305 (-0.465, -0.143) | <0.001 | 0.0894 (-0.0552, 0.348) | 0.23 |
| Va/po[h] | -0.00988 (-0.0585, 0.0316) | 0.62 | -0.127 (-0.272, 0.013) | 0.09 | -0.137 (-0.275, 0.00135) | 0.05 | 0.0564 (-0.51, 0.939) | 0.65 |
| Ps[i] | -0.0118 (-0.0651, 0.0341) | 0.60 | -0.137 (-0.291, 0.0122) | 0.08 | -0.149 (-0.296, -0.00263) | 0.05 | 0.064 (-0.487, 0.897) | 0.62 |
| Sm[j] | -0.0315 (-0.0981, 0.0133) | 0.20 | -0.131 (-0.298, 0.0291) | 0.12 | -0.163 (-0.324, -0.000566) | 0.05 | 0.17 (-0.247, 1.38) | 0.23 |
| Wm[k] | -0.0146 (-0.0704, 0.0318) | 0.53 | -0.0941 (-0.252, 0.0583) | 0.24 | -0.109 (-0.259, 0.0416) | 0.16 | 0.0929 (-1.37, 1.63) | 0.62 |
| Gcf[l] | -0.0254 (-0.0824, 0.0154) | 0.25 | -0.18 (-0.324, -0.041) | 0.01 | -0.205 (-0.344, -0.0654) | 0.002 | 0.112 (-0.082, 0.521) | 0.25 |

[a]ACME, average causal mediation effect that measures the indirect effect of the predictor (Aβ(40)*56) on the cognitive outcome through the mediator (ITG Aβ Plaques). [b]ADE, average direct effect that measures the direct effect of the predictor on the cognitive outcome, not through the mediator. [c]Total Effect, sum of ACME and ADE. [d]Proportion Mediated, percentage of total effect mediated through the mediator. [e]CI, confidence interval. [f]MMSE, mini-mental state examination. [g]Em, episodic memory. [h]Va/po, visuospatial ability/perceptual orientation. [i]Ps, perceptual speed. [j]Sm, semantic memory. [k]Wm, working memory. [l]Gcf, global cognitive function. ITG Aβ Plaques, immunohistologically determined β-amyloid plaque loads in the inferior temporal gyrus.

**Table S8 Mediation analysis of the contribution of ITG Aβ Plaques to the effects of Aβ(42)*56 on cognitive and memory impairment**

| | ACME[a] | | ADE[b] | | Total Effect[c] | | Proportion Mediated[d] | |
|---|---|---|---|---|---|---|---|---|
| | Estimate (95% CI[e]) | *P* | Estimate (95% CI[e]) | *P* | Estimate (95% CI[e]) | *P* | Estimate (95% CI[e]) | *P* |
| MMSE[f] | 0.00644 (-0.253, 0.281) | 0.96 | -1.32 (-2.34, -0.31) | 0.01 | -1.31 (-2.3, -0.317) | 0.01 | -0.00207 (-0.323, 0.244) | 0.96 |
| Ep[g] | -0.0106 (-0.0604, 0.029) | 0.59 | -0.192 (-0.359, -0.0281) | 0.02 | -0.203 (-0.362, -0.0404) | 0.01 | 0.0382 (-0.199, 0.427) | 0.60 |
| Va/po[h] | 0.00426 (-0.0292, 0.0433) | 0.81 | -0.133 (-0.271, 0.00358) | 0.06 | -0.128 (-0.262, 0.00715) | 0.06 | -0.016 (-0.697, 0.532) | 0.83 |
| Ps[i] | 0.00729 (-0.0267, 0.0502) | 0.69 | -0.18 (-0.325, -0.0371) | 0.01 | -0.172 (-0.313, -0.0297) | 0.02 | -0.0268 (-0.546, 0.211) | 0.70 |
| Sm[j] | -0.00177 (-0.0405, 0.0367) | 0.92 | -0.217 (-0.364, -0.072) | 0.003 | -0.218 (-0.36, -0.0757) | 0.002 | 0.00489 (-0.217, 0.22) | 0.92 |
| Wm[k] | -0.00169 (-0.0393, 0.0357) | 0.92 | -0.151 (-0.294, -0.0101) | 0.03 | -0.152 (-0.29, -0.0137) | 0.03 | 0.00653 (-0.436, 0.417) | 0.92 |
| Gcf[l] | -0.00121 (-0.0373, 0.0352) | 0.94 | -0.183 (-0.322, -0.0473) | 0.01 | -0.185 (-0.318, -0.0505) | 0.004 | 0.00399 (-0.267, 0.251) | 0.94 |

[a]ACME, average causal mediation effect that measures the indirect effect of the predictor (Aβ(42)*56) on the cognitive outcome through the mediator (ITG Aβ Plaques). [b]ADE, average direct effect that measures the direct effect of the predictor on the cognitive outcome, not through the mediator. [c]Total Effect, sum of ACME and ADE. [d]Proportion Mediated, percentage of total effect mediated through the mediator. [e]CI, confidence interval. [f]MMSE, mini-mental state examination. [g]Em, episodic memory. [h]Va/po, visuospatial ability/perceptual orientation. [i]Ps, perceptual speed. [j]Sm, semantic memory. [k]Wm, working memory. [l]Gcf, global cognitive function. ITG Aβ Plaques, immunohistologically determined β-amyloid plaque loads in the inferior temporal gyrus.
